# A selective autophagy pathway for phase separated endocytic protein deposits

**DOI:** 10.1101/2020.05.26.116368

**Authors:** Florian Wilfling, Chia-Wei Lee, Philipp Erdmann, Yumei Zheng, Stefan Jentsch, Boris Pfander, Brenda A. Schulman, Wolfgang Baumeister

**Author notes:** These authors contributed equally to this work (Florian Wilfling, Chia-Wei Lee). Deceased during the course of this study (Stefan Jentsch).

## Abstract

Autophagy eliminates cytoplasmic content selected by autophagy receptors, which link cargoes to the membrane bound autophagosomal ubiquitin-like protein Atg8/LC3. Here, we discover a selective autophagy pathway for protein condensates formed by endocytic proteins. In this pathway, the endocytic yeast protein Ede1 functions as a selective autophagy receptor. Distinct domains within Ede1 bind Atg8 and mediate phase separation into condensates. Both properties are necessary for an Ede1-dependent autophagy pathway for endocytic proteins, which differs from regular endocytosis, does not involve other known selective autophagy receptors, but requires the core autophagy machinery. Cryo-electron tomography of Ede1-containing condensates – at the plasma membrane and in autophagic bodies – shows a phase-separated compartment at the beginning and end of the Ede1-mediated selective autophagy pathway. Our data suggest a model for autophagic degradation of membraneless compartments by the action of intrinsic autophagy receptors.

**Highlights:**

- Ede1 is a selective autophagy receptor for aberrant CME protein assemblies
- Aberrant CME assemblies form by liquid-liquid phase separation
- Core autophagy machinery and Ede1 are important for degradation of CME condensates
- Ultrastrucural view of a LLPS compartment at the PM and within autophagic bodies

## Introduction

Macroautophagy, hereafter referred to as autophagy, is a highly conserved and ubiquitous process that eliminates bulky cytosolic components in response to starvation and cellular stresses. Ultimately, cargoes destined for autophagic elimination are sequestered into a *de novo* synthesized double-membrane autophagosome that fuses with the vacuole/lysosome for degradation (Dikic, 2017; Wen and Klionsky, 2016). Biogenesis of autophagosomes is initiated at the phagophore assembly site or pre-autophagosomal structure (PAS) in close association with the endoplasmic reticulum (ER) by hierarchical recruitment of conserved autophagy protein machineries (Carlsson and Simonsen, 2015; De Tito et al., 2020; Hurley and Young, 2017; Mizushima et al., 2011; Noda and Inagaki, 2015).

Specific cargoes are recruited to autophagosomes by virtue of interaction with so-called selective autophagy receptors (hereafter called autophagy receptors), which also often display <u>A</u>tg8-interacting <u>m</u>otifs (AIMs) (Nakatogawa et al., 2009; Reggiori et al., 2012). Autophagy receptor AIM sequences in turn bind Atg8, a ubiquitin-like protein, covalently linked to phosphatidylethanolamine (PE) in the isolation membrane (Farre and Subramani, 2016; Ohsumi, 2001). Thus, autophagy receptors couple their cargoes to the autophagosome biogenesis pathway, and allow packaging and subsequent disposal of a wide range of cytosolic entities.

Much of our understanding of selective autophagy derives from matching receptors with important cargoes, such as organelles, aggregates or large macromolecular complexes (Behrends et al., 2010; Dikic and Elazar, 2018; Gatica et al., 2018; Kirkin, 2020; Kirkin and Rogov, 2019). However, our knowledge of molecular mechanisms underlying selective sequestration of autophagic cargoes remains rudimentary. Interestingly, the mammalian autophagy receptor p62, which targets ubiquitylated misfolded proteins for autophagic degradation, phase separates into condensates through multivalent interactions established with multiple ubiquitins linked in chains (Sun et al., 2018). Moreover, the Cvt-cargo itself is a complex that forms condensates through phase separation (Yamasaki et al., 2020). In both cases, the AIM sequences display relatively low affinity for Atg8, and thus concentration of cargo-receptor complexes prior to their enclosure has been proposed to facilitate avid interaction with multiple Atg8 molecules linked to a membrane (Johansen and Lamark, 2020; Kirkin and Rogov, 2019; Zaffagnini and Martens, 2016). Moreover, these findings raise the tantalizing possibility liquid-liquid phase separation may contribute to receptor and cargo selection in other ways in other selective autophagy pathways.

Here, using an unbiased approach to identify Atg8-interacting proteins, we report a previously undescribed autophagy pathway for phase separated endocytic proteins deposits. Normally, proteins mediating clathrin-mediated endocytosis (CME) assemble and disassemble into complex protein and membrane machineries in a highly orchestrated manner, to ultimately transport diverse cargo molecules from the cell surface to the interiors. CME depends on coordinated stepwise plasma membrane (PM)-localized assemblies of over 60 different proteins mediating a temporally-ordered series of membrane and protein transformations (Goode et al., 2015; Lu et al., 2016; McMahon and Boucrot, 2011; Merrifield and Kaksonen, 2014). In the early phase of CME, endocytic proteins and cargoes accumulate at the PM, followed by invagination and scission of the vesicle in the late phase (Kukulski et al., 2012). Failure in the assembly, particularly during the early phase where actin is not yet recruited, leads to clustering of unproductive endocytic protein assemblies with varying stability in both yeast and mammalian cells (Boeke et al., 2014; Kirchhausen et al., 2014; Mettlen et al., 2018). However, both the drivers of CME protein clusters during impaired endocytosis, and the fates of the clusters, remain unknown. We uncover an unexpected role of autophagy in the turnover of atypical deposits containing CME proteins. Our data suggest that the key early CME protein Ede1 has a dual role, and also functions as intrinsic autophagy receptor, via AIM-dependent recruitment of Atg8 to sites of atypical CME protein assembly. We further show the material and cellular structural properties of these endocytic protein deposits as phase separated assemblies, describe molecular features of Ede1 contributing to deposit formation and their autophagic degradation, and cellular structural properties of these phase separated assemblies from their beginning at the PM to their ultimate delivery to autophagosomes. Our findings thus provide a route for endocytic protein turnover via atypical assemblies, show a role for phase separation in cargo recruitment and degradation in selective autophagy, and illuminate cellular structural mechanisms depending on phase separation to mediate protein delivery from the PM and cytoplasm into a specialized membrane-bound organelle.

## Results

### Quantitative proteomics identifies Atg8 binding to clathrin proteins

We sought to uncover unknown autophagy pathways through a proteomics approach identifying receptors and cargoes by virtue of their binding to Atg8 in *Saccharomyces cerevisiae* (Figure 1A). Specifically, we performed label-free quantitative affinity-purification mass spectrometry of GFP pulldowns from a genomically modified yeast strain expressing N-terminally <u>e</u>nhanced <u>g</u>reen fluorescent <u>p</u>rotein (eGFP)-tagged Atg8 and treated with rapamycin, a drug that activates the autophagy pathway by inactivating the TORC1 complex (Noda and Ohsumi, 1998). 225 proteins were reproducibly and statistically significantly enriched between biological replicates (Figure S1A). Among those, we found the Atg8-conjugation machinery (Atg3, Atg4 and Atg7), which covalently links Atg8 to the lipid phosphatidyl ethanolamine (Ohsumi, 2001), as well as other proteins involved in autophagic processes such as the cytoplasm to vacuole targeting (Cvt) pathway receptor Atg19 (Figure 1B). Our approach also revealed autophagic cargo proteins such as Ape1, the target of the Cvt pathway, indicating that not only direct interactors of Atg8 but also receptor-cargo complexes were preserved (Figure 1B). Gene Ontology (GO-term) analysis for the 225 specific eGFP-Atg8 interactors revealed distinct macromolecular complexes, such as the 40S ribosome, the nuclear pore complex or the clathrin-mediated endocytosis machinery (Figure 1C, highlighted by STRING analysis of protein-protein interaction networks (Figure S1B)). Strikingly, the Atg8 pulldown significantly enriched many proteins involved in the early steps of CME (Figures 1D and S1C). In contrast, the actin machinery, which is recruited during the late phase, was not enriched (Figure S1D).

**Figure 1.**
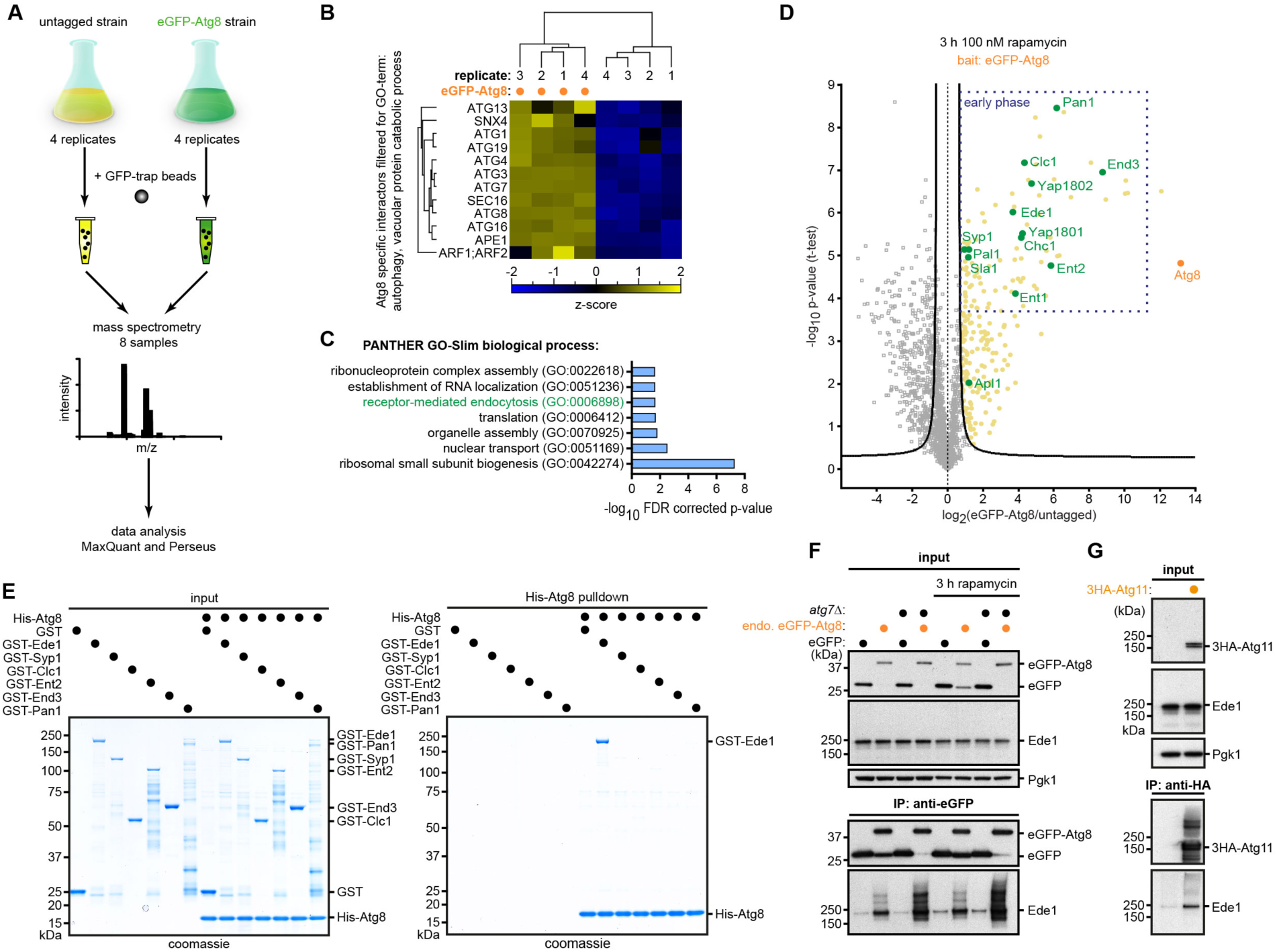
Quantitative proteomics identifies the clathrin endocytosis machinery as autophagy substrate. **(A)** Workflow for the eGFP-Atg8 pulldown and quantitative MS (qMS) analysis. Four biological replicates of a strain expressing eGFP-Atg8 under the control of the *ADH* promoter or four replicates of a control strain were subjected to immunoprecipitation against eGFP after the induction of autophagy with 100 nM rapamycin for 3 hours. The different replicates were analyzed by label-free qMS. **(B)** Hierarchical cluster analysis of label-free eGFP-Atg8 pulldowns after qMS analysis confirms known Atg8 interactors. eGFP-Atg8 specific interactors were filtered for GO-terms autophagy and vacuolar protein catabolic process. Matching ANOVA-positive and FDR-corrected hits are displayed by hierarchical clustering of the respective z-scores for each sample. **(C)** Gene Ontology GO-term analysis of proteins enriched in the eGFP-Atg8 pulldown. GO enrichment analysis was performed using the GENEONTOLOGY search engine (http://geneontology.org). **(D)** Volcano plot representing Atg8-interacting proteins. Depicted are logarithmic ratios of protein intensities against negative logarithmic *p*-values of *t*-test performed from quadruplicates. The hyperbolic curve separates specifically interacting proteins (marked with yellow) from background (square; grey) (FDR: 0.05, S0: 1.0). eGFP-Atg8 interactors involved in CME are marked in green. eGFP-Atg8 is marked in orange. Dark blue box indicates endocytic machinery proteins involved in the early phase of endocytosis. **(E)** Atg8 associates specifically and directly with purified Ede1 *in vitro*. Ni-NTA pulldown of recombinant His-tagged Atg8 (His-Atg8, 8 nM) with different recombinant GST-tagged endocytic machinery proteins (0.8 nM each) including Ede1, Syp1, Clc1, Ent2, End3, or Pan1. **(F)** Validation of Ede1-Atg8 interaction by coIP of eGFP-Atg8 expressed under the endogenous promoter followed by Western blot in WT and *atg7*Δ cells with or without treatment with 100 nM rapamycin for 3 hours. A strain expressing eGFP under the Atg8 promoter serves as control. Pgk1 serves as loading control. **(G)** The autophagic scaffold protein Atg11 interacts with Ede1 *in vivo*. Atg11 expressed under the *ADH* promoter was 3HA-epitope-tagged, and its interaction with Ede1 was probed by co-immunoprecipitation and immunoblotting with anti-Ede1 antibody. *See also Figure S1*.

Given that almost the complete early CME machinery is found enriched in Atg8 pulldowns, we hypothesized that at least one of these proteins might directly bind to Atg8. To test this, we purified several recombinant clathrin coat assembly proteins that were present in our qMS experiment and incubated them *in vitro* with recombinant Atg8. Of these proteins, only Ede1 bound Atg8 (Figure 1E). Immunoprecipitation (IP) from yeast lysates confirmed the interaction of endogenously expressed Ede1 and Atg8 (Figure 1F). This was enhanced when Atg8 lipidation and autophagy were blocked by deletion of *ATG7*, suggesting that the Ede1-Atg8 complex is subject to autophagic turnover (Figure 1F). The interaction was only mildly increased when autophagy was activated by the addition of rapamycin (Figure 1F), suggesting a nutrient-independent function.

To gain insights into the function of Ede1 binding to Atg8, we considered that most of the known autophagy receptors not only bind Atg8 but also to the scaffold protein Atg11 to locally activate isolation membrane formation (Gatica et al., 2018; Torggler et al., 2016). Indeed, we detected Ede1 and Atg11 interaction by co-immunoprecipitation (coIP) (Figure 1G). Together, our data identify the early CME machinery as binding partners of Atg8, and point toward Ede1 as a potential autophagy receptor.

### The early endocytic protein Ede1 recruits Atg8 directly via AIM interactions

Next, we sought to determine the mechanism of Ede1 binding to Atg8. Ede1 is a multidomain protein containing three N-terminal Eps15-homology (EH) domains that bind the NPF tri-peptide motif, followed by proline-rich (PP) and central coiled-coil (CC) domains. At the C-terminus, it harbors a ubiquitin-associated domain (UBA), important for binding ubiquitylated cargo proteins (Boeke et al., 2014; Lu and Drubin, 2017). However, none of these domains are known to bind Atg8. Because most known Atg8-binding proteins display a so-called Atg8-interacting motif (AIM in yeast, aka LC3-interacting region LIR in higher eukaryotes) sequence [(W/F/Y)xx(L/I/V) flanked by at least one proximal acidic residue] that binds to two adjacent hydrophobic pockets in Atg8 (Noda et al., 2010), we used as control an Atg8 (Y49A, L50A) mutant impaired for binding in both sites and unable to bind AIM-containing proteins (Ho et al., 2009). The mutations abrogated Ede1-Atg8 interaction, supporting the idea that Ede1 is an AIM-dependent receptor for Atg8 (Figure 2A).

**Figure 2.**
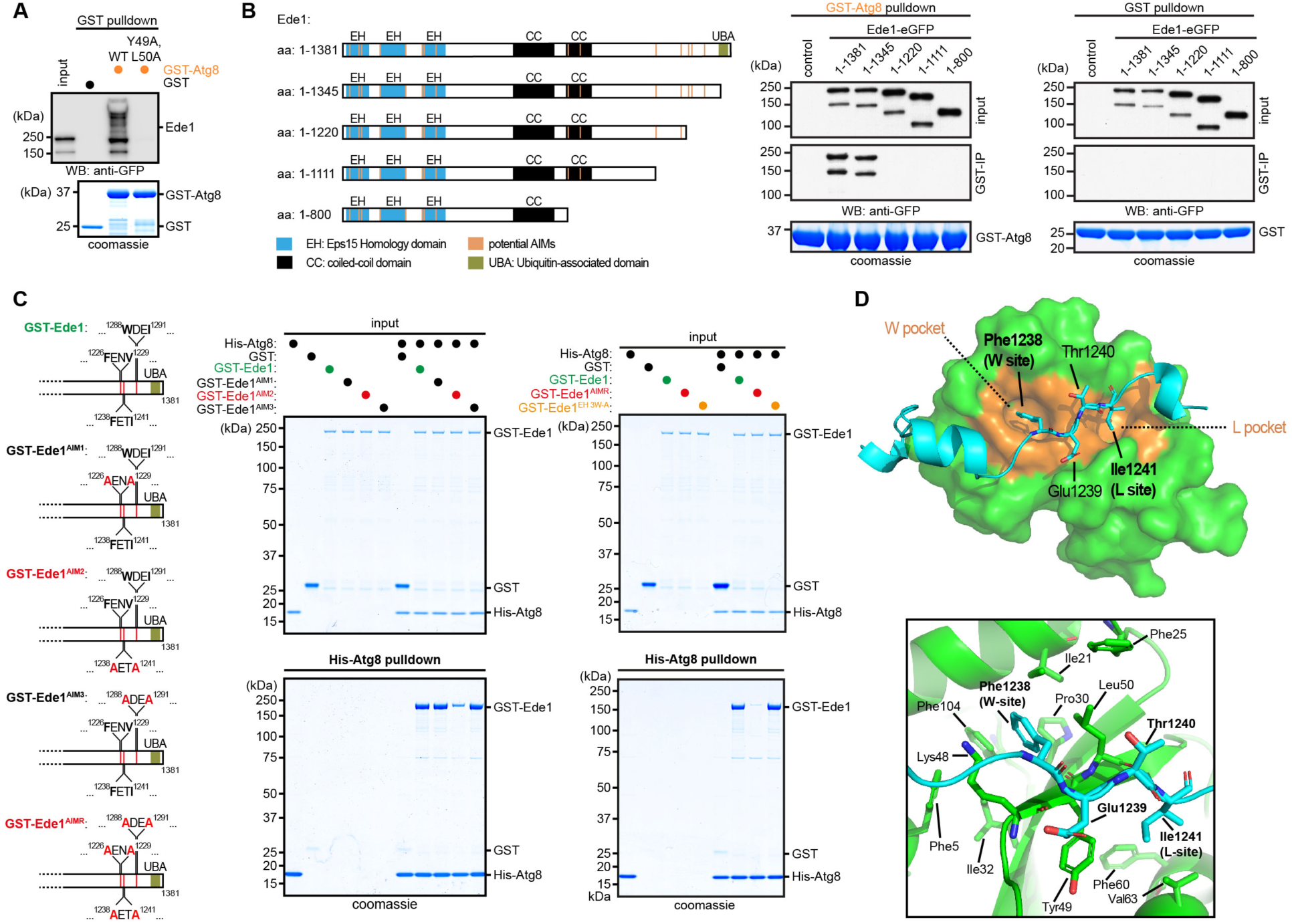
Ede1 recruits Atg8 directly via classical AIM interactions. **(A)** Ede1 binds to Atg8 in an AIM-dependent fashion. Yeast cell lysates from a WT strain were incubated with recombinant His-tagged Atg8 or the Y49A, L50A mutant (defective in AIM-dependent interactions, (Ho et al., 2009)), followed by Ni-NTA pulldown and immunoblotting with anti-Ede1 antibody. AIM: Atg8-interacting motif. **(B)** Deletion of a 161 amino acid C-terminal fragment of Ede1 abolishes binding to Atg8. Atg8-deficient binding mutants of Ede1 are identified by GST pulldown from lysates expressing different eGFP-tagged C-terminal deletion mutants incubated with recombinant GST-Atg8 or GST alone and immunoblotting with anti-GFP antibody. Illustration shows truncation mutants used to map the Atg8 binding site. EH: Eps15-homology domain; CC: coiled-coil domain; UBA: ubiquitin-associated domain; aa: amino acid. **(C)** The binding between Ede1 and Atg8 is mediated by classical AIMs. Ni-NTA pulldown of recombinant His-tagged Atg8 (His-Atg8, 8 nM) with GST-tagged Ede1 wildtype, or single AIM mutants (Ede1^AIM1^, Ede1^AIM2^ or Ede1^AIM3^, 0.8 nM each) (left), or triple AIM mutant (Ede1^AIMR^, 0.8 nM) (right). To exclude NPF-tripeptide binding to the EH domain, point mutants were introduced in each of the EH domains of Ede1 (Ede1^EH 3W-A^, 0.8 nM) (Dores et al., 2010). Illustration shows position of mutants used to map the AIMs. UBA: ubiquitin-associated domain. **(D)** Crystal structure of Ede1(1220-1247) peptide binding to Atg8 reveals that Ede1(1238-1241) (AIM2) serves as a classical AIM. The association of Ede1 peptide to Atg8 features Ede1 Phe1238 and Ile1241 docking in the W-pocket and the L-pocket of Atg8 respectively (top). AIM2 binding to Atg8 is achieved by extensive hydrophobic interactions among its W/L-sites with the same set of Atg8 residues engaged as characterized in other classical AIM studies (bottom) (Johansen and Lamark, 2020). *See also Figure S2*.

We computationally predicted potential AIMs in Ede1 and created C-terminal truncations lacking them in various combinations (Kalvari et al., 2014). Deletion of the last three predicted AIMs abolished the binding of Ede1-eGFP to GST-Atg8, while deletion of the C-terminal UBA domain alone had no effect (Figure 2B). Notably, AIM motifs two and three (AIM2/3) are conserved among different yeast species (Figure S2A) and are located in a large intrinsically disordered region (Figure S2B), a typical feature of most validated AIMs (Popelka and Klionsky, 2015). To test roles of individual AIMs, we expressed and purified GST-Ede1 constructs with all possible combinations of AIMs inactivated by alanine replacements for the key residues, and checked binding to recombinant Atg8 (Figures 2C and S2C). Mutations in AIM2 (Ede1^AIM2^) strongly impaired the interaction (Figure 2C). Double mutation of AIM2 and AIM3 (Ede1^AIM2+3^) further reduced binding (Figure S2C), which was lost completely when all the putative AIM motifs were mutated (Ede1^AIMR^) (Figure 2C). We further showed that these Ede1 AIMs are sufficient to bind Atg8 using a peptide-binding assay (Figure S2D). As control, mutations in the EH domains (Ede1^EH 3W-A^) which bind NPF-tripeptide (Dores et al., 2010), had no influence on the binding between Atg8 and Ede1 (Figure 2C).

To determine the structural basis for the Atg8/Ede1 interaction, we pursued a crystal structure of Atg8 in complex with Ede1 fragments. We obtained crystals with an Ede1 region (aa: 1220-1247) containing AIM1 and AIM2 (Figure 2D and Table S1). A high-resolution structure showed that AIM1 folds within a short helix, and the aromatic Phe1238 of AIM2 engages the W-site of Atg8 and the Ile1241 the L-site, whereas the negatively charged Asp1242 forms ionic interactions with Arg28, reassembling other known classical AIM-Atg8 interactions (Kirkin and Rogov, 2019). Ede1 also harbours additional Atg8-binding elements including a few acidic residues upstream and a helix downstream to AIM2 (Figure S2E). Intriguingly, similar interactions were observed previously between a mammalian GABARAP Atg8 ortholog and ankyrin protein, where the pre-LIR acidic side-chains and a post-LIR helix stabilized the complex (Figure S2F) (Li et al., 2018). This raises a possibility that similar extended binding patterns supplemental to the existing canonical LIR/AIM-binding mechanism may play a more general and important role in Atg8-family recruitment (Wirth et al., 2019).

### Ede1 is necessary for Atg8 binding to aberrant CME protein deposits

To ascertain if Ede1 displays properties of an autophagy receptor, we introduced the *ede1*^AIMR^ mutant into the endogenous *EDE1* locus. Analysis of eGFP-Atg8 co-immunoprecipitates by Western blotting confirmed that Atg8 binding to Ede1 *in vivo* depends on the AIMs (Figure 3A). Because AIM-dependent binding to Atg8 is often a feature of selective autophagy receptors, we first tested a role in binding the conventional cargo Ape1. Mutation of Ede1 did not affect Atg8 recruitment of this Cvt cargo, nor did deletion of *APE1* impact the interaction between Ede1 and Atg8 (Figure 3A). Moreover, mutating the AIMs in Ede1 had no effect on the rate of autophagosomal biogenesis, bulk autophagy or any other selective autophagy pathways examined (Figures S2G-K).

**Figure 3.**
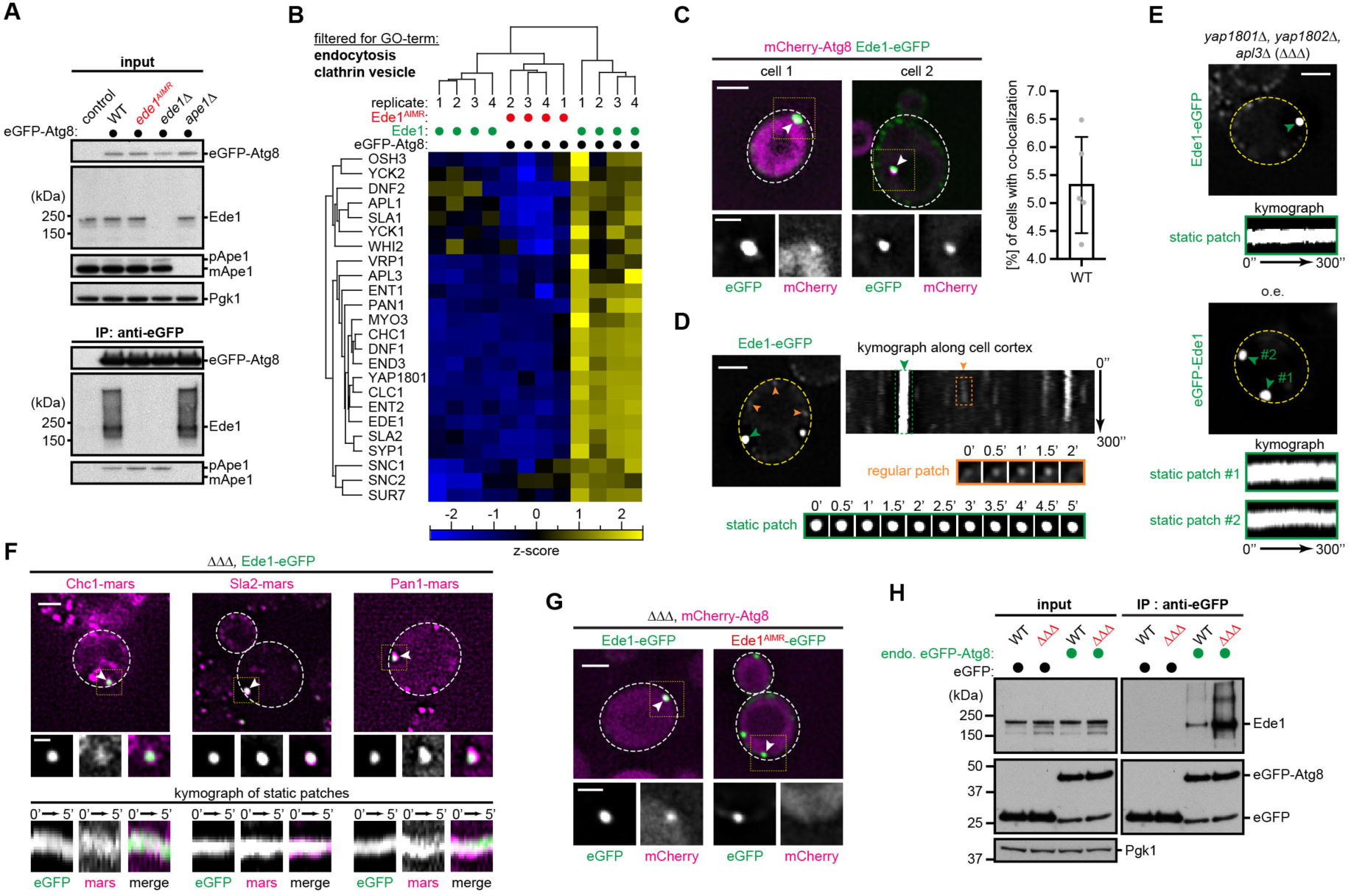
Ede1 serves as a receptor for atypical CME assemblies. **(A)** Ede1 interacts with Atg8 in an AIM-dependent manner also *in vivo*. Validation of the AIM-dependent binding of Ede1 to Atg8 in WT, Ede1^AIMR^-mutated (*ede1*^AIMR^), Ede1-deficient (*ede1*Δ), or Ape1-deficient (*ape1*Δ) cells after 3 hours of 100 nM rapamycin treatment by immunoprecipitation of eGFP-Atg8 and immunoblotting with the respective antibodies. Pgk1 was used as a loading control. **(B)** Clathrin-mediated endocytic proteins bind Atg8 specifically through Ede1. Hierarchical cluster analysis filtered for the GO-term (endocytosis) for all Ede1-dependent Atg8 interactors identified by qMS (Figure S3A, red box). **(C)** mCherry-Atg8 (p*ADH*) and Ede1-eGFP (endogenous promoter) co-localize at distinct sites of Ede1 clustering in a subset of wildtype cells. Quantification was performed for at least 500 cells (n = 5). Lower panels show magnifications of boxed areas. Scale bar represents 2 µm and 1 µm for inset. **(D)** Ede1 clusters are distinct from typical endocytic patches. Ede1-eGFP forms stable accumulations in a subpopulation of wildtype cells, which are distinct from regular endocytic patches (left). Kymograph representation of a movie (1 frame/5 s) of cells expressing Ede1-eGFP (endogenous promoter) in wildtype cells (right). Scale bar represents 2 µm. Orange arrows and box highlights regular endocytic Ede1 patches; green highlights atypical Ede1 clusters. **(E)** Atypical stable accumulations of Ede1 are particularly detected in early phase endocytic mutants. Deletion of the endocytic adopter proteins *yap1801*Δ, *yap1802*Δ, and *apl3*Δ (hereafter *ΔΔΔ*) or overexpressing of Ede1 (p*ADH*) triggers stable clustering of Ede1. Kymograph representation of a movie (1 frame/5 s) of cells expressing Ede1-eGFP in the ΔΔΔ background (top) or cells overexpressing eGFP-Ede1 under *ADH* promoter (bottom). Scale bar represents 2 µm. Green highlights atypical Ede1 clusters. **(F)** The CME machinery proteins Chc1, Sla2 and Pan1 stably co-localize at sites of Ede1-eGFP clustering in ΔΔΔ cells. Kymograph representation of a two-color movie (1 frame/20 s) of cells expressing Ede1-eGFP and either Chc1-mars, Sla2-mars or Pan1-mars in the ΔΔΔ background. Lower panels show magnifications of boxed areas. Scale bar represents 2 µm or 1 µm for inlets. **(G)** Atg8 co-localizes AIM-dependent with Ede1-eGFP clusters in *ΔΔΔ* mutant cells. Colocalization experiments between mCherry-Atg8 (p*ADH*) and Ede1-eGFP or Ede1^AIMR^-eGFP in the ΔΔΔ strain background. Lower panels show magnifications of boxed areas. Scale bar represents 2 µm and 1 µm for inset. **(H)** Ede1-Atg8 interaction is increased when the early phase of endocytosis is impaired. CoIP of eGFP-Atg8 (endogenous promoter) followed by Western blot in WT and ΔΔΔ cells. Strains expressing GFP under the Atg8 promoter serve as control. n = 3 biologically independent experiments. *See also Figures S2, S3 and S4*.

Considering the original proteomics results (Figure 1), since among the Atg8-binding CME proteins tested, only Ede1 directly recruits Atg8, we wondered if the other Atg8-binding endocytosis proteins identified might be recruited to Atg8 via Ede1. To verify this idea, we designed a qMS experiment in which the eGFP-Atg8 interactome from wildtype cells was compared to cells with the Atg8-binding deficient mutant *ede1*^AIMR^. As control served an untagged Atg8 strain. As expected, we found three types of significant binders by multiple-sample testing and cluster analysis (Figure S3A). The first category binds Atg8 specifically in an Ede1-dependent manner (Ede1-dependent Atg8 binders), whereas the second binds Atg8 independently of Ede1 (Atg8 binders). The last type of interactors are unspecific binders, which bind to the beads/antibody in the absence of the eGFP antigen. GO term enrichment analysis showed “endocytosis”, “plasma membrane”, and “transmembrane transporter activity” significantly enriched among Ede1-dependent Atg8 binders (Figures S3B-C). Importantly, a clear Ede1-dependent enrichment of many endocytosis machinery proteins involved in the early stages of CME could be seen (Figure 3B) and confirmed in multiple cases (Syp1, Chc1, Clc1, Yap1801, Yap1802, Apl1, Sla1, Sla2, Ent1, Ent2, and End3) by coIP-western experiments (Figure S3D).

To gain insights into functional roles of Ede1-Atg8 interactions, we sought to visualize their properties in cells. We fluorescently tagged both proteins with either eGFP or mCherry, respectively. Using live-cell imaging, Ede1 and Atg8 showed colocalization in ∼5% of wildtype cells at sites of strong Ede1 accumulation (Figure 3C). These Ede1 clusters are morphologically distinct from typical endocytic patches: time-lapse microscopy showed that in contrast to sites of active endocytosis in which Ede1 leaves the endocytic assembly within 30 – 240 sec (Lu et al., 2016), the foci with Atg8 are stable and persist throughout the experiment (5 min) (Figure 3D). To exclusively monitor these sites of Ede1/Atg8 interaction, we turned to a bimolecular fluorescence complementation (BiFC) strategy (Kerppola, 2008) by labelling Atg8 and Ede1 with either the N-terminal (VN) or C-terminal (VC) Venus fragments. Indeed, the fluorescent BiFC signal was observed at few sites, close to the PM, but distinct from regular sites of endocytosis (Figure S4A). Importantly, the BiFC signal was abolished in *ede1^AIMR^* mutant cells, although protein levels were comparable (Figure S4B), validating the specificity of this system.

Are these observed foci, sites where Ede1 mediates the binding of endocytic proteins to Atg8? Indeed, mars-tagged Chc1, Syp1, Sla2, End3, and Pan1 all colocalized at Ede1/Atg8 BiFC foci (Figure S4C). Notably, these clusters, that sporadically occurred in proliferating wildtype cells, resembled those that have been reported to accumulate when Ede1 is overexpressed or when a specific combination of early endocytosis adaptor proteins is deleted (*yap1801Δ, yap1802Δ, apl3Δ*; hereafter called ΔΔΔ) (Boeke et al., 2014). Thus, to test if the Ede1/Atg8 interaction coincides with such defective endocytic protein assemblies, we performed time-lapse microscopy of eGFP-tagged Ede1 in both the *ΔΔΔ* mutant or the Ede1 overexpression strain. Indeed, focal accumulations of eGFP-tagged Ede1 are stable and persist during the whole duration of the experiments in both strains (Figure 3E), and they resemble the sites of Ede1/Atg8 interaction seen in wildtype cells. Moreover, both strains likewise exacerbated accumulation of other CME proteins in these stable, non-endocytic foci (Figures 3F and S4D). The strong clustering of Ede1 was independent of Ede1 tagging with eGFP, since cells overexpressing untagged Ede1 showed strong focal accumulation of Sla2 or Pan1 tagged with the monomeric TagGFP2 protein (Figure S4E). To explore if Ede1 recruits Atg8 to sites of aberrant CME assembly, we monitored fluorescently tagged Atg8 and Ede1 in *ΔΔΔ* mutant or Ede1 overexpressing cells. Under these conditions, Atg8 colocalized with Ede1 only at sites of aberrant Ede1 clustering (Figures 3G and S4F). This was dependent on the AIM region in Ede1 since mCherry-Atg8 failed to colocalize with Ede1 clusters in *ede1*^AIMR^ mutant cells (Figures 3G and S4F). If defects in endocytic protein assembly triggers Atg8 recruitment to these sites, we hypothesized that binding of Ede1 to Atg8 should be increased in the *ΔΔΔ* mutant compared to wildtype. Indeed, we observed enhanced interaction between eGFP-Atg8 and Ede1 measured by eGFP coIP, suggesting that Ede1 recruits Atg8 particularly to aberrant sites of CME assembly (Figure 3H).

Taken together, the data show: (1) Ede1 co-recruits early CME proteins to Atg8, (2) sites of their colocalization do not display normal endocytic mobility, and (3) they accumulate in yeast impaired for endocytosis. We refer hereafter to these atypical Ede1 clusters as *E*de1-dependent e*N*docytic protein *D*eposit (END), which have similar properties and composition in WT and mutant cells, but are increased when endocytosis is impaired.

### Ede1 and the core autophagy machinery are necessary for autophagic degradation of atypical endocytic protein deposits

Given that Atg8 is a constituent of the END assemblies, we hypothesized a role for autophagy. To explore this, we first tested whether Ede1 is subject to vacuolar degradation using the standard eGFP fusion assay, where eGFP persists after liberation upon degradation of the fusion partner in the vacuole; the compact fold of eGFP renders it resistant to vacuolar hydrolases (Cheong and Klionsky, 2008; Shintani and Klionsky, 2004). To allow monitoring of eGFP cleavage under rich growth conditions, we performed GFP-trap pulldowns followed by Western blot analysis of endogenously Ede1-eGFP tagged strains (Figure 4A). This led to enrichment of free eGFP which depended on both Atg8 and on Ede1 AIM motifs, indicating that some Ede1-eGFP is constitutively subjected to autophagic degradation (Figures 4B-C). Moreover, this autophagic degradation of Ede1, as monitored by monomeric eGFP, was enhanced by mutations impairing CME, such as the *ΔΔΔ* strain or knockout of clathrin light chain (*clc1Δ*) known to cause persistent patch formation of early CME proteins (Carroll et al., 2012) (Figures 4B and S5A).

**Figure 4.**
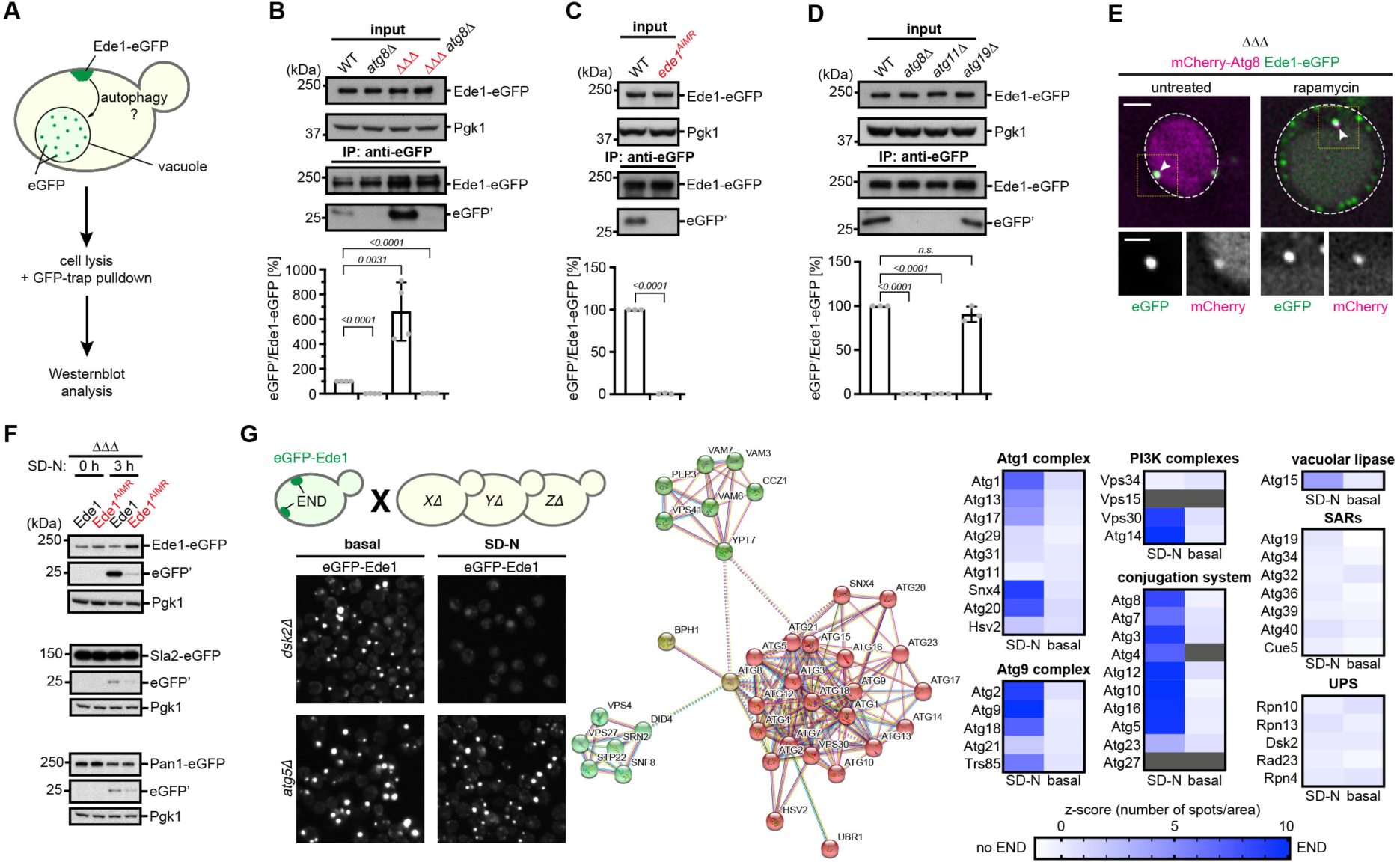
Ede1-dependent degradation of aberrant CME assemblies by the core autophagy machinery. **(A)** Workflow of monitoring endogenously expressed Ede1-eGFP cleavage during logarithmic cell growth in wildtype cells. Wildtype cells are lysed and subjected to coIP using GFP-trap resin. Both full length Ede1-eGFP and monomeric eGFP are monitored by Western blot analysis. **(B)** Autophagic degradation of Ede1 is observed during logarithmic growth in WT, and enhanced in ΔΔΔ mutant cells. eGFP pulldowns were performed in WT or ΔΔΔ mutant cells expressing endogenously eGFP-tagged Ede1. The pulldown of vacuolar enriched free eGFP from WT or ΔΔΔ mutant was compared to cells additionally deficient in the core autophagy protein Atg8 (*atg8*Δ). A quantification of the ratio between free eGFP’ and full length Ede1-eGFP is shown. Data are mean ± s.d. of n = 4 biologically independent experiments. Statistical analysis was performed using two-tailed Student’s *t*-tests, *p*-values are indicated. Pgk1 serves as loading control for the input. **(C-D)** Autophagic turnover of Ede1 is AIM-dependent as well as dependent on Atg11 but is unaffected by Atg19. Same experimental setup as in (B), but with either *ede1*^AIMR^ cells (C), or *atg11*Δ and *atg19*Δ mutant cells (D) compared to WT cells. Data are mean ± s.d. of n = 3 biologically independent experiments. Statistical analysis was performed using two-tailed Student’s *t*-tests, *p*-values are indicated. Pgk1 serves as loading control for the input. **(E)** Ede1-eGFP is targeted to the vacuole upon rapamycin treatment. Recruitment of Atg8 to sites of Ede1 accumulation is visualized by colocalization of mCherry-Atg8 (p*ADH*) and Ede1-eGFP in the ΔΔΔ strain background before and after 100 nM rapamycin treatment for 3 hours. Arrow on the left image shows colocalization at sides of Ede1 clustering. Arrow on the right image shows colocalization at the PAS. Lower panels show magnifications of boxed areas. Scale bar represents 2 µm and 1 µm for inset. **(F)** END accumulated Sla2 and Pan1 are preferentially degraded by autophagy. The mid coat protein Sla2 or the late coat protein Pan1 were C-terminally eGFP-tagged in WT or ΔΔΔ (*yap1801Δ*, *yap1802Δ*, *apl3Δ*) cells with endogenous expression of Ede1 or Ede1^AIMR^, and its starvation-induced autophagic processing was analyzed by eGFP-cleavage assay. Top shows degradation of Ede1-eGFP or Ede1^AIMR^-eGFP under the same conditions. **(G)** A Systematic screen reveals particularly core proteins involved in autophagy as important factors for the END degradation. eGFP-Ede1 was crossed against a single gene knockout library. Cells were imaged by fluorescence microscopy under rich condition and after after 24 hours of nitrogen starvation. The protein-hits from the screen were subjected to STRING, and hierarchical cluster analyses. *See also Figure S5*.

We next asked if autophagy plays a role in END degradation, by testing effects of deletions of the autophagic scaffold protein Atg11 important for most selective autophagy pathways, or of Atg19 that is a selective autophagy receptor for the Cvt but not other autophagy pathways (Lynch-Day and Klionsky, 2010). Enrichment of free eGFP depended on the autophagic scaffold protein Atg11, but not Atg19, raising the possibility of phagophore assembly at sites of Ede1 accumulation under conditions that do not stimulate bulk (non-starvation) or other selective forms of autophagy (Figure 4D). In agreement with these results, we obtained a similar BiFC signal for Atg11 and Ede1 labelled with either the VC or VN respectively as seen for Ede1 and Atg8 (Figure S5B). Additionally, we observed colocalization of RFP-Atg8 with the Ede1-VN VC-Atg11 BiFC signal indicating the recruitment of the autophagy machinery to the same sites (Figure S5C). Vacuolar accumulation of monomeric eGFP could also be visualized by fluorescence microscopy of *ΔΔΔ* mutant cells expressing Ede1-eGFP and mCherry-Atg8 after treatment of cells with rapamycin (Figure 4E). Also, eGFP accumulation was detected after endogenous expression of tagged fusions of other CME proteins (Sla2-eGFP or Pan1-eGFP), which we found clustered at END sites. Notably, the Ede1 AIMR-mutation strongly impaired appearance of monomeric eGFP, thus suggesting the Ede1-Atg8 interaction is a determinant of END degradation (Figure 4F).

To explore at a systemwide level which autophagic proteins are crucial for degradation of END assemblies, we performed a high-content microscopy screen by crossing an eGFP-Ede1 overexpression strain against a deletion library of nonessential single yeast gene knockouts (Figure 4G). We then analysed presence of Ede1 clusters before and after nitrogen starvation (Figure 4G). This revealed 100 proteins (z-score higher than 1.5) (Table S2) with stabilized Ede1 clusters after nitrogen starvation, most belonging to the core autophagy machinery, ESCRT and a set of proteins mediating vacuolar docking and fusion (Figure 4G). Detailed analysis of the autophagy pathway revealed a strong dependency on the core autophagy machinery. Notably, under nitrogen starvation Atg11 is not essential which is consistent with previous findings (Matscheko et al., 2019). Moreover, other known autophagy receptors as well as proteins important for proteasomal degradation did not affect degradation of the Ede1 clusters under these conditions (Figure 4G). Our results therefore indicate that END assemblies are degraded by Ede1-dependent autophagic degradation. Moreover, the importance of direct, AIM-dependent Ede1 binding to Atg8 in this process suggests that Ede1 acts as selective autophagy receptor for END protein assemblies.

### ENDs exhibit hallmarks of LLPS

Cargoes of selective autophagy are often discrete entities – membrane-encased organelles in mitophagy or ER-phagy, encapsulated pathogenic microbes in xenophagy, or if proteins then either specific structural assemblies (e.g. Cvt, proteasome-phagy or ferritinophagy) or assemblies with particular material properties in aggrephagy. Thus, we sought to characterize the material properties of these END assemblies. We first examined the mobility of Ede1 within these clusters using Fluorescence Recovery after Photobleaching (FRAP). FRAP experiments with eGFP-Ede1 showed: (1) Ede1 moves dynamically within the aberrant CME assemblies, with a halftime of 1.87 ± 0.76 s (Figure 5A); (2) the eGFP-Ede1 pool within the clusters exchanges with the cytosol, since continuous bleaching of the cytoplasm led to a decrease of eGFP-fluorescence within these clusters (Figure S6A); (3) the majority of Ede1-eGFP does not obviously form insoluble protein aggregates since Hsp104-mars, a marker for cytosolic protein aggregation in yeast (Inoue et al., 2004; Shorter and Lindquist, 2004), was absent from sites of Ede1-eGFP accumulation (Figure 5B).

**Figure 5.**
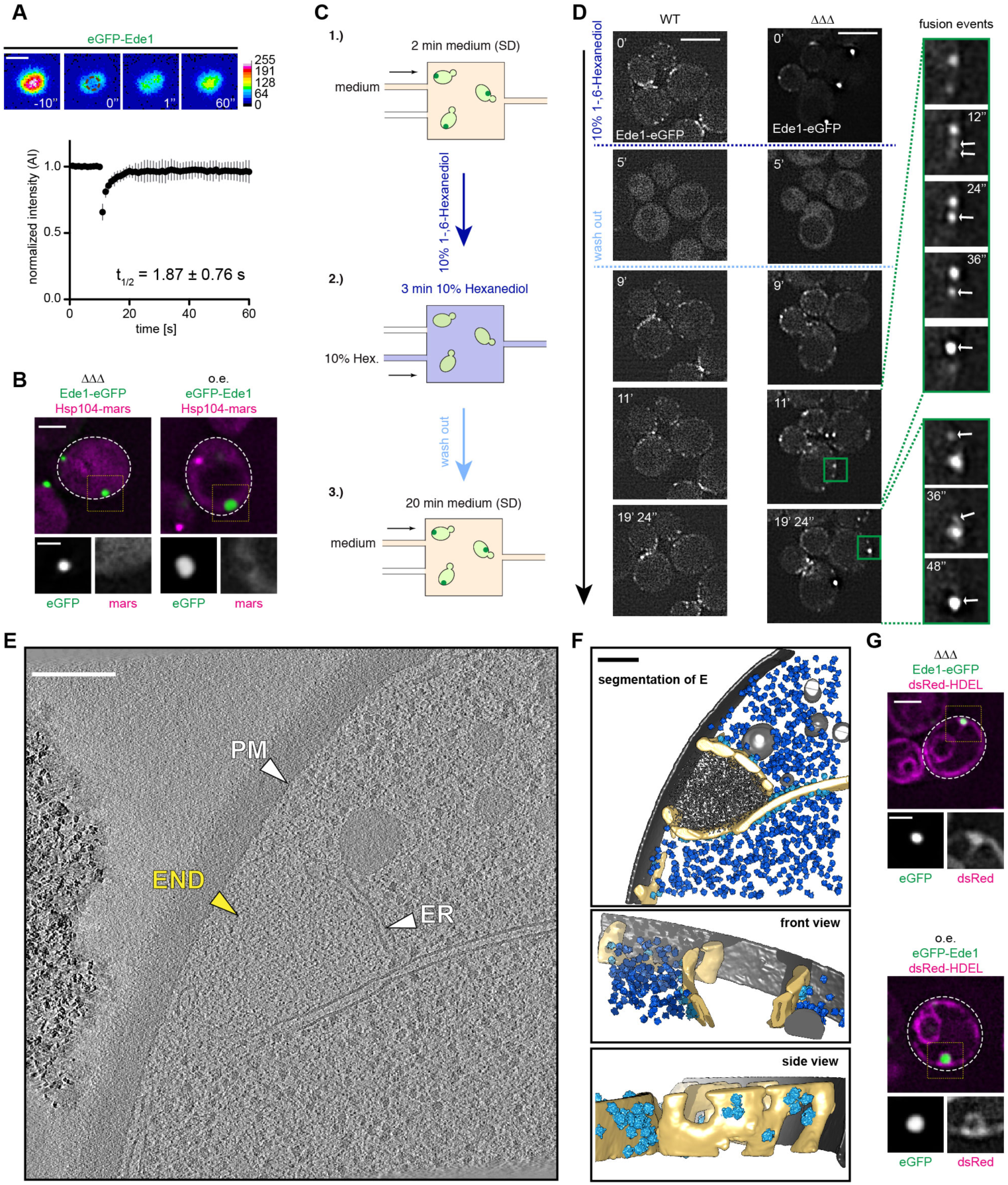
PM-Attached END Displays Hallmarks of LLPS. **(A)** The fraction of PM accumulated eGFP-Ede1 in the END is mobile. Fluorescence recovery after photobleaching experiments were performed for cells overexpressing eGFP-Ede1 under the *ADH* promoter. A fraction within the END was bleached (brown circle) and recovery of the signal was followed over a time course of 60 s (1 frame/1 s). Representative images are shown. Quantification shows the recovery of the eGFP-Ede1 signal from 13 independent experiments as mean ± s.d.. Scale bar represents 1 µm. **(B)** The cytosolic disaggregase Hsp104 is not enriched at sites of Ede1 accumulation as visualized by colocalization experiments between Hsp104-mars and Ede1-eGFP in ΔΔΔ mutant cells (left), or between Hsp104-mars and eGFP-Ede1 in cells where *EDE1* is under the control of the *ADH* promoter (right). Scale bar represents 2 µm and 1 µm for inset. **(C)** Experimental setup to visualize the material properties of END in single cells. Monitoring of ΔΔΔ mutant cells expressing Ede1-eGFP are trapped in a microfluidic device before, during, and after treatment with 10% 1-,6-Hexanediol. After 2 min the medium stream is switched to medium containing 10% 1-,6-Hexanediol and 10 µg/ml digitonin. After further 3 min medium is changed back. Fluorescence images are recorded for 26 min (1 frame/12 s). **(D)** Only ΔΔΔ cells show fusion of individual Ede1-eGFP foci at the PM after Hexanediol treatment. Addition of 10% 1-,6-Hexanediol for 3 min dissolves all endocytic events as monitored by tracking of Ede1-eGFP signal in wildtype or ΔΔΔ mutant cells (1 frame/12 s). Wash out of 1-,6-Hexanediol specifically leads to fusion of individual Ede1-eGFP foci at the PM in ΔΔΔ mutant but not wildtype cells. Scale bar represents 5 µm and 1 µm for inset. **(E)** Correlative cryo-ET targeting Ede1-eGFP signal reveals architecture of the END in ΔΔΔ cells. An average 2D section of the original tomogram shows a region correlated to Ede1-eGFP signal (marked with yellow) in the ΔΔΔ strain. Scale bar represents 200 nm. **(F) Top:** Segmentation of (E) shows the drop-like protein condensate (dark grey) at the PM, which is surrounded by a fenestrated membrane compartment (light yellow). Ribosomes are shown in light (membrane-bound) and dark blue (free); the PM is shown in dark grey; membranes surrounding the protein accumulation are segmented in light yellow. **Middle:** Shown is a sliced-through front view of (E) without the density corresponding to the endocytic protein accumulation. The protein density within the fenestrated ER is clearly devoid of all ribosomes (light and dark blue) and extends directly from the plasma membrane (dark grey). A side view of (E) shows ribosomes in light blue, which are attached to the membrane in a fashion typical for the ER. Membrane-associated ribosomes are exclusively found on parts of the fenestrated ER facing the cytosol. **(G)** The luminal ER marker dsRed-HDEL is adjacent to the END as visualized by colocalization experiments between dsRed-HDEL and Ede1-eGFP in ΔΔΔ mutant cells (top), or between dsRed-HDEL and eGFP-Ede1 in Ede1 overexpression (p*ADH*) cells (bottom). Scale bar represents 2 µm and 1 µm for inset. *See also Figure S6*.

Because the FRAP results are consistent with liquid-like properties, we considered that assemblies within the endocytic machinery are often formed through weak multivalent interactions between a large network of proteins (Smith et al., 2017). Such interactions are often also the driving force for liquid-liquid phase separation (LLPS) of proteins (Banani et al., 2017; Harmon et al., 2017; Martin and Mittag, 2018). Thus, to test if the Ede1-containing clusters are condensates, we tested effects of hexanediol, an aliphatic alcohol that has been used to dissolve liquid-like assemblies in a reversible manner in living cells (Alberti et al., 2019). Single cell experiments using a microfluidics device allowed us to track Ede1-eGFP fluorescence expressed from the endogenous locus in both wildtype and *ΔΔΔ* mutant cells before, during, and after hexanediol treatment (Figure 5C). A three-minute treatment with hexanediol dissolved the endocytic protein assemblies in both strains and led to dispersed cytosolic localization of Ede1. Following wash out, Ede1-eGFP started to re-accumulate at the PM (Figure 5D). Wildtype cells expressing Ede1-eGFP formed regular endocytic patches at the PM (Figure 5D and Movie S1), whereas ΔΔΔ mutant cells additionally showed fusion of individual endocytic patches into larger assemblies (Figure 5D and Movie S2). Collectively, these data suggest that END sites are dynamic compartments containing an ensemble of endocytic proteins exhibiting hallmarks of LLPS. Notably, this conclusion is corroborated by a study posted on bioRxiv during the course of our manuscript preparation (Kozak and Kaksonen, 2019).

### Visualizing END assemblies at the PM by cryo-electron tomography

To get a higher-resolution understanding of the END condensates, we applied correlative cryo-electron tomography (cryo-ET). To enrich for and localize those sites for structural studies *in situ*, we employed the ΔΔΔ strain where Ede1 was endogenously tagged with eGFP. This enabled targeted cryo-ET at sites of correlated fluorescence signal on ∼100 nm thick lamellas obtained by 3D-correlative cryo-focused ion beam (FIB) milling (Figure S6B). Cryo-ET revealed several structural properties of those sites. First, all recorded tomograms showed a distinct assembly with drop-like shape and localized proximal to the PM (Figures 5E-F and S6C-D) (Movie S3). Second, there is no overtly regular or recognizable electron density in the droplet. Third, the area attributed to the endocytic protein condensate is characterized by a striking exclusion of ribosomes, emblematic of distinct assembles or organelles that are either tightly packed or have a specific boundary. Notably, this resembles such exclusion zones observed during normal CME (Figures 5E-F) (Idrissi et al., 2008; Kukulski et al., 2012; Mulholland et al., 1994; Takagi et al., 2003), although the area of condensation defining the END was much larger than regular exclusion zones seen during CME. Fourth, the protein density corresponding to the END condensate was surrounded by tubular membranes which we attribute to the endoplasmic reticulum (ER) based on attached ribosomes (Figures 5E-F) as well as fluorescence imaging of the luminal ER marker dsRed-HDEL (Figure 5G). Intriguingly, the ER was not completely closed around the END condensate, but displayed openings (Figure 5F). This supports the notion that ribosome exclusion must occur by means other than membranes. The same characteristic features were also observed for the strain in which eGFP-Ede1 was overexpressed (Figures S6C-D) (Movies S4-5). The only notable difference was the increased space occupied by the condensate protein density as compared to the *ΔΔΔ* strain, likely due to increased Ede1 levels (Figures S6C-D).

### An Ede1 prion-like domain region is essential for phase separation and autophagic degradation

To elucidate the molecular basis for END phase separation, and to interrogate a potential functional role, we sought to identify a responsible domain. Because another autophagy receptor, p62, undergoes phase separation in the presence of K63 polyubiquitin chains in a manner important for autophagic degradation of p62 bodies (Sun et al., 2018), we first tested roles of ubiquitin binding domains in CME constituents. Ede1 harbours a C-terminal UBA (Ubiquitin-associated) domain, and the epsins Ent1 and Ent2 each display UIMs (Ubiquitin-interacting motif) (Lu and Drubin, 2017). We created a strain (*UBDΔ*) in which ubiquitin binding domains were removed or mutated and simultaneously overexpressed eGFP-Ede1 (Figure S7A). Although the average size of the END was slightly smaller (Figure S7B), phase separation *per se* was not obviously impaired (Figure S7B). Accordingly, we found that Ede1 binding to Atg8 was not impaired in the *UBDΔ* background, whereas binding of polyubiquitinated cargo proteins was strongly reduced (Figure S7C). Furthermore, eGFP accumulated in the vacuole upon subjecting WT or *UBDΔ* cells expressing eGFP-Ede1 to nitrogen starvation (Figure S7D).

As we did not find evidence for ubiquitin-mediated phase separation, we considered whether Ede1’s multidomain structure may encompass a distinct element for phase separation. Many phase-separating proteins encompass intrinsically disordered prion-like domains (PrLDs), which are often necessary for driving intracellular phase transitions (Molliex et al., 2015; Patel et al., 2015). *In silico* analysis of the Ede1 sequence identified a prion-like domain (PrLD) between the N-terminal EH domains and the central coiled-coil region (previously annotated as proline-rich domain) (Figure 6A) (Zambrano et al., 2015). Stepwise deletion from the N-terminus showed that removal of the N-terminal region including the PrLD, but not of the N-terminal EH domains alone, diminished accumulation of Ede1 as observed by the reduced size of the condensates (Figure 6A). Ede1 lost its ability to phase separate and became cytosolic when the predicted PrLD and a C-terminal flanking region was deleted (Ede1^549-1381)^ (Figure 6A). Moreover, in cells expressing Ede1^549-1381^ other CME proteins Chc1 and Sla2 also lost their ability to aberrantly cluster (Figure 6B), suggesting that Ede1’s PrLD domain is essential for phase separation and END site formation.

**Figure 6.**
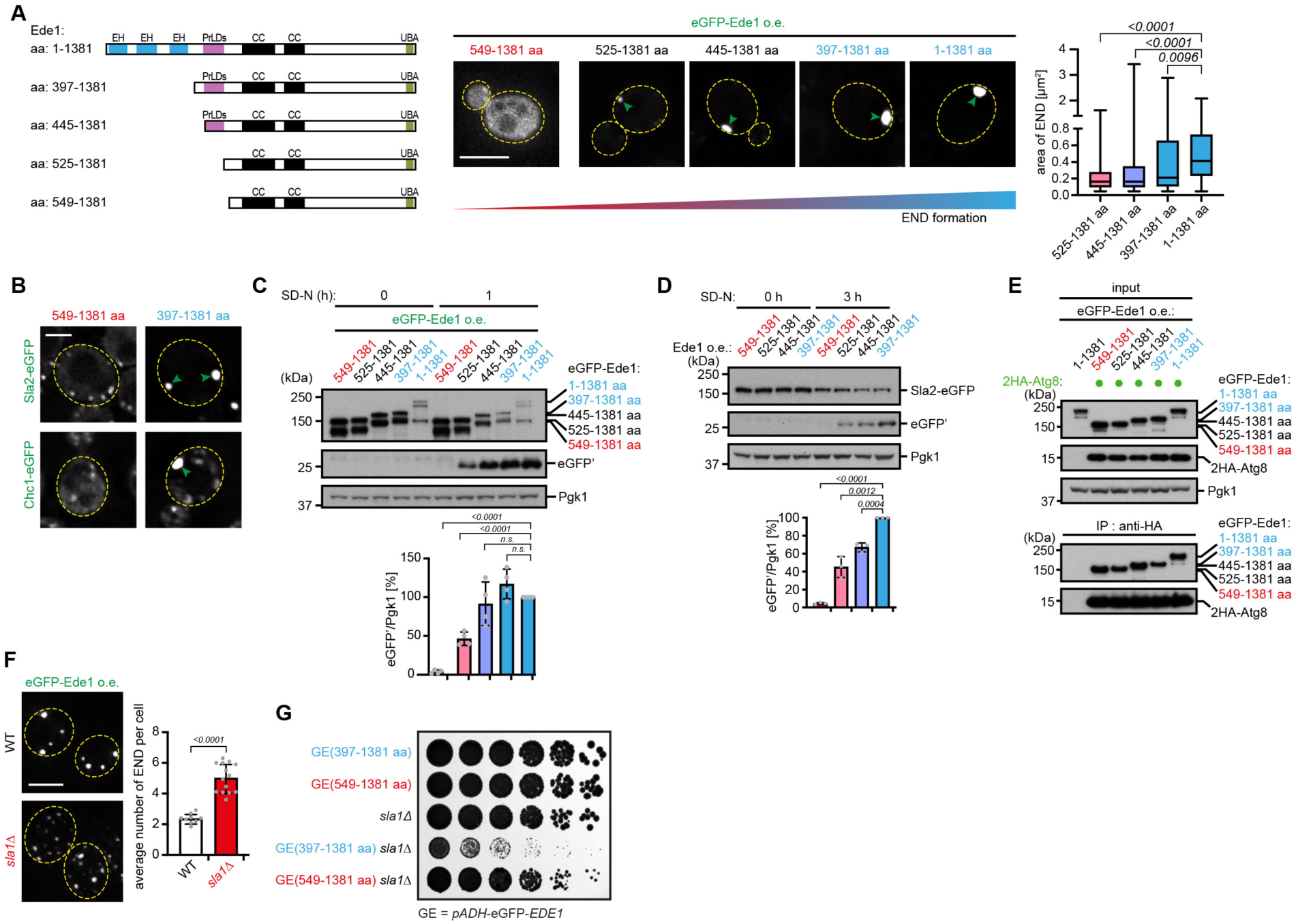
Phase Separation of Ede1 Is Essential for Autophagic END Degradation. **(A)** An *in silico* predicted prion-like domain (PrLD) in Ede1 is important for liquid-liquid phase separation. The PrLD was predicted using PrionW with a pWaltz cut-off of 70 (Zambrano et al., 2015). The different N-terminal truncations of Ede1 were expressed under the *ADH* promoter and formation of the END was monitored by fluorescence microscopy of the eGFP-Ede1 signal. Spots size was analyzed computational from maximum z-projections of the corresponding strains for a total of more than 500 spots of n = 3 biologically independent experiments. Statistical analysis was performed using two-tailed Student’s *t*-tests, *p*-values are indicated. Illustration shows N-terminal truncation mutants used to map the region important to undergo phase-separation. EH: Eps15-homology domain; PrLD: prion-like domain; CC: coiled-coil domain; UBA: ubiquitin-associated domain; aa: amino acid. Scale bar represents 5 µm. **(B)** END accumulation of the other CME proteins Sla2 and Chc1 depends on phase-separation of Ede1. Endogenously eGFP-tagged Sla2 or Chc1 accumulates in the END as visualized by fluorescence microscopy in the *ADH* promoter driven N-terminal EH truncation mutant of Ede1 (397-1381aa) but not in the PrLD-truncation mutant of Ede1 (549-1381aa). Scale bar represents 2 µm. **(C)** Autophagic degradation of Ede1 depends on its PrLD. Mutants expressing N-terminal eGFP-Ede1 truncations from an *ADH* promoter are analyzed by eGFP-cleavage assay before and after 1 hour of nitrogen starvation. Quantifications of free eGFP’ levels normalized to Pgk1 are shown. Pgk1 serves as a loading control. Data are mean ± s.d. of n = 4 biologically independent experiments. Statistical analysis was performed using two-tailed Student’s *t*-tests, *p*-values are indicated. **(D)** PrLD of Ede1 is necessary for autophagic degradation of other CME proteins accumulated in the END. Cells in which Sla2 are endogenously eGFP-tagged and co-expressing different N-terminal Ede1 truncation mutants from the *ADH* promoter are analyzed by eGFP-cleavage assay before and after 3 hours of nitrogen starvation. Quantifications of free eGFP’ levels normalized to Pgk1 are shown. Data are mean ± s.d. of n = 3 biologically independent experiments. Statistical analysis was performed using two-tailed Student’s *t*-tests, *p*-values are indicated. **(E)** Ede1 binding to Atg8 is independent of the presence of the PrLD. Atg8 expressed under the *ADH* promoter was 2HA-epitope-tagged, and its interaction with different N-terminal eGFP-tagged Ede1 truncation mutants (under the *ADH* promoter) was probed by coimmunoprecipitation and immunoblotting with anti-GFP antibody. **(F)** Deletion of the late coat protein Sla1 leads to END formation at multiple sites. END is monitored by overexpression of eGFP-Ede1 from the *ADH* promoter in WT and *sla1*Δ cells. Scale bar represents 5 µm. Quantification shows average number of END per cell derived from WT and *sla1Δ* mutant cells. Data are mean ± s.d. of n = 9 for WT and n = 14 for *sla1Δ* cells biologically independent experiments. Statistical analysis was performed using two-tailed Student’s *t*-tests, *p*-values are indicated. **(G)** Deletion of the PrLD domain in Ede1 rescues synthetic sickness with *sla1*Δ. Spotting analysis of different mutant strains, cells in log phase were spotted onto yeast YPD plates and incubated for 3 days at 30 °C. *See also Figure S7*.

With a mutant Ede1 defective in phase separation, we could test its impact on autophagic degradation. The different N-terminal Ede1 deletion constructs were analysed for autophagic turnover by GFP-cleavage assays. Autophagic degradation of Ede1 was strikingly correlated with its capacity to form condensates (Figure 6C). The cytosolic Ede1^549-1381^ mutant was no longer degraded in the vacuole, and degradation of Ede1^525-1381^, which showed only slight condensation, was substantially inhibited (Figure 6C). Similar dependencies were obtained for the degradation of the endocytic machinery proteins Chc1 and Sla2, which were degraded only if Ede1 was able to phase separate (Figures 6D and S7E). To rule out the possibility that these truncation mutants are defective in Atg8 binding, we monitored Atg8-Ede1 interaction of these by coIP experiments. Notably, for all mutants the binding to Atg8 was not affected, emphasizing that accumulation of Ede1 and other CME proteins is key to autophagic degradation (Figure 6E).

### Ede1 condensation is toxic when the transition from the early to the late phase is impaired

In all tested conditions, END assemblies coexist next to regular endocytic assemblies. To better understand what causes conversion from regular CME to END assemblies, we performed an unbiased synthetic interaction screen using cells overexpressing eGFP-Ede1 (Figure S7F), which identified a strong negative genetic interaction with the gene encoding Sla1 (Figures S7F-G). As the crucial function of Sla1 is linked to facilitate the transition of an endocytic patch from the early to the late phase (Carroll et al., 2012; Goode et al., 2015; Sun et al., 2015) and deletion of Sla1 extends the early phase of endocytosis (Sun et al., 2017), we speculated that Ede1 overexpression in the *sla1Δ* mutant might now affect endocytic assembly more globally. Indeed, fluorescence microscopy revealed that eGFP-Ede1 overexpression in *sla1Δ* cells led to an increase in number of END assemblies (Figure 6F). To test if the synthetic sickness resulted from the ability of Ede1 to phase separate, we introduced the Ede1^549-1381^ into *sla1Δ* background. In contrast to Ede1^397-1381^ *sla1Δ* mutant cells, Ede1^549-1381^ *sla1Δ* mutant cells showed an almost complete rescue of the phenotype, suggesting that indeed phase separation of Ede1 was cause of the problem in *sla1Δ* cells (Figure 6G).

### Visualizing END condensate degradation

Since LLPS compartments are increasingly described to undergo autophagic degradation, we sought to visualize how END condensates are degraded by autophagy. To examine the trajectory of the END condensate from the PM to an autophagosome, we conducted fluorescence microscopy experiments using a microfluidics device and fluorescence tracking of eGFP-Ede1 and mCherry-Atg8 in individual cells. After inducing nitrogen starvation to synchronize autophagic turnover, we observed clustering of mCherry-Atg8 specifically at sites of Ede1 accumulation. Small pieces pinched off of these structures into the cytosol in an Atg7-dependent manner and were positive for both marker proteins (Figure 7A and Movies S6-7), underlining that CME condensates are delivered in fractions to the vacuole.

**Figure 7.**
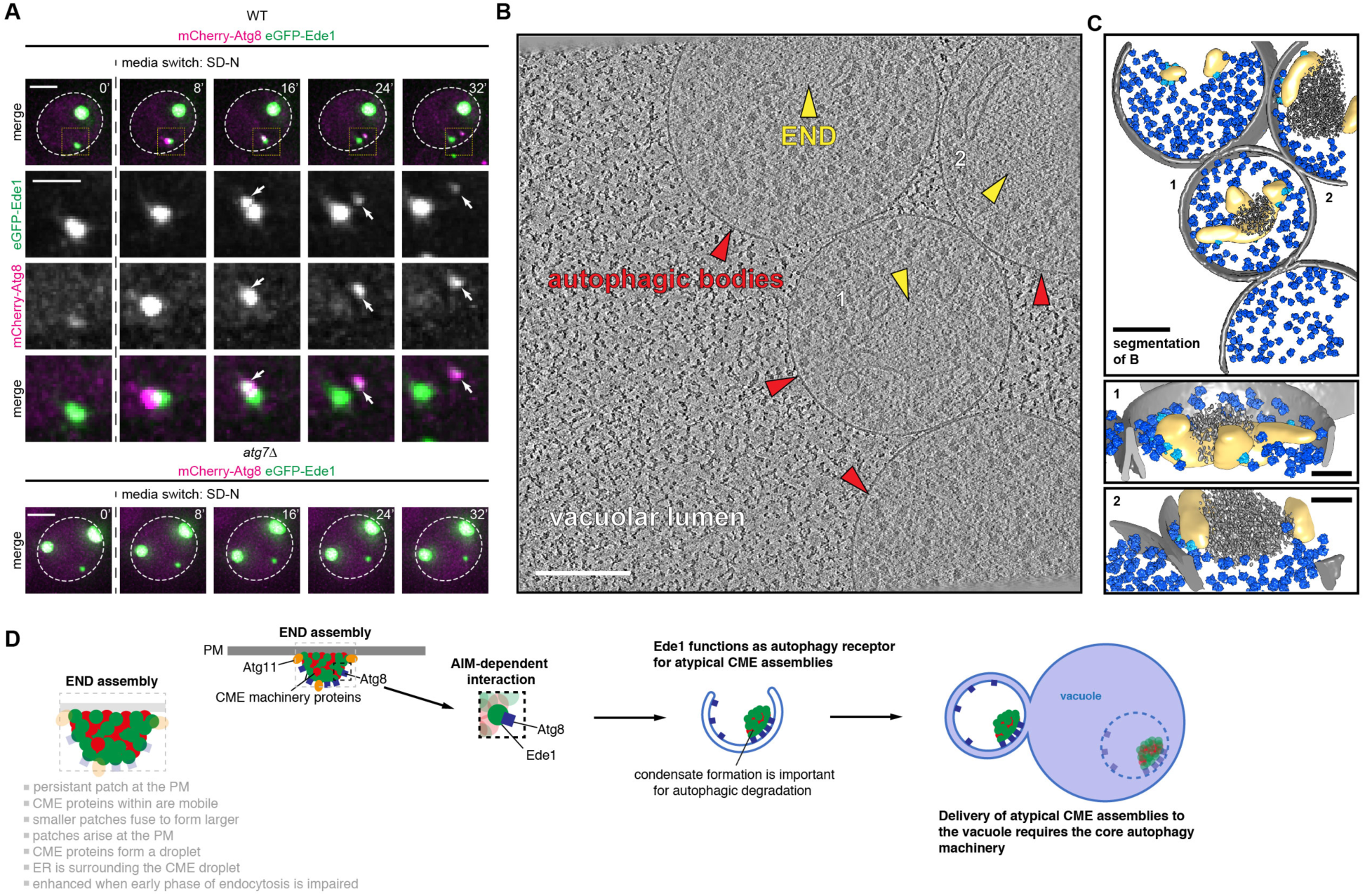
Visualization of Autophagic END Degradation. **(A)** Small portions of the END are engulfed by autophagy. Monitoring of wildtype or Atg7-deficient (*atg7*Δ) cells expressing eGFP-Ede1 and mCherry-Atg8 under the *ADH* promoter by fluorescence microscopy in a microfluidics device in 8 min intervals after switching to media lacking any nitrogen source. Maximum intensity z-projections are shown for the individual time points for both channels. Scale bar represents 1 µm. **(B)** The END accumulates together with ER in autophagic bodies. Ultra-structural analysis of ΔΔΔ cells expressing Ede1-eGFP by correlative cryo-ET in Atg15-deficient (*atg15*Δ) cells after nitrogen starvation. The example shows Ede1-eGFP protein accumulation together with fenestrated ER within autophagic bodies inside the vacuole. Shown is an average 2D section of the original tomogram. Scale bar represents 200 nm. **(C)** Segmentation of (B) shows END containing autophagic bodies. Ribosomes are shown in light (membrane attached) and dark blue (free); the autophagic body membrane is shown in light grey; drop-like protein accumulations are indicated in dark grey (by a density threshold) and membranes surrounding the protein accumulation are segmented in light yellow. Insets show close-ups for the END density corresponding to 1 and 2. Scale bar represents 200 nm and 100 nm for inset. **(D)** Model of END clearance by Ede1-dependent autophagy. Ede1 recruits Atg8 particularly to sites of atypical non-endocytic protein assembly, which we named END (short for: Ede1-dependent eNdocytic protein Deposit). END assemblies have distinct properties by liquid-liquid phase separation and their formation is enhanced when the early phase of endocytosis is disturbed. Interaction of Ede1 with the autophagic scaffold Atg11 and its requirement for END condensate degradation highlights this pathway as a branch of selective autophagy. The PrLD of Ede1 is essential for both phase separation and autophagic degradation of endocytic protein constituents of END assemblies.

To obtain snapshots of the END condensates after delivery to autophagic bodies, we turned to cryo-ET. Notably, lack of the vacuolar lipase Atg15 preserves autophagic bodies within the vacuole (Epple et al., 2001). In addition, our screen showed that deletion of the vacuolar lipase *ATG15* inhibits CME condensate degradation and led to accumulation of foci in the vacuole (Figure 4D). We thus performed our correlative cryo-ET workflow with the *ΔΔΔ atg15Δ* strain. Indeed, the fluorescent signal correlated to autophagic bodies within the lumen of the vacuole (Figure 7B). The structures inside these autophagic bodies were drop-like and similar to the ones observed before on the cytosolic side of the PM (Figure 5E), despite much smaller. Notably, ribosomes were likewise excluded from these condensates (Figure 7C and Movie S8). Interestingly, parts of the ER membrane were still attached to the CME condensates inside of the autophagic bodies (Figure 7C), raising the possibility of a role of the ER in preserving the condensate and/or in degradation. Overall, the data visualizes for the first-time phase condensates within autophagic bodies.

## Discussion

In this study, we define a new selective autophagy pathway, and its underlying molecular and cellular mechanisms, from the point of cargo recognition to delivery to autophagosomes. Starting with identification of CME proteins associated with Atg8, our data suggest that binding is mediated by a constituent of this complex machinery acting as a selective autophagy receptor, namely Ede1. Classical AIM-dependent interactions between Ede1 and Atg8 recruit proteins that normally mediate CME but phase-separate into atypical non-endocytic condensates – i.e. END assemblies - into autophagosomes. Interaction of Ede1 with Atg11 and its requirement for END condensate degradation highlights this pathway as a branch of selective autophagy. Moreover, we define an Ede1 PrLD as essential for both phase separation and autophagic degradation of endocytic protein constituents of END assemblies. Finally, the correlative cryo-ET experiments reveal for the first time the native environment of END assemblies at the PM and in autophagic bodies. On this basis, we propose a model in which defects in the early phase of CME protein assembly triggers phase separation via Ede1 PrLDs, while Ede1 is also associated with other endocytic proteins at the PM (Figure 7D). Ede1 may thus serve as an intrinsic member of the complex providing surveillance or at least serving as an effector for atypical or aberrant CME assemblies.

Ede1 thereby plays a dual role as initiator of CME and as determinant for clearance of defective CME protein “END” assemblies via its build-in Atg8 recruitment signal. We propose to call such proteins “intrinsic autophagy receptors”. Intrinsic autophagy receptors would be integral and functionally important subunits within normal complexes, but if needed also direct such complexes to autophagosomes for degradation. Thus, intrinsic autophagy receptors may provide an in-built quality control or other regulatory mechanism differing from conventional receptors that become dynamically recruited by some extrinsic signal for autophagic degradation. Intrinsic receptors would be ideally positioned to distinguish normal from aberrant protein complexes. Indeed, the translocon subunit Sec62 and the nucleoporin Nup159 are examples for such intrinsic receptors (Fumagalli et al., 2016; Lee et al., 2020a).

A remarkable feature of the END pathway is that the cargo shows characteristics of LLPS. By a combination of different approaches, we describe hallmarks of LLPS for aberrant CME assemblies: (1) Although Ede1 within the END assemblies is interchangeable with the cytosolic pool, the END condensate appears drop-like; (2) Photobleaching of a fraction of the END condensate leads to recovery through internal rearrangements; (3) END condensate growth is achieved by fusion/Oswald ripening of smaller assemblies. Remarkably, the drop-like shape of these assemblies is preserved at the PM and in autophagic bodies as visualized in our cryo-ET experiment. Central to the autophagic recognition of END assemblies is the ability of Ede1 itself to condensate. Deletion of the PrLD in Ede1 leads to loss of its ability to undergo LLPS and abolishes END condensate formation. Interestingly, although the PrLD truncated version of Ede1 is still able to bind Atg8, the degradation of both Ede1 and of the proteins normally found in the END condensates is highly impaired. This is in agreement with earlier observations that cargo recognition in autophagy pathways usually involves highly avid interactions between receptor and Atg8 (Zaffagnini and Martens, 2016). Moreover, it fits to the current concept that LLPS allows to establish these highly avid interactions to selectively enclose the cargo (Wang and Zhang, 2019). Notably, Ede1 does not need its UBA domain for phase separation and as such is clearly different from other receptors such as p62 which phase separates in the presence of polyubiquitin chains via the p62–ubiquitin interaction (Sun et al., 2018). Given that cells contain numerous membraneless compartments, we anticipate that there could be many other forms of selective autophagy, with remarkably distinct cargoes, awaiting discovery under different cellular stresses that increase need for cargo degradation, much like impairing endocytosis did here. Moreover, the concept of intrinsic receptors might be expandable for other proteins driving LLPS.

In case of endocytosis, it has been suggested that LLPS of endocytic proteins serves as a principal mechanism by which the complex assembly of CME proteins during endocytosis is achieved (Bergeron-Sandoval et al., 2018; Day et al., 2019; Kozak and Kaksonen, 2019). What restricts liquid-liquid phase separation of CME proteins during regular endocytosis and inhibits END formation? Notably, the PrLD domain of Ede1 that we identified as being indispensable for LLPS and autophagic degradation contains thirteen experimentally validated phosphorylation sites. Introduction of negatively charged phosphoserine, phosphothreonine, or both into the Ede1 PrLD would likely increase electrostatic repulsion between PrLDs and might restrict self-assembly. Such a concept would be analogous to the LLPS protein FUS, where phosphorylation in the PrLD disrupts phase separation (Monahan et al., 2017). In contrast, genetic perturbations of the early phase of endocytosis enhances formation of END assemblies. While we do not know all the functional consequences of these mutants, the major phenotype appears to be improper clathrin assembly (Boettner et al., 2011; Weinberg and Drubin, 2012). Taken together, we speculate that CME proteins that are not involved in CME, either due to subunit excess, defects, imperfect engagement with CME assemblies, or other defects may provoke atypical assembly into phase-separated condensates with Ede1. Importantly, the ability of Eps15, the human homolog of Ede1, to phase-separate (Day et al., 2019) and to bind LC3 (mammalian Atg8 homologue) (Bejarano et al., 2012) is conserved, raising the possibility that this END autophagy pathway could also be evolutionary conserved. Notably, genetic perturbation of the early phase of endocytosis in human cells results in atypical accumulation of clathrin, Eps15 and AP2 at the PM (Kirchhausen et al., 2014; Mettlen et al., 2018; Meyerholz et al., 2005). Therefore, we anticipate that features we showed regulating END condensates in *S. cerevisiae* most likely occur in mammalian cells too.

Finally, our ultrastructural dissection of the END at the plasma membrane and within autophagic bodies provides first insights into the architecture of these condensates. Notably, we observed the presence of fenestrated ER around the CME condensate, which is intriguing, since the ER is involved in autophagosome biogenesis (Hurley and Young, 2017; Mizushima et al., 2011; Noda and Inagaki, 2015; Shibutani and Yoshimori, 2014). However, parts of the ER appear to be degraded along with the endocytic proteins. It is therefore appealing to speculate a twofold involvement of ER - first, by providing a membrane source for autophagosome biogenesis (Maeda et al., 2019; Osawa et al., 2019; Schutter et al., 2020; Valverde et al., 2019) and second, by regulating the dynamics of the END condensate, a concept recently established for processing bodies (Lee et al., 2020b). Importantly, our ultrastructural dissection provides the first view how LLPS condensates can be packed into autophagosomes, with ER at the periphery, and future experiments are needed to determine the role of the ER in this process. It is noteworthy that autophagy initiation also depends on Atg1 signalling via a membrane-associated phase separated droplet organizing Atg proteins at the PAS (Fujioka et al., 2020). It seems likely that future studies will reveal other networks between membranes and phase separated proteins in autophagy. A structural platform is now provided by visualization of such interactions spanning from the cytoplasm to autophagosomes.

## Acknowledgments

We thank S. Schkoelziger, S. Kienle and L. Rieger for technical assistance, J. Rech for help with pepscan analysis and robot-based techniques, the MPIB Imaging Facility, in particular G. Cardone and M. Spitaler for help with image analysis, N. Nagaraj and S. Uebel for mass spectrometry and protein analysis, D. Scott and S. Jayaraman for help with crystallography. Thomas Wollert and Scott Emr for reagents, Jürgen Plitzko and Miroslava Schaffer for microscope support, Ulrich Hartl and Manajit Hayer-Hartl for discussions, Anna Bieber and Cristina Capitanio for the critical reading of the manuscript. This work is supported by the Max Planck Society (to S.J, B.P., B.A.S. (SCHU 3196/1-1) and W.B.), Deutsche Forschungsgemeinschaft (DFG, German Research Foundation) (to S.J., B.P., and W.B.), Center for Integrated Protein Science Munich (to S.J.), ERC Advanced Grant (ERC-2013-AdG-339176) (to S.J.), and the Louis-Jeantet Foundation (to S.J.). F.W. was supported by an EMBO Long-term Fellowship ALTF 764-2014. P.S.E. was supported by an Alexander von Humboldt returners fellowship. Data were collected at Southeast Regional Collaborative Access Team (SER-CAT) 22-ID (or 22-BM) beamline at the Advanced Photon Source, Argonne National Laboratory. SER-CAT is supported by its member institutions (see www.ser-cat.org/members.html), and equipment grants (S10_RR25528 and S10_RR028976) from the National Institutes of Health.

## Authors Contributions

S.J. and F.W. designed the study. F.W., P.E., S.J. and B.A.S. designed experiments and analyzed data. F.W., C.W.L., Y.Z. and P.E. conducted the experiments. F.W., B.P., P.E., B.A.S. and W.B. wrote the manuscript. All authors discussed the results and commented on the manuscript.

## Methods

### Yeast cell culture, starvation, eGFP cleavage assay, spotting assay and cloning

All strains used in this study are derivatives of *Saccharomyces cerevisiae* and are listed in Tables S3 and S4, respectively. Standard protocols for transformation, mating, sporulation, and tetrad dissection were used for yeast manipulations (Dunham et al., 2015). Yeast cultures were inoculated from overnight cultures, grown using standard growth conditions and media (Sherman, 2002). If not indicated otherwise, cells were cultured at 30 °C in YPD-medium (1% yeast extract, 2% peptone and 2% glucose). For eGFP cleavage assay, cells were grown to mid-log phase (optical density at 600 nm (OD_600_) of 1.0) then switched into SD-N medium (synthetic minimal medium lacking nitrogen; 0.17% YNB without amino acids and ammonium sulfate, supplemented with 2% glucose) and incubated for the indicated time. Alternatively, cells were treated with 100 nM rapamycin for the indicated times. Mitophagy was examined as described previously (Tal et al., 2007). Autophagy substrates were tagged at the endogenous, chromosomal location with eGFP and their vacuolar degradation following starvation was monitored by accumulation of the released eGFP moiety, which is largely resistant to vacuolar degradation. For spotting growth assay, 5 x 10^4^ cells from overnight cultures were 5-fold serially diluted and dropped onto YPD plates. Plates were scanned after an incubation for 3 days at 30 °C. Chromosomally tagged strains and knockout strains were constructed by a PCR-based integration strategy (Knop et al., 1999). Standard cloning and site-directed mutagenesis techniques were used.

### Immunoblot techniques

Yeast protein extraction was performed as described before (Cheong and Klionsky, 2008). The TCA (Trichloroacetic acid; to a final concentration of 10% for 20 min on ice) precipitated cell pellet was resuspended in 100 µl MURB buffer (50 mM Na_2_HPO_4_, 25 mM 2-[N-morpholino]ethanesulfonicacid (MES), pH 7.0, 1% SDS, 3 M urea, 0.5% 2- mercaptoethanol, 1 mM NaN_3_ and 0.05% bromphenol blue) and disrupted by vortexing with an equal volume of acid-washed glass beads for 5 min. Followed by an incubation at 70 °C (1400 rpm) for 10 min. Proteins were separated by NuPAGE 4%–12%gradient gels (Invitrogen), transferred onto PVDF membranes (Immobilon^®^-P) and then analysed using specific antibodies (see section “Antibodies”). For quantification, proteins were transferred onto low fluorescence PVDF membranes (Immobilon^®^-FL), and analysed using specific antibodies followed by fluorescent IRDye^®^ secondary antibodies on an Odyssey^®^ Fc imager (LI-COR).

### Coimmunoprecipitation

Yeast cell lysates from 200 OD_600_ were prepared by cell disruption on a multitube bead-beater (MM301 from Retsch GmbH) in lysis buffer (100 mM Hepes pH 7.4, 150 mM NaCl, 1% NP-40, 10% glycerol, 50 mM NaF, 2 mM phenylmethylsulfonyl fluoride (PMSF), and EDTA-free protease inhibitor cocktail (cOmplete Tablets, Roche)) with zirconia/silica beads. The extracts were cleared by centrifugation at 8000 g for 10 min and supernatants were incubated with GFP-Trap_A matrix (ChromoTek) or anti-HA magnetic beads (Thermo Scientific) for 2 h with head-over-tail rotation at 4 °C and followed by 5 times washing steps with lysis buffer to remove nonspecific background binding. Bound proteins were eluted by adding HU loading buffer and incubation at 65 °C for 10 min.

### Mass spectrometry and data analysis

To analyze the Atg8 interactome for the different yeast strains, four biological replicates of cells were grown at 30 °C in 200 ml YPD medium each. At OD 1.0, cells were incubated for 3 h with 100 nM rapamycin. Yeast cells were subsequently collected by centrifugation and lysates were prepared by cell disruption on a multitube bead-beater (MM301 from Retsch GmbH) in lysis buffer (20 mM Hepes pH 7.5, 150 mM KOAc, 1% NP-40, 5% glycerol, 10 mM N-Ethylmaleimide, 1 mg/ml Pefabloc SC (Roche), and EDTA-free protease inhibitor cocktail (cOmplete Tablets, Roche)) with zirconia/silica beads. The extracts were cleared by centrifugation at 2000 g for 10 min. Supernatants were incubated with GFP-Trap_M or GFP-Trap_MA matrix (ChromoTek GmbH) for 1 h on a rotary wheel at 4 °C. Magnetic beads were washed 2 times with lysis buffer and 4 times with washing buffer (50 mM Tris pH 7.5, 150 mM NaCl, 1 mg/ml Pefabloc SC (Roche), and EDTA-free protease inhibitor cocktail (cOmplete Tablets, Roche)) to remove any residual detergent. The supernatants from the beads were removed and the beads were incubated with a buffer containing 4 M urea and 20 mM DTT in 25 mM Tris pH 8.0 buffer for 10 minutes followed by incubation with 40 mM chloroacetamide for 20 minutes for alkylation of cysteines. The sample was diluted to final concentration of 1 M urea with digestion buffer (25 mM Tris pH 8.0) and vortexed. The sample was digested for 2 h with 0.5 µg of endoproteinase lysine-C (Wako chemicals) and then digested with 0.5 µg of trypsin (Promega) overnight. The digested peptides were purified using StageTip (Rappsilber et al., 2003). Peptides were loaded on a 15 cm column (inner diameter = 75 microns) packed with C18 reprosil three micron beads (Dr Maisch GmbH) and directly sprayed into a LTQ-Orbitrap XL instrument operated in a data-dependent fashion. Up to top five precursors were selected for fragmentation by CID and analyzed in the iontrap. The raw data were processed using MaxQuant (Cox and Mann, 2008) version 1.6.0.15. Peak lists generated were searched against a yeast ORF database using Andromeda search engine (Cox and Mann, 2008) built into Maxquant. Proteins were quantified using the MaxLFQ algorithm (Cox et al., 2014). Analysis was performed using Perseus (Tyanova et al., 2016) version 1.5.4.2 as described before (Hubner et al., 2010). The mass spectrometry proteomics data have been deposited to the ProteomeXchange Consortium via the PRIDE (Perez-Riverol et al., 2019) partner repository.

### Protein sample production for crystallography

*S. cerevisiae* Atg8 (1-116) with stabilizing mutation K26P was expressed and purified following the same production protocol for human homolog LC3 (Zheng et al., 2017b). Atg8 protein was mixed with HPLC-purified synthetic Ede1 (1220-1247) peptide at a molar ratio of 1:2, followed by a 5-minute incubation at 4 °C. The complex was purified by size-exclusion chromatography using Superdex75 10/300 GL (Sigma) column in buffer containing 20 mM HEPES (pH 7.0), 100 mM NaCl, and 5 mM DTT. The peak fractions were concentrated to 33 mg/mL as the final sample.

### Crystallization of Ede1-Atg8 complex

Crystals were obtained using vapor diffusion method with macro seeding procedure for improvement. 1 μL of protein was mixed with 1 μL of well buffer containing 1.3 M ammonium sulfate and 0.1 M sodium acetate (pH 4.8), and was allowed to grow in hanging drops at 4 °C for 24 hours. The resulting crystals were crushed and diluted in the same well buffer to be used as seeds. Final crystals were obtained by mixing 0.2 µL seeds, 1 µL of protein sample, and 1 µL of well buffer containing 1.1 M ammonium sulfate, 0.1 M sodium acetate (pH 4.8), and 0.01 M sodium iodide after growth at 4 °C for 48 - 72 hours in hanging drops before harvest. Crystals were cryo-protected in 25% glycerol in addition to the mother liquor, and were flash-frozen in liquid nitrogen 48 hours prior to the data collection.

### Data collection, processing, refinement and deposition

Data for Ede1-Atg8 complex was collected at Advanced Photon Source (APS) SER-CAT beamline 22-BM. Use of the Advanced Photon Source was supported by the U. S. Department of Energy, Office of Science, Office of Basic Energy Sciences, under Contract No. W-31-109-Eng-38. The dataset was created from a single crystal, and was processed and scaled by HKL2000. Phase was solved by molecular replacement using Phaser in CCP4, adopting Atg19-Atg8 complex structure (PDB 2ZNP) as search model. Model was built in Coot. And refinements were primarily performed by Refmac in CCP4, and finalized by Refmac in Phenix. The coordinate and structure factors for Ede1-Atg8 were deposited to the RCSB Protein Data Bank.

### Expression and purification of GST-fusion proteins and His-Atg8

GST-fusion proteins and His-Atg8 were expressed in E. coli Rosetta^TM^ 2(DE3) (for GST-fusion proteins) or Rosetta^TM^ 2(DE3)pLysS (for His-Atg8) cells, respectively. Expression was induced with 1 mM IPTG in a 1 L culture of LB (lysogeny broth) for 20 h at 22 °C. Cells were harvested by centrifugation and lysed in lysis buffer (GST-fusion proteins: 40 mM Tris pH 7.5, 150 mM NaCl, 5 mM DTT, EDTA-free protease inhibitors cocktail (cOmplete Tablets, Roche), 1 mg/ml Pefabloc SC (Roche); His-Atg8: 40 mM Tris pH 7.5, 500 mM NaCl, 5 mM MgCl_2_, 5 mM 2-Mercaptoethanol, 20 mM Imidazole, 10% glycerol (w/v), EDTA-free protease inhibitors cocktail (cOmplete Tablets, Roche), 1 mg/ml Pefabloc SC (Roche)) using an EmulsiFlex C3 homogenizer (Avestin). DNA was digested using SM DNase (final 75 U/ml, 15 min on ice). Supernatant containing soluble proteins was collected by centrifugation (20000 rpm, 30 min). GST-fusion proteins and His-Atg8 were affinity purified using Glutathione Sepharose 4 Fast Flow (GE Healthcare) or Ni-NTA agarose (Qiagen) respectively (2.5 h on a rotary wheel at 4 °C). The resins were recovered by gravity-flow chromatography. Resins were subsequently washed 3 times with 25 ml lysis buffer, followed by 3 times with washing buffer (GST-fusion proteins: 40 mM Tris pH 7.5, 450 mM NaCl, 5 mM DTT; His-Atg8: 40 mM Tris pH 7.5, 500 mM NaCl, 5 mM MgCl_2_, 5 mM 2- Mercaptoethanol, 70 mM Imidazole, 10% glycerol (w/v)). Bound proteins were eluted from the individual resin (GST-fusion proteins: 40 mM Tris pH 7.5, 150 mM NaCl, 5 mM DTT, 50 mM glutathione; His-Atg8: 40 mM Tris pH7.5, 500 mM NaCl, 5 mM MgCl_2_, 5 mM 2- Mercaptoethanol, 270 mM Imidazole, 10% glycerol (w/v)). The eluted proteins were collected and dialyzed (50 mM Tris pH 7.5, 150 mM NaCl, 20% glycerol (w/v), overnight 4 °C). Purified proteins were directly frozen after dialysis and stored in aliquots at −80 °C until further use. The identity of the different proteins was confirmed by SDS-PAGE. Protein concentration was determined by Pierce^TM^ BCA protein assay kit.

### Atg8 *in vitro* binding assay

The *in vitro* binding assay was performed by incubation of the indicated protein combinations in 1 ml assay buffer (50 mM Tris pH 7.5, 150 mM NaCl, 5% glycerol (w/v), 20 mM Imidazol, 0.1% Triton X-100) for 1h at room temperature. 50 µl were used as input control and mixed with an equal amount of HU loading buffer (8 M Urea, 5% SDS, 200 mM Tris-HCl pH 6.8, 20 mM dithiothreitol (DTT), and Bromophenol blue 1.5 mM). The rest of the supernatant was added to 100 µl Ni-NTA agarose (Qiagen) slurry and incubated for 2.5 hours at 4 °C on a rotary wheel. The resin was collected by centrifugation (800 rpm, 1 min) and washed 6 times with 1 ml washing buffer (50 mM Tris pH 7.5, 150 mM NaCl, 5% glycerol (w/v), 20 mM Imidazol, 1% Triton X-100). After the last washing step, the supernatant was removed (27G needle) and the proteins were eluted with 50 µl of elution buffer (50 mM Tris pH 7.5, 150 mM NaCl, 5% glycerol (w/v), 270 mM Imidazol, 1% Triton X-100). The eluate was transferred into a new tube (27G needle), mixed with an equal amount of HU loading buffer and analyzed by SDS-PAGE. Proteins separated by SDS-PAGE were stained with PageBlue Protein Staining Solution (Thermo Scientific).

### GST-Atg8 pulldown with yeast cell extract

The indicated GST-fusion proteins were incubated (3 nM) diluted in 1 ml assay buffer (50 mM Tris pH 7.5, 150 mM NaCl, 5% glycerol (w/v, 0.1% Triton X-100) and mixed with 50 µl Glutathione Sepharose 4 Fast Flow (GE Healthcare) slurry. The mixture was incubated for 2.5 hours at 4 °C on a rotary wheel. Meanwhile, the yeast lysate was prepared as described above (mass spectrometry, 20 µl were kept as input control and mixed with 200 µl HU loading buffer). The Glutathione Sepharose was collected by centrifugation (800 rpm, 1 min) and washed one time with washing buffer (50 mM Tris pH 7.5, 150 mM NaCl, 5% glycerol (w/v), 1% Triton X-100) followed by one time with the yeast lysis buffer (20 mM Hepes pH 7.5, 150 mM KOAc, 1% NP-40, 5% glycerol, 1 mg/ml Pefabloc SC (Roche), and EDTA-free protease inhibitor cocktail (cOmplete Tablets, Roche)). Subsequently, 300 µl of the yeast lysate was mixed with 700 µl yeast lysis buffer and added to the resin (2.5 h, 4 °C rotary wheel). The resin was washed three times with yeast lysis buffer and the supernatant was in the final step removed with a 27G needle. The bound proteins were denatured by addition of 50 µl HU loading buffer and incubation at 65 °C for 10 min.

### Fluorescence microscopy

For fluorescence microscopy, yeast cells were grown in low fluorescence synthetic growth medium (yeast nitrogen base without amino acids and without folic acid and riboflavin (FORMEDIUM)) supplemented with all essential amino acids and 2% glucose. The next day cells were diluted to OD 0.1 and grown till mid log phase (0.5 – 0.8 OD) before imaged. Microscopy slides were pre-treated with 1 mg/ml Concanavalin A (ConA) solution. 7- Amino-4-Chlormethylcumarin (CMAC) was part of the Yeast Vacuole Marker Sampler Kit (Y7531) from Thermo Fisher and staining was performed according to manufacturer recommendations. The widefield imaging was performed at the Imaging Facility of the Max Planck Institute of Biochemistry (MPIB-IF) on a GE DeltaVision Elite system based on an OLYMPUS IX-71 inverted microscope, an OLYMPUS (100X/1.40 UPLSAPO and 60X/1.42 PLAPON) objective and a PCO sCMOS 5.5 camera. Images were deconvolved using the softWoRx^®^ Software (default values except: method: additive enhanced, 20 iterations). Image analysis was performed using ImageJ/Fiji (Schindelin et al., 2012; Schneider et al., 2012).

### FRAP and FLIP analysis

For FRAP analysis 131 frames were collected with a time interval of 1 s. 10 frames were collected before bleaching with a high laser power (point bleaching) for 1 frame. 120 frames were collected after bleaching. Double normalization was performed as described before (Phair et al., 2004). Fitting of normalized FRAP curves was performed using a single exponential equation with the software Origin (version 2019b). FLIP analysis was performed as described before (Snapp, 2013).

### Yeast spot analysis

Quantification of fluorescent spots in yeast cells was performed using the software ImageJ/Fiji (Schindelin et al., 2012; Schneider et al., 2012), complemented with the plugins provided by the ImageScience update site. Cells in each image were identified with YeastSpotter (Lu et al., 2019), which was installed locally and controlled through Fiji. The workflow was implemented as a series of actions in an ImageJ Toolset, each action performing a processing step on all the images in given directory. As a first step, the stacks of fluorescent images acquired were compressed by maximum intensity projection. YeastSpotter was then executed on DIC (Differential Inference Contrast) images from the same field of view, to generate a mask identifying position and size of cells, to apply to the corresponding fluorescence image. In order to eliminate possible detection errors, the candidate cells were screened for diameter (larger than 4 µm) and aspect ratio (smaller than 5). Each cell was analyzed independently, with their spots being first enhanced with a Laplacian of Gaussian operator (FeatureJ: Laplacian plugin, smoothing scale 63 nm) and then detected by thresholding. The threshold was set adaptively to 9 times the standard deviation above the average intensity of the cell. In order to avoid that the average intensity was biased by the strength and number of spots, this one was calculated after excluding all the pixels with intensity 3 times the standard deviation above an initial estimate from all the pixels in the cell.

### Microfluidics experiments

Microfluidics experiments were performed on a CellASIC^®^ ONIX microfluidic platform (Merck) with the CellASIC^®^ ONIX microfluidic plates for haploid yeast cells. Cells were loaded from a logarithmic growing culture (OD 0.5 – 0.8) according to manufacturer’s manual and imaged in 8 min intervals over a time course of 8 h. At each position for each time point a z-stack was recorded (300 nm). Imaging was performed with a constant media flow rate of 1 psi. To rapidly switch media, flow rate was set to 5 psi for 5 min and then switched back to a flow rate of 1 psi. Cells were initially cultured in low fluorescence synthetic growth medium and then switched to SD-N medium. The microscope used was the same as described in the Fluorescence microscopy section.

### Automated yeast library manipulations and high-throughput microscopy

SGA and microscopic screening were performed using an automated microscopy set-up as previously described (Collins et al., 2010), using the Biomek FXP (Beckman Coulter) and an Opera Phenix high content screening system (PerkinElmer). The yeast knockout library was from Thermo Fisher. Images were acquired using a 60X water lens with excitation at 490/20 nm and emission at 535/50 nm (GFP). After acquisition, images were analysed and manually reviewed using the Harmonie 4.8 software for the visual analysis of END compartment or Cellprofiler 2.2 and Matlab 2018a for analysis of colony growth. Each strain was generated in duplicates. For the visual screen 4 images per strain were acquired. Images were analyzed by defining the number of END spots per image divided by the total cell area. A z-score for each individual value against the screen mean was computed. Table S1 shows hits with a z-score higher then 1.5.

### Electron microscopy

#### Cryo-EM sample preparation

For cryo-ET experiments, yeast cultures were inoculated from overnight cultures and grown in YPD media at 30 °C to an OD of 0.6. To allow for later 3D-correlation, cells were supplemented with fiducial markers (Dynabeads MyOne, Thermo Fisher Scientific) at 1:16 dilution. The cell suspension containing fiducials (4 µL) was applied to holey carbon R2/1 copper, or holey SiO_2_ R1/4 TEM grids (Quantifoil) to allow for more robust correlation and cells were plunge frozen on a Vitrobot Mark IV (FEI, settings: blotforce = 10, blottime = 10 s, temperature = 30 °C, humidity = 90%). Samples were stored under LN_2_ until use. Grids containing vitrified yeast cells were clipped in modified Autogrids, and fluorescent image stacks (typically 70 slices at 300 nm steps) were recorded in confocal mode at selected squares using a 40x long distance air objective (Zeiss EC Plan-Neofluar 40x/0.9 NA Pol) on a fluorescence microscope equipped with a cryo module (Corrsight, FEI) using the 488 laser channel. Potential sites of Ede1 accumulation were identified in the FLM stacks and correlated using custom software (Arnold et al., 2016) with the SEM/IB images of a dual beam focused ion beam microscope (FIB Quanta 3D FEG, FEI) equipped with a Quorum PP3000T cryo-system (Quorum Technologies, Laughton, United Kingdom) and a homemade 360° cryo-stage cooled by an open nitrogen circuit (Rigort et al., 2010). Lamellas were cut at correlated sites analogous to published protocols (Schaffer et al., 2017).

#### Cryo-EM data acquisition

Cryo-tomograms were acquired on a transmission electron microscope (Titan Krios, FEG 300 kV, FEI) with a post-column energy-filter (968 Quantum K2, Gatan) with a defocus range of −5 μm to −3.5 µm and an EFTEM magnification of 42000x (calibrated pixel size 3.42 Å). Images were recorded with a direct detection camera (K2 Summit, Gatan) in dose-fractionation mode and a total dose of ∼ 140 e^-^ / Å^2^ per tomogram using the SerialEM software package (Mastronarde, 2005). The acquisition was controlled by an in-house script running a dose-symmetric tilt scheme (Hagen et al., 2017) with an angle increment of 2° between the range of 70° and −50° starting at 10° to compensate for the lamella pre-tilt (∼ 12°). Frames were aligned using motioncorr2 (Zheng et al., 2017a), and tilt-series alignment as well as tomogram reconstruction was performed in IMOD (Kremer et al., 1996).

#### Cryo-EM template matching and subtomogram averaging

Ribosome positions were determined by template matching on 2x binned tomograms (IMOD bin 4, 13.68 Å pixel size) using the PyTom(Hrabe et al., 2012) software package. A reference was constructed from ∼300 manually picked ribosomes, which were aligned in PyTom using a fast-rotational matching (FRM) algorithm. The reference was further truncated at the small subunit to allow removal of false positives. For each tomogram, the 600 highest scoring cross-correlation peaks were extracted, and the subtomograms aligned (FRM) and classified using constrained principal component analysis (CPCA) to remove false positives as judged by the absent small subunit in the class averages. After initial processing in PyTom, further classification and refinement of the unbinned subtomogram averages was performed in Relion 2.1 (Scheres, 2012), including normalization and CTF estimation with CTFFIND 4.1.5 (Rohou and Grigorieff, 2015). The final list of particles (∼11000) yielded a ribosome structure at 16 Å resolution (0.143 FSC criterion), which was used in all animations with appropriate binning.

#### Visualization

To enhance the contrast of TEM insets, the sum projections of 5 slices each around a central slice are shown. For 3D visualization, membranes were manually segmented in AMIRA 6.2 (FEI) and subtomogram averages of ribosomes (2x binned) were placed at positions derived from template-matching and cleaned by classification using custom MATLAB (Mathworks) scripts. Static scenes and videos were rendered in Chimera (Pettersen et al., 2004) at 13.68 Å pixel size (bin 2x).

### Antibodies

Monoclonal antibodies against HA-epitope (clone F-7) and Ubiquitin (P4D1) were purchased from Santa Cruz Biotechnology, Dpm1 (clone 5C5A7) and Pgk1 antibodies (clone 22C5D8) were from Invitrogen. The monoclonal GFP antibody (clones 7.1 and 13.1) was purchased from Sigma-Aldrich (former Roche). Antibodies against Atg8 and Ede1 were raised in rabbits against a recombinant Ede1 fragment (amino acids 1-400) or recombinant full length Atg8 expressed and isolated from *E. coli*. The individual serum was affinity purified to enrich for the antigen specific antibodies. Anti-Ape1 serum was a gift from Thomas Wollert.

### Quantification and statistical analysis

Error bars represent SD as indicated in the figure legends. Data were processed in Prism. Statistical analysis of differences between two groups was performed using a two-tailed, unpaired t test;

## Supplement Figures

**Figure S1.**
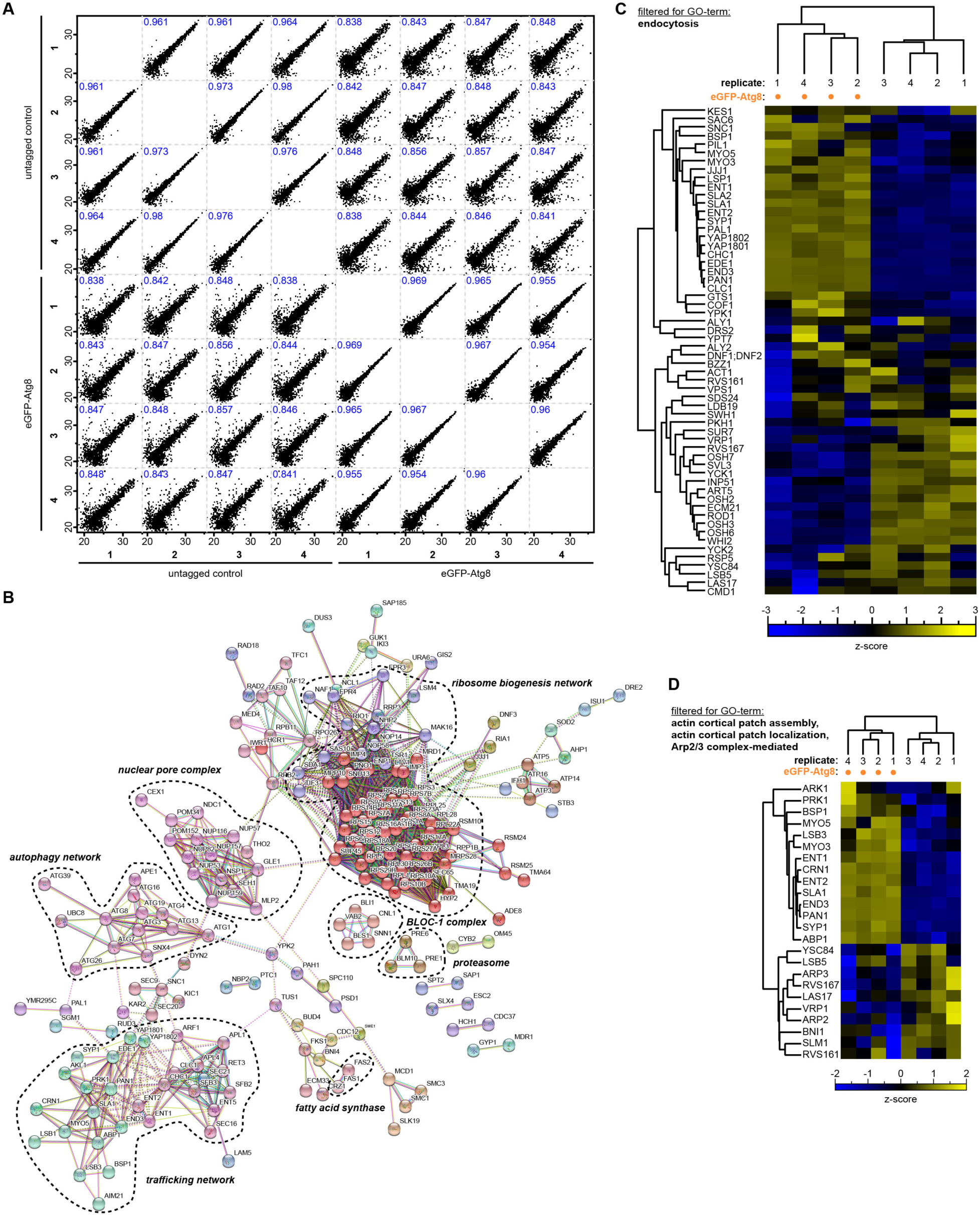
qMS Identified Components of Endocytic Machinery as Novel Atg8 Interactors, **Related to Figure 1**. **(A)** Multi scatter plot comparing all samples in relation to each other. The Pearson coefficient is indicated in each box with blue. **(B)** STRING analysis of protein-protein interaction networks of the 225 specific eGFP-Atg8 interactors reveals binding of distinct complexes. Only protein with at least one connection. are shown. **(C-D)** Hierarchical cluster analysis of Atg8 interactors filtered for GO-terms endocytosis or actin machinery. All eGFP-Atg8 specific interactors were filtered for GO-terms endocytosis (D) or Arp2/3 complex mediated actin nucleation, actin cortical patch assembly and actin cortical patch localization (E). Matching ANOVA-positive and FDR corrected hits are displayed by hierarchical clustering of the respective z-scores for each sample.

**Figure S2.**
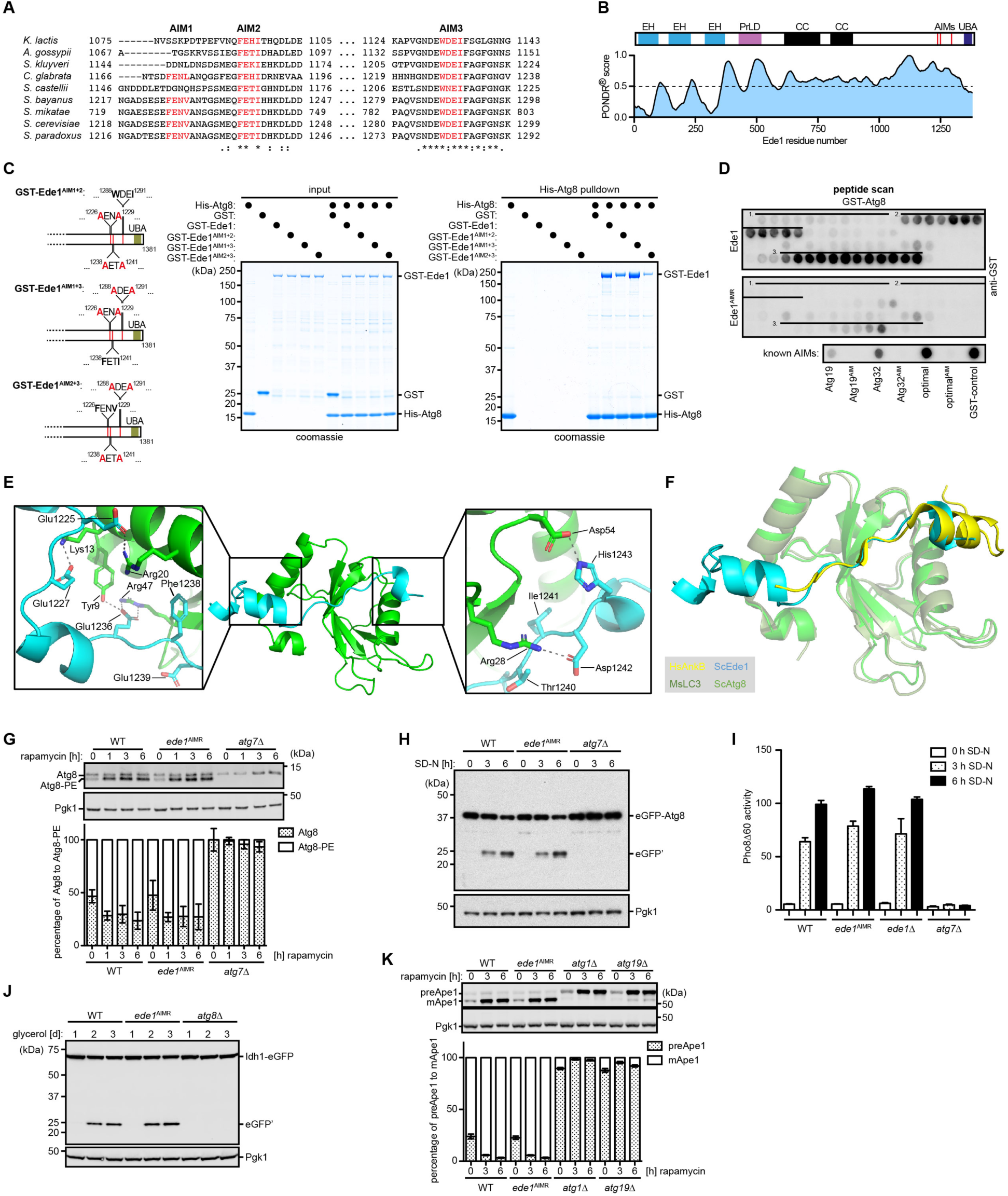
Autophagy Pathways Are Unaffected in *ede1*^AIMR^ Mutant Strain, Related to Figures 2 and 3. **(A)** AIM2 and 3 are conserved among different yeast species as shown by multiple sequence alignment (https://www.ebi.ac.uk/Tools/msa/clustalo/). **(B)** AIMs are located in the intrinsic disordered C-terminal region of Ede1. Output of intrinsic disorder prediction program PONDR (http://www.pondr.com) for Ede1, with schematic views of Ede1 structural elements. EH: Eps15-homology domain; PrLD: prion-like domain; CC: coiled-coil domain; UBA: ubiquitin-associated domain. **(C)** The binding between Ede1 and Atg8 is mainly mediated by AIM2 and 3 *in vitro*. Ni-pulldown assays were used to enrich for Ede1-Atg8 complexes, after recombinant His-tagged Atg8 (His-Atg8, 8 nM) was incubated with different double AIM mutants of GST-tagged Ede1 (Ede1^AIM1+2^, Ede1^AIM1+3^ or Ede1^AIM2+3^; 0.8 nM each). Graph shows position of mutants used to map the AIMs. UBA: ubiquitin-associated domain. **(D)** Peptide scanning analysis confirms the location of the three Atg8-intertacting motifs (AIM). A membrane-bound array of 176 overlapping peptides of 15 aa, with an offset of 1 aa, covering the entire C-terminal sequence of Ede1, was analyzed for binding to GST-Atg8 by immunoblotting with GST-specific antibodies. As control the same array was synthesized with the important aa in the respective AIM mutated. **(E)** Ede1 (1220-1247) peptide renders extended interactions with Atg8 additional to AIM binding. The acidic sidechains of Ede1 pre-AIM sequences form massive polar bonds with the sidechains in the Atg8 N-terminal helix (left). Ede1 post-AIM residues also contribute additional polar contacts to Atg8 (right). **(F)** The binding pattern of Ede1 peptide to Atg8 resembles that of ankyrin-derived peptides to Atg8 homologs in mouse (Li et al., 2018). To show an example, a complex of HsAnkB-MsLC3B (PDB No. 5YIS) is superimposed to Ede1-Atg8 (conducted in pymol). **(G)** Atg8-lipidation is unaffected in *ede1*^AIMR^ mutant cells. Genome replacement of the *EDE1* locus by *ede1*^AIMR^ does not affect Atg8-lipidation as monitored by immunoblotting against Atg8 in wildtype (WT), *ede1*^AIMR^ or Atg7-deficient (*atg7*Δ) cells after addition of 100 nM rapamycin for different durations. Six biologically independent experiments are quantified and the mean ± s.d. is shown. Pgk1 serves as loading control. **(H)** Vacuolar degradation of eGFP-Atg8 is unaffected in *ede1*^AIMR^ mutant cells. N-terminally eGFP-tagged Atg8 under the *ADH* promoter was checked for its autophagic degradation by eGFP-cleavage assay in wildtype (WT), *ede1*^AIMR^ and Atg7-deficient (*atg7*Δ) cells following nitrogen starvation in SD-N medium at different times points (3 and 6 hours) at 30 °C. Pgk1 serves as loading control. **(I)** Bulk autophagy is not affected in *ede1*^AIMR^ mutant cells. Wildtype (WT), *ede1*^AIMR^, Ede1-deficient (*ede1*Δ) and Atg7-deficient (*atg7*Δ) cells were grown to mid-log phase in YPD, and then incubated in SD-N for 3 or 6 hours. Samples were analyzed by the Pho8Δ60 assay (Noda and Klionsky, 2008). The average of four independent experiments is quantified and the mean ± s.d. is shown. **(J)** Mitophagy is not impaired in *ede1*^AIMR^ mutant cells. Mitophagy was monitored by eGFP-tagging of Idh1 in wildtype (WT), *ede1*^AIMR^, and Atg8-deficient (*atg8*Δ) cells, followed by immunoblotting against eGFP before and after induction of mitophagy in YPG medium (YP-glycerol) for the indicated time. Pgk1 serves as loading control. **(K)** Targeting of the Cvt cargo protein Ape1 to the vacuole is not affected in *ede1*^AIMR^ mutant cells. Ape1 processing in the vacuole, i.e. pre-Ape1 (preApe1) to mature Ape1 (mApe1), is monitored before and after addition of 100 nM rapamycin for the indicated time in wildtype (WT), *ede1*^AIMR^, Atg1-deficient (*atg1*Δ) and Atg19-deficient (*atg19*Δ) cells. Cell lysates for each strain were subjected to immunoblotting against Ape1. The average of eight independent experiments is quantified and the mean ± s.d. is shown. Pgk1 serves as loading control.

**Figure S3.**
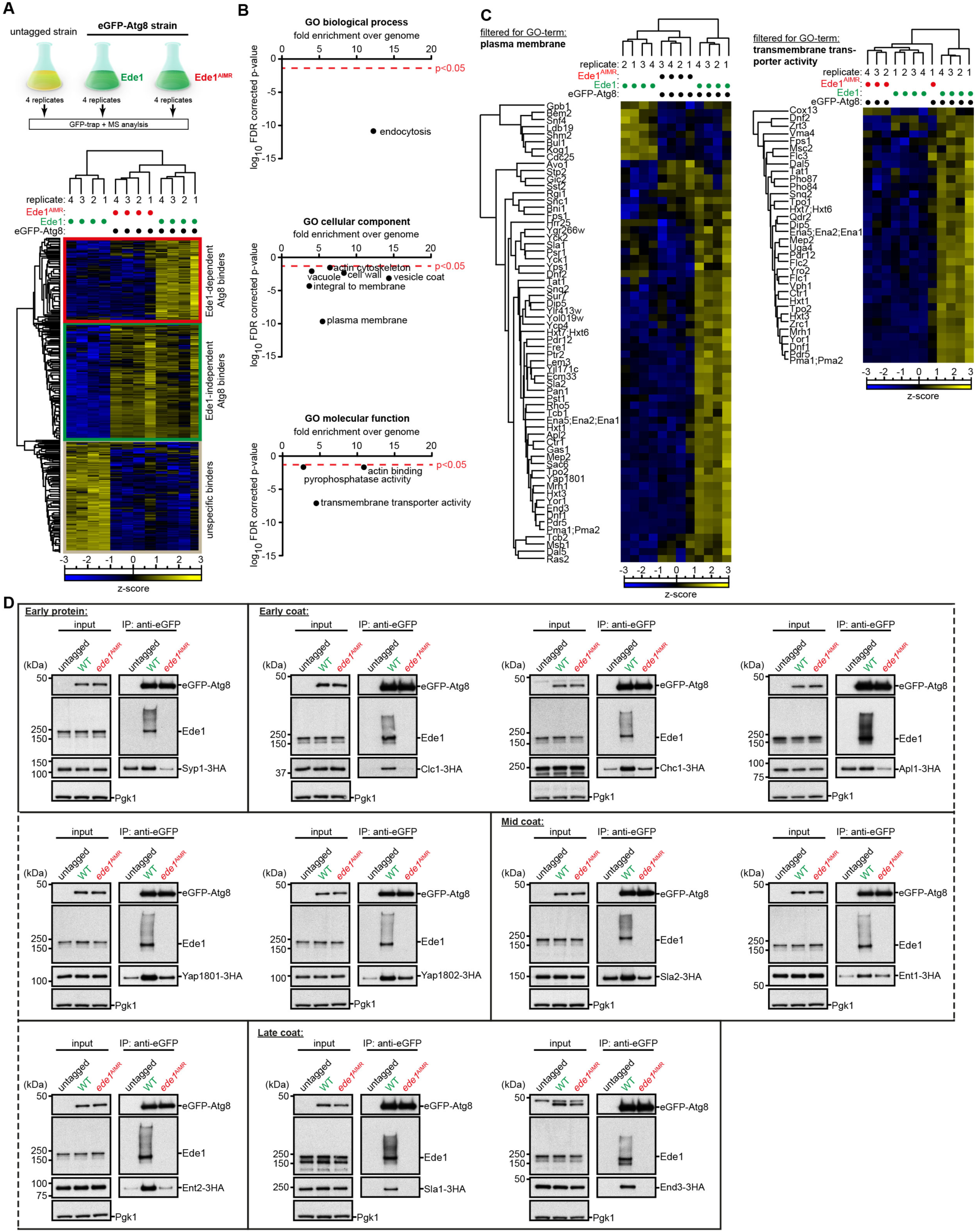
Ede1 Is an Autophagy Receptor for Endocytic Machinery Proteins, Related to Figure 3. **(A)** Workflow of eGFP-Atg8 pulldown with subsequent qMS analysis. Four biological replicates of a strain expressing eGFP-Atg8 under the *ADH* promoter with WT Ede1 or mutant Ede1^AIMR^ present as well as four replicates of an untagged wildtype strain were subjected to immunoprecipitation against eGFP after the induction of autophagy with 100 nM rapamycin for 3 hours and analysed by label-free quantitative MS. Shown is the hierarchical clustering of the respective z-scores for ANOVA-positive and FDR corrected hits from each sample. The red box highlights Ede1-dependent Atg8 interactors. The green box highlights Ede1-independent Atg8 interactors and grey box shows unspecific binders. **(B)** Clathrin-mediated endocytosis and PM proteins bind Atg8 specifically through Ede1. Gene Ontology GO-term analysis of proteins enriched in the eGFP-Atg8 pulldown that bind in an Ede1-dependent manner. The results for the GO enrichment analysis were obtained by using the GENEONTOLOGY search engine (http://geneontology.org). **(C)** Hierarchical cluster analysis of label-free eGFP-Atg8 pulldowns identifies Ede1-dependent Atg8 interactors for GO-term plasma membrane or transmembrane transporter activity. Pulldown and qMS analysis of N-terminally EGFP-tagged Atg8 (p*ADH*) interactors determined from wildtype (WT), *ede1*^AIMR^ or untagged cells. Shown is the hierarchical clustering of the respective z-scores for ANOVA-positive and FDR corrected hits from each sample after filtering proteins significantly enriched for the GO-term (cellular component) plasma membrane (left) or (molecular function) transmembrane transporter activity (right). **(D)** Validation of the Ede1 dependent cargo binding to eGFP-Atg8. Ede1-dependent cargo proteins identified by qMS (Syp1, Clc1, Chc1, Apl1, Yap1801, Yap1802, Sla2, Ent1, Ent2, Sla1 and End3) were C-terminal tagged with 3HA in wildtype (WT) and *ede1*^AIMR^ mutant cells and checked for their binding to eGFP-Atg8 by coimmunoprecipitation using GFP-trap beads. Binding of Ede1 to eGFP-Atg8 was monitored by immunoblotting with an antibody against Ede1. A strain without the eGFP-tagged Atg8 served as control. Pgk1 serves as loading control for the input.

**Figure S4.**
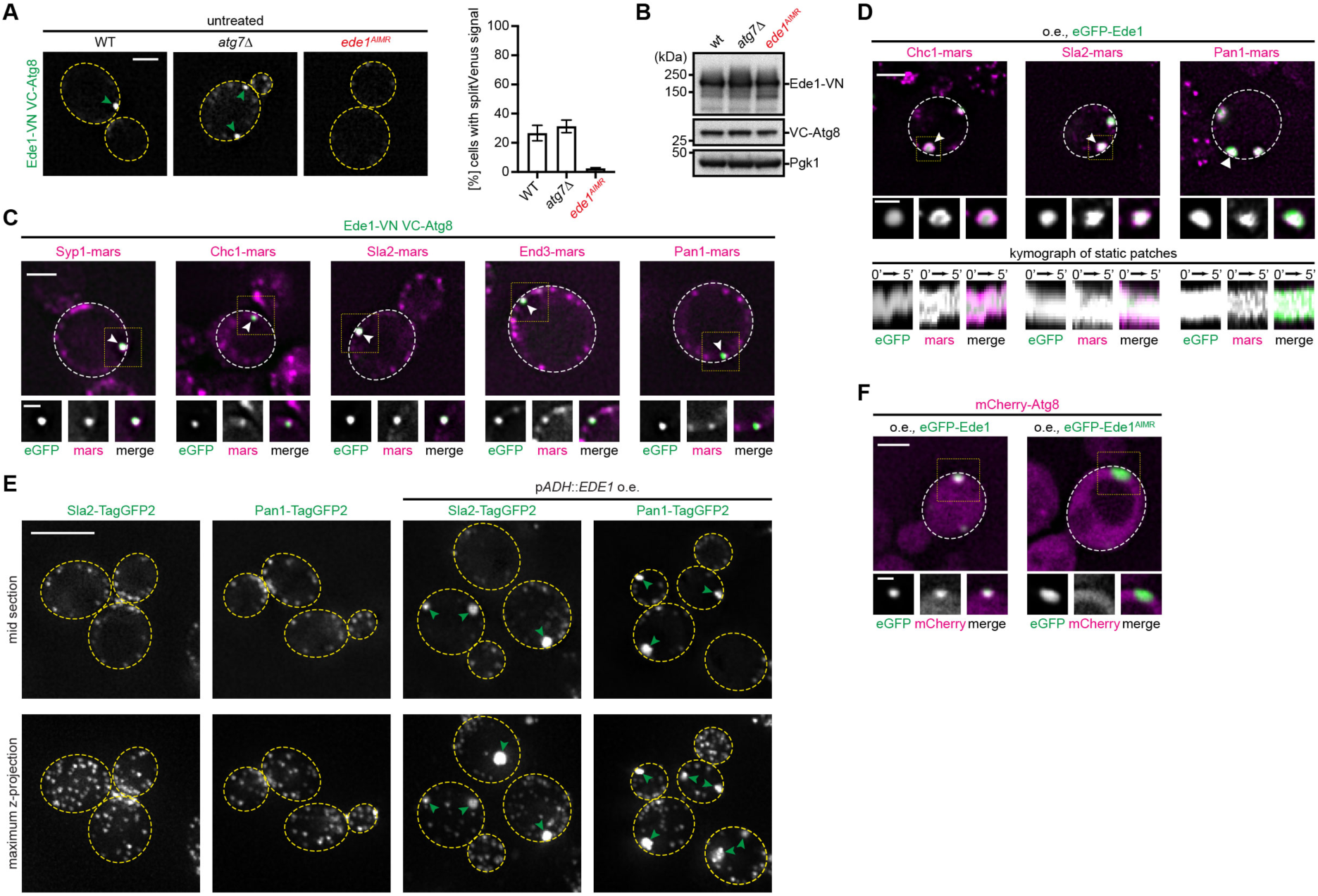
CME Proteins Accumulate in END, Related to Figure 3. **(A)** Ede1 interacts with Atg8 at distinct sites at the cell periphery. Interaction of Ede1 with Atg8 is shown by BiFC fluorescent signal resulting from VC-Atg8 (p*ADH*) and Ede1-VN in WT, *atg7*Δ, or *ede1*^AIMR^ cells. Arrowheads indicate interaction sites at the PM. Quantifications are based on 500 or more cells from two independent experiments and displayed as mean ± s.d.. Scale bar represents 2 µm. **(B)** Protein levels of splitVenus (VN and VC) tagged Ede1 and Atg8 are unchanged in different mutant strains. Immunoblotting against Ede1 and Atg8 with the respective antibodies in WT, Atg7-deficient (*atg7*Δ), or *ede1*^AIMR^ mutant cells. **(C)** Endocytic proteins accumulate at sites of Ede1/Atg8 interaction. Different mars-tagged proteins from the endocytic machinery (Syp1 (early protein), Chc1 (early coat), Sla2 (mid coat) and Pan1/End3 (late coat)) colocalize to sites of Ede1-Atg8 interaction marked by the BiFC signal of p*ADH::*VC-*ATG8* and Ede1-VN. Scale bar represents 2 µm and 1 µm for inset. **(D)** The CME proteins Chc1, Sla2 and Pan1 stably colocalize at sites of eGFP-Ede1 accumulation. Kymograph representation of a two-color movie (1 frame/20 s) recorded by fluorescence microscopy of cells overexpressing eGFP-Ede1 under the control of the *ADH* promoter and either Chc1-mars, Sla2-mars, or Pan1-mars. Scale bar represents 2 µm or 1 µm for inlets and kymographs. **(E)** Stable accumulation of CME proteins is independent of the eGFP-tag. Sla2 and Pan1 where endogenously tagged with the monomeric fluorescent protein TagGFP2 in cells where Ede1 expression was either controlled by the endogenous or the *ADH* promoter. Green arrows indicate sites of CME protein accumulation. Scale bar represents 5 µm. **(F)** Atg8 co-localizes AIM-dependent with eGFP-Ede1 clusters in Ede1 overexpressing cells. Colocalization experiments between mCherry-Atg8 (p*ADH*) and eGFP-Ede1 or eGFP-Ede1^AIMR^ (p*ADH*). Lower panels show magnifications of boxed areas. Scale bar represents 2 µm and 1 µm for inset.

**Figure S5.**
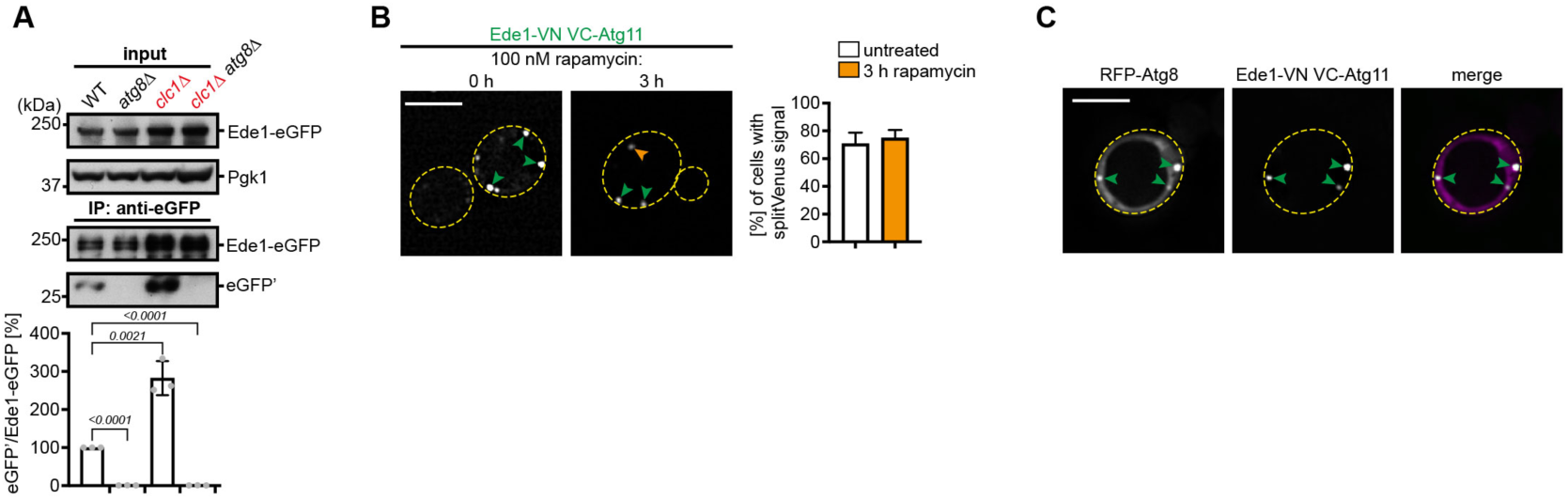
Involvement of Atg11 Scaffold in Autophagic END Degradation, Related to Figure 4. **(A)** Autophagic turnover of Ede1 is detected during logarithmic growth in WT, and enhanced in *clc1*Δ mutant cells. eGFP pulldowns were performed in WT or *clc1*Δ mutant cells expressing endogenously eGFP-tagged Ede1. The pulldown of vacuolar enriched free eGFP from WT or *clc1*Δ mutant was compared to cells additionally deficient in the core autophagy protein Atg8 (*atg8*Δ). A quantification of the ratio between free eGFP’ and full length Ede1-eGFP is shown. Data are mean ± s.d. of n = 3 biologically independent experiments. Statistical analysis was performed using two-tailed Student’s *t*-tests, *p*-values are indicated. Pgk1 serves as input control. **(B)** Ede1 interacts with Atg11 at similar sites as seen for Atg8. Interaction of Ede1 with Atg11 is shown by BiFC fluorescent signal resulting from VC-Atg11 (p*ADH*) and Ede1-VN before and after a 3 hour treatment with 100 nM rapamycin. Green arrowheads indicate interaction sites at the END, whereas orange arrowheads indicate interaction sites at the PAS. Quantifications are based on 500 or more cells from two independent experiments and displayed as mean ± s.d.. Scale bar represents 5 µm. **(C)** RFP-Atg8 co-localizes to sites of Ede1/Atg11 interaction. Cells expressing p*ADH::*VC-*ATG11* and Ede1-VN accumulate RFP-Atg8 at sites of BiFC signal. Arrowheads indicate interaction sites at the END. Scale bar represents 5 µm.

**Figure S6.**
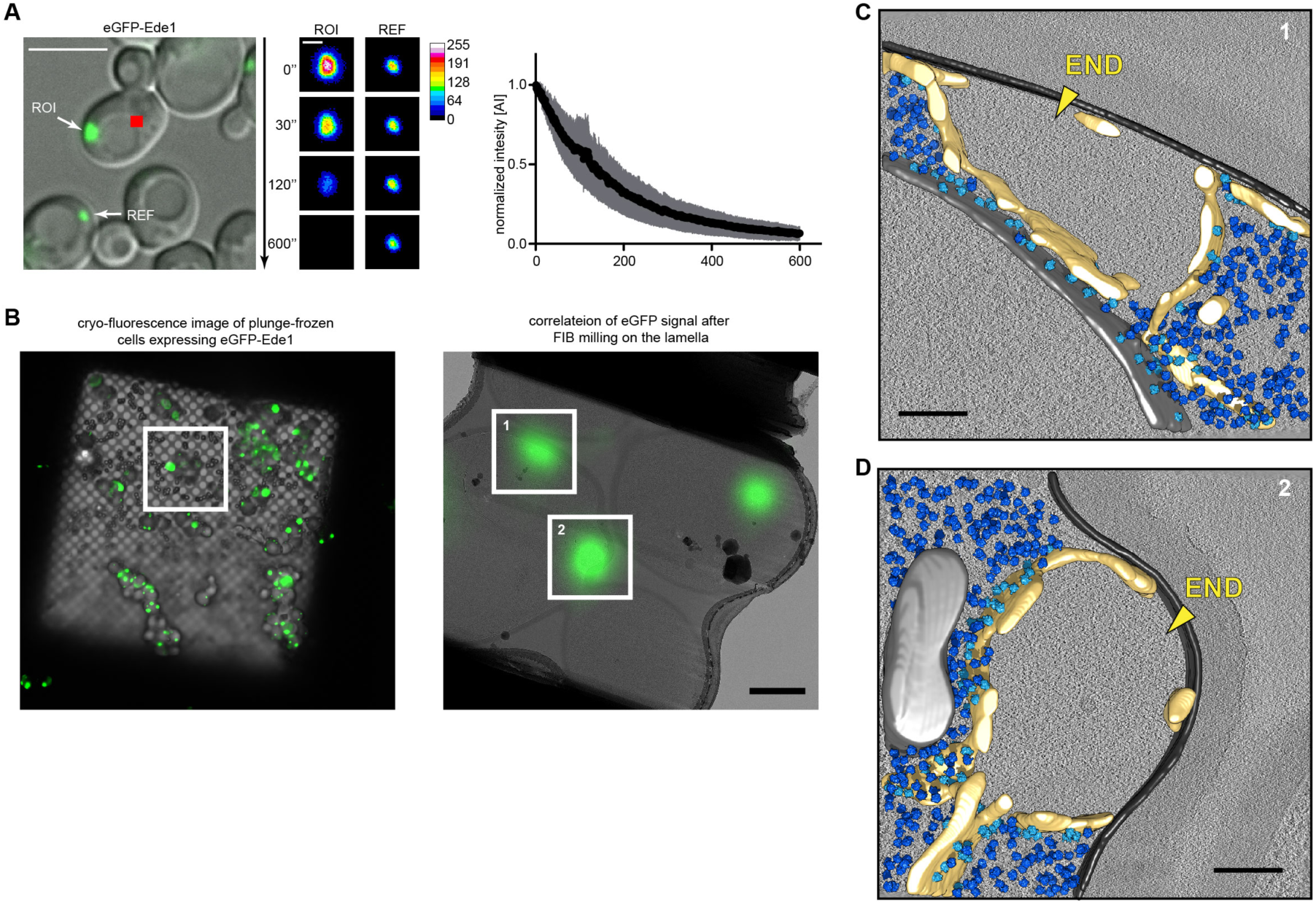
Biophysical and Ultrastructural Dissection of END, Related to Figure 5. **(A)** eGFP-Ede1 adjacent to the PM can readily exchange with the cytosol. Fluorescence loss in photobleaching (FLIP) experiments were performed for cells overexpressing eGFP-Ede1 under the control of the *ADH* promoter. The indicated spot (red) is bleached after each image for a time course of 600 s and the fluorescence within the region of interest (ROI) is measured. As reference for bleaching effects serves the END of a neighboring cell. Representative images of one experiment are shown. Quantification shows the loss of eGFP-Ede1 signal over time from at least three independent experiments as mean ± s.d.. Scale bar represents 5 µm and 1 µm for inset. n = 4 biologically independent experiments. **(B-D)** Ultrastructural dissection of the Ede1 accumulations in the overexpression strain by 3D-correlated cryo-light and cryo-electron microscopy (cryo-CLEM). After cryopreservation, potential sites of interest are located by fluorescence light microscopy (FLM) ((B), left image) and scanning electron microscopy (SEM). Correlative beads are both located in FLM and the ion beam (IB) mode of a dual beam focused ion beam (FIB) SEM to obtain a 3D correlation between both imaging modes. Predicted points of interests are then calculated from the FLM 3D stacks data, promising sites are selected and cut using the cryo-FIB resulting in thin lamellas. Finally, 2D correlation can be performed between beads in the FLM and transmission electron microscope (cryo-TEM) images to obtain points of interest on the lamellas ((B), right image), which are then analyzed in detail by high-resolution cryo-TEM (cryo-CLEM). In this example two tomograms are recorded at the sites of correlated signal ((B) right image 1 and 2, corresponding to C and D), reconstructed and segmented. Both examples (C and D) show areas of Ede1 protein accumulations, which are surrounded by endoplasmic reticulum. The interior of the amorphous protein structure is clearly devoid of all ribosomes (light and dark blue) and originates directly from the plasma membrane (light grey). Membrane-associated ribosomes (light blue) are exclusively found on parts of the ER facing the cytosol. Other organelles in the vicinity, such as the nuclear membrane (light grey, C) or mitochondria (light grey, D), appear unchanged. The scale bar in (B, left) represents 20 µm, in (B, right) 1 µm and (C and D) 200 nm.

**Figure S7.**
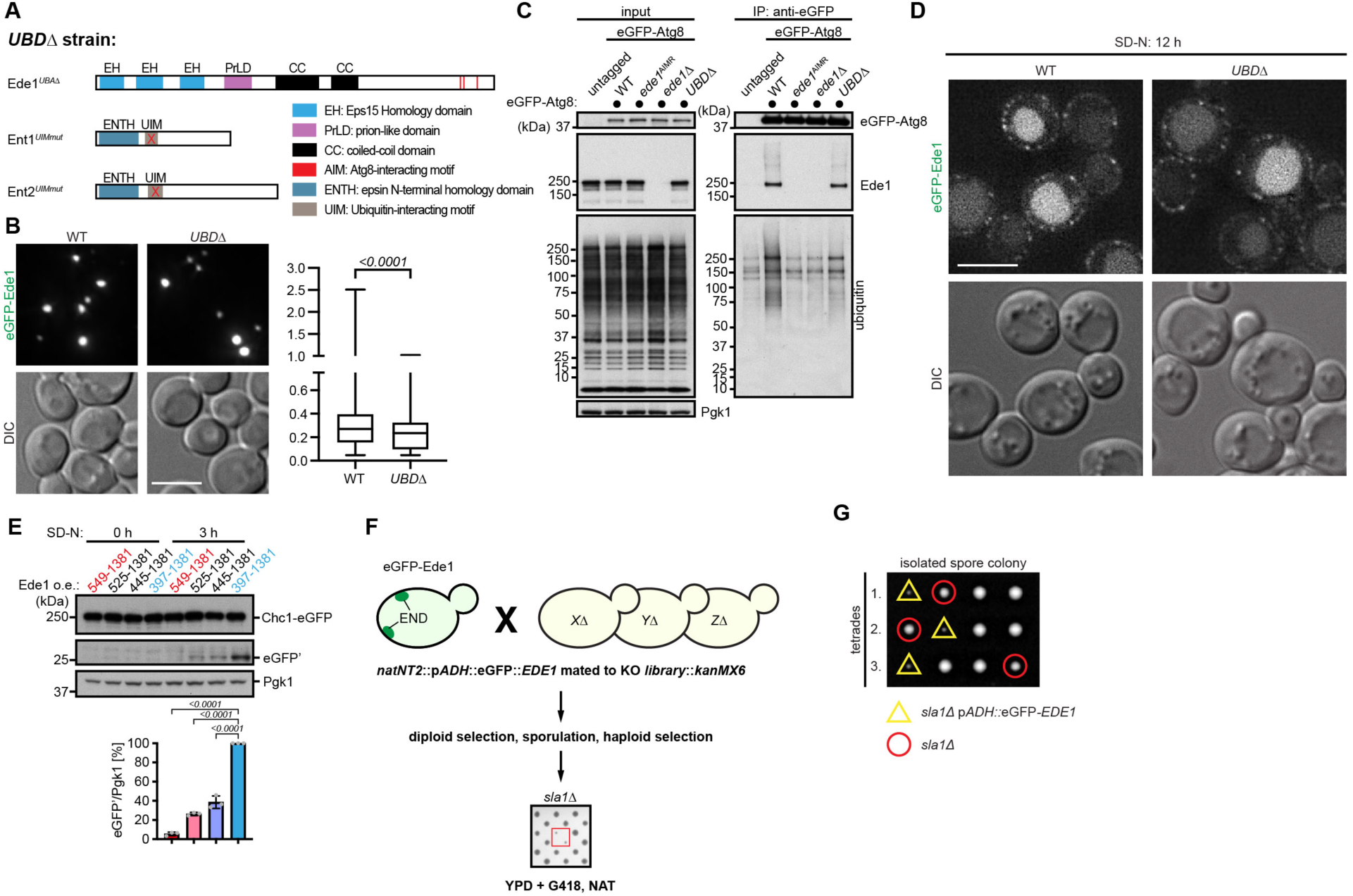
Phase-Separation of Ede1 but Not Ubiquitin Binding Is Important for Autophagic Degradation, Related to Figure 6. **(A-B)** Ubiquitin binding motifs within endocytic machinery proteins are dispensable for END formation. Illustration shows mutants used to create the *UBD*Δ strain (A). EH: Eps15-homology domain; PrLD: prion-like domain; CC: coiled-coil domain; UBA: ubiquitin-associated domain; ENTH: epsin N-terminal homology domain; UIM: Ubiquitin-interacting motif. END formation driven by expression of Ede1 under the control of the *ADH* promoter is monitored by fluorescence microscopy of WT and *UBD*Δ (deletion of the UBA domain of Ede1, and mutation of both UIMs in Ent1 and Ent2) mutant cells (B). Spots size was analyzed computational from maximum z-projections of the corresponding strains for a total of more than 3000 spots of n = 6 biologically independent experiments. **(C)** Ede1-Atg8 interaction is unaffected in *UBD*Δ cells but binding of ubiquitylated cargo proteins is decreased. Atg8 expressed under the *ADH* promoter was eGFP-tagged, and its interaction with Ede1 was probed by co-immunoprecipitation and immunoblotting with the respective antibodies in WT, *ede1*^AIMR^, *ede1*Δ, or *UBD*Δ cells. Pgk1 was used as loading control for the input. **(D)** Autophagic turnover of Ede1 in *UBD*Δ cells is unaffected. Cells expressing eGFP-Ede1 (p*ADH*) in WT or *UBD*Δ cells were starved for 12 hours in medium lacking nitrogen before monitored by fluorescence microscopy. **(E)** PrLD of Ede1 is necessary for autophagic degradation of Ede1-dependent substrate Chc1. Cells in which Chc1 was endogenously eGFP-tagged and co-expressing different N-terminal Ede1 truncation mutants from the *ADH* promoter were analyzed by eGFP-cleavage assay before and after 3 hours of nitrogen starvation. Quantifications of free eGFP’ levels normalized to Pgk1 are shown. Data are mean ± s.d. of n = 3 biologically independent experiments. Statistical analysis was performed using two-tailed Student’s *t*-tests, *p*-values are indicated. **(F)** Scheme of the robot-based synthetic lethal screen, which identified the strong synthetic interaction between overexpression of Ede1 and Sla1. A library of nonessential gene deletions (all G418resistant) was mated with a bait strain overexpressing Ede1 from the *ADH* promoter (NAT resistant). Diploid cells were selected and subjected to sporulation. Finally, haploid cells were selected by pinning on YPD plates containing both selection markers G418 and NAT. **(G)** Manual crossing of a *sla1*Δ with the eGFP-Ede1 overexpression strain followed by tetrad dissection confirms the screening results.

Supplementary Movie 1 related to Figure 5:

Movie for single cell analysis experiment presented in Figure 5D. Shown here on the left is the imaging of Ede1-eGFP in wildtype cells before, during and after treatment with 10% 1-,6- Hexanediol and 10 µg/ml digitonin. Shown on the right is the DIC channel.

Supplementary Movie 2 related to Figure 5:

Movie for single cell analysis experiment presented in Figure 5D. Shown here on the left is the imaging of Ede1-eGFP in *yap1801*Δ, *yap1802*Δ, and *apl3*Δ (ΔΔΔ) mutant cells before, during and after treatment with 10% 1-,6-Hexanediol and 10 µg/ml digitonin. Shown on the right is the DIC channel.

Supplementary Movie 3 related to Figure 5:

Original cryo-ET data with an animated segmentation presented in Figure 5E-F.

Supplementary Movie 4 related to Supplement Figure S6:

Original cryo-ET data presented in Supplementary Figure S6C.

Supplementary Movie 5 related to Supplement Figure S6:

Original cryo-ET data presented in Supplementary Figure S6D

Supplementary Movie 6 related to Figure 7:

Movie for single cell analysis experiment presented in Figure 7A. Shown here on the left is the dual imaging of eGFP-Ede1 (p*ADH*) (green) and mCherry-Atg8 (p*ADH*) (magenta) and on the right the DIC channel in the wildtype background over time.

Supplementary Movie 7 related to Figure 7:

Movie for single cell analysis experiment presented in Figure 7A. Shown here on the left is the dual imaging of eGFP-Ede1 (p*ADH*) (green) and mCherry-Atg8 (p*ADH*) (magenta) and on the right the DIC channel in the Atg7-deficient (*atg7*Δ) background over time.

Supplementary Movie 8 related to Figure 7:

Original cryo-ET data with an animated segmentation presented in Figure 7C.

**Supplementary Table 1.** Crystallography data collection and refinement

|  | ScEde1 (1220-1247) – ScAtg8 (1-116) |
| --- | --- |
| **Data collection** |  |
| Space group | P4<sub>3</sub> |
| Cell dimensions |  |
| *a*, *b*, *c* (Å) | 49.642, 49.642, 123.446 |
| $\alpha$, $\beta$, $\gamma$ (°) | 90, 90, 90 |
| Wavelength (Å) | 0.979 |
| Resolution (Å)* | 46.06-1.773 (1.836-1.773) |
| R<sub>sym</sub> or R<sub>merge</sub> * | 0.054 (0.060) |
| *I* / $\sigma$ (*I*) * | 29.36 (4.62) |
| *CC* $\frac{1}{2}$ * | 0.996 (0.862) |
| Completeness (%) * | 99.97 (99.65) |
| Wilson B-factor (Å<sup>2</sup>) | 24.92 |
| Redundancy * | 8.8 (7.6) |
| **Refinement** |  |
| Resolution (Å) | 46.06-1.773 |
| No. reflections | 28859 |
| R<sub>work</sub> * | 0.1756 (0.2240) |
| R<sub>free</sub> * | 0.2028 (0.2676) |
| Completeness (%) | 99.94 |
| No. atoms | 2421 |
| Protein | 2205 |
| Water | 216 |
| B factors (Å<sup>2</sup>) | 28.90 |
| Protein | 28.20 |
| Solvent | 36.60 |
| R.m.s. deviations |  |
| Bond lengths (Å) | 0.008 |
| Bond angles (°) | 1.17 |
| Ramachandran plot |  |
| Favored (%) | 98 |
| Outliers (%) | 0 |
| Clashscore | 2.72 |

\* Value in parenthesis are for the highest resolution shell.

**Supplementary Table 2.**
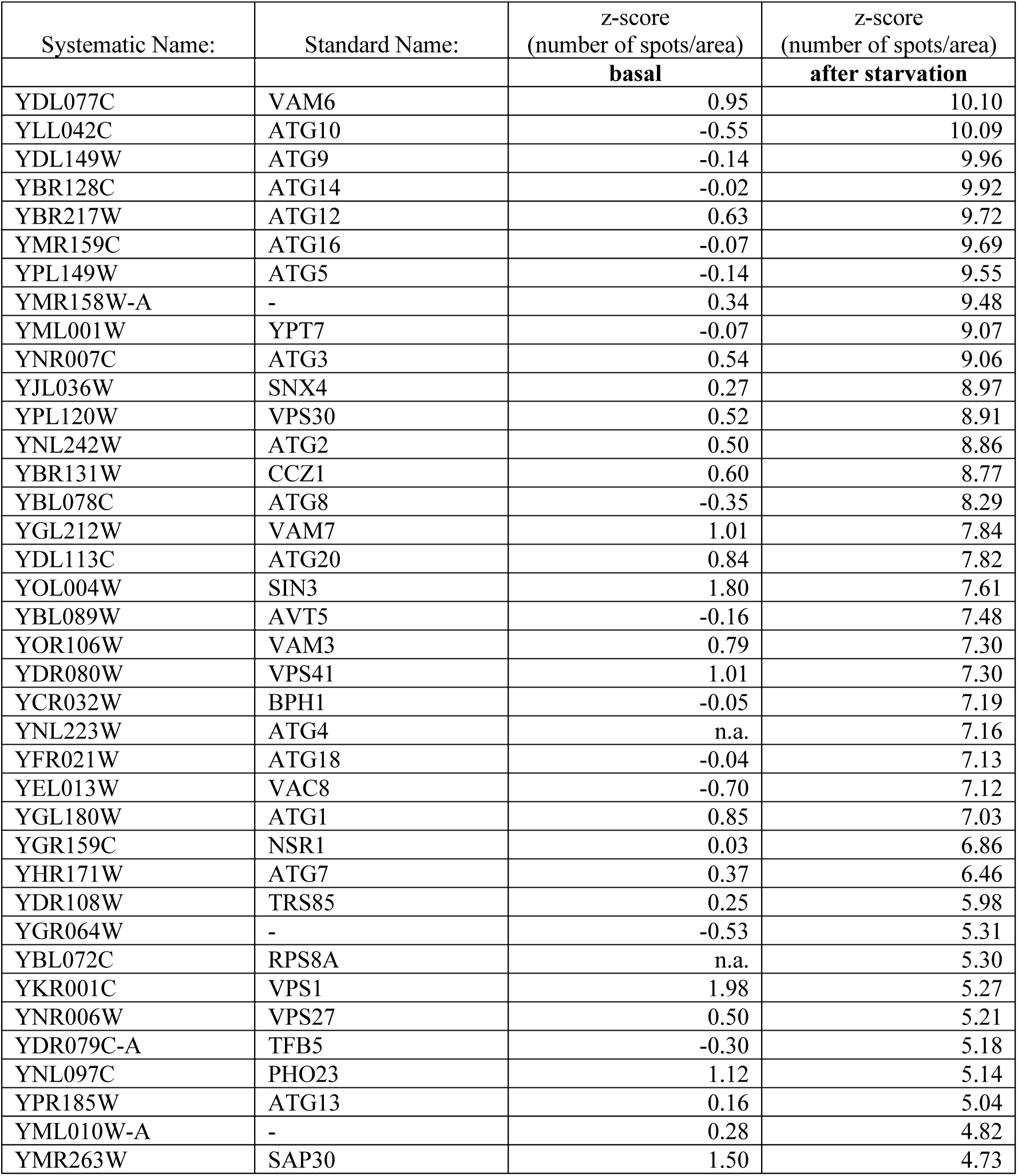
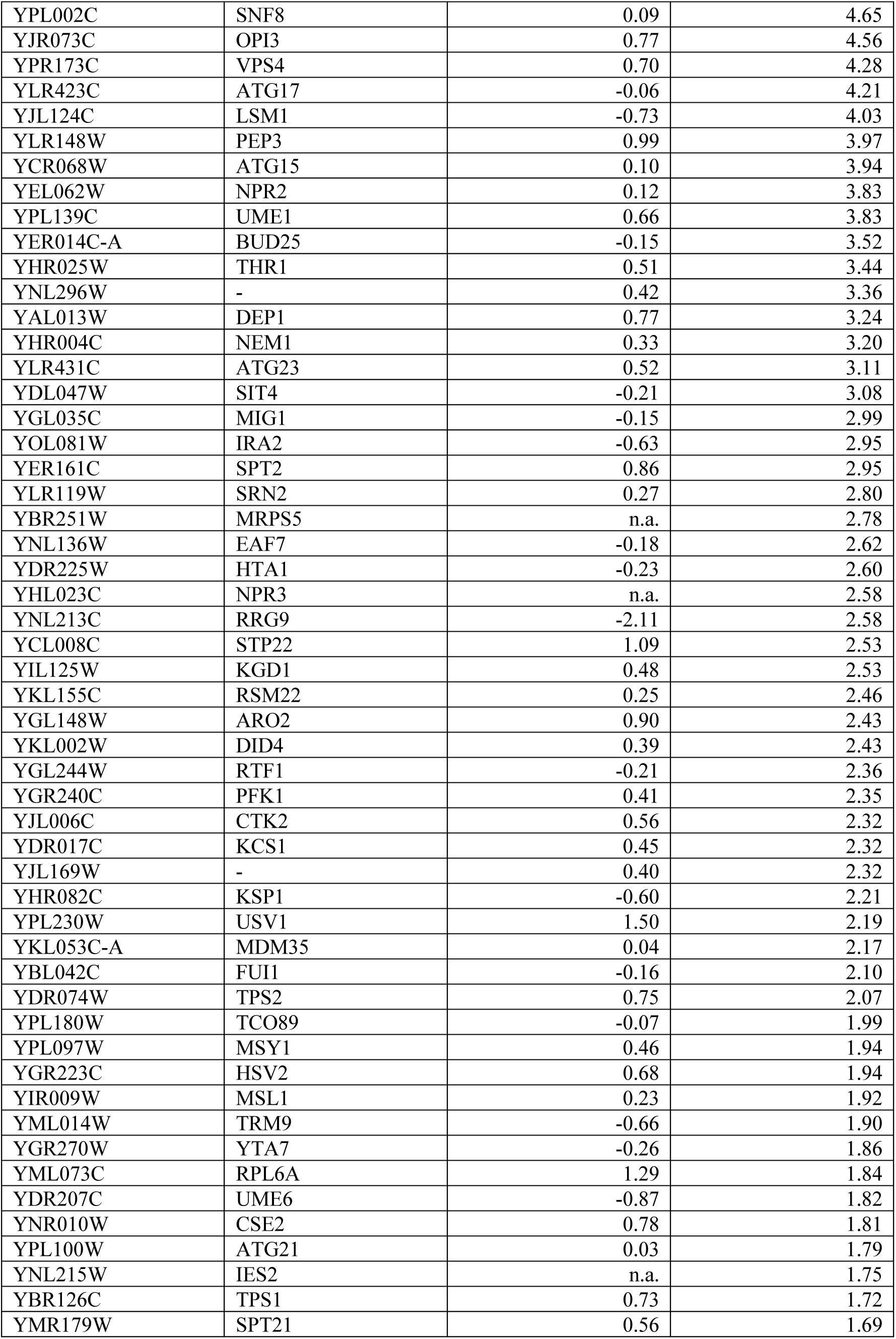
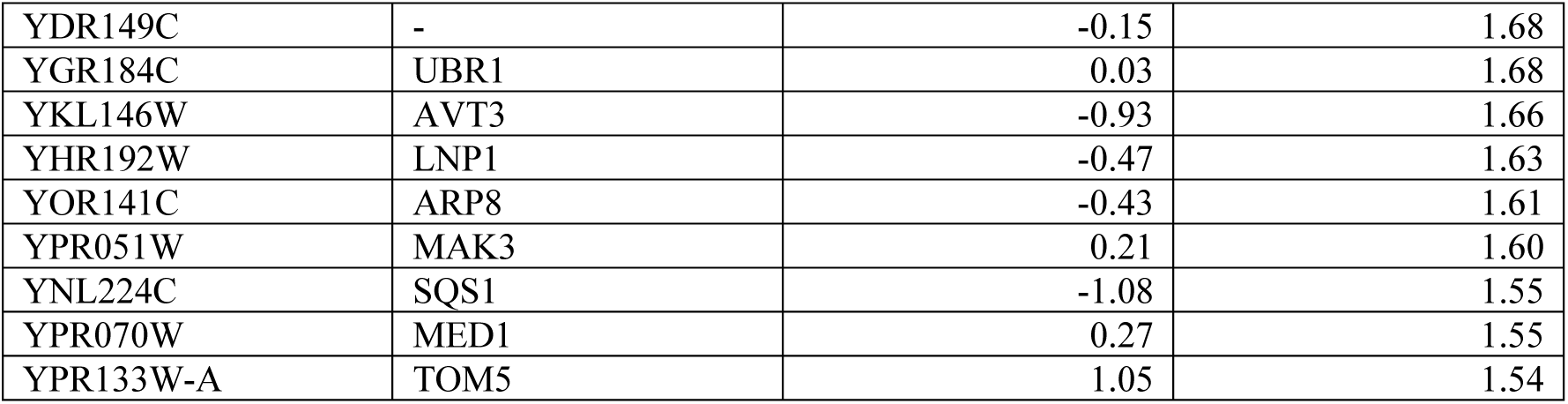
Gene knockouts that stabilize the END compartment upon nitrogen starvation.

| Systematic Name: | Standard Name: | z-score<br>(number of spots/area) | z-score<br>(number of spots/area) |
| --- | --- | --- | --- |
|  |  | **basal** | **after starvation** |
| YDL077C | VAM6 | 0.95 | 10.10 |
| YLL042C | ATG10 | -0.55 | 10.09 |
| YDL149W | ATG9 | -0.14 | 9.96 |
| YBR128C | ATG14 | -0.02 | 9.92 |
| YBR217W | ATG12 | 0.63 | 9.72 |
| YMR159C | ATG16 | -0.07 | 9.69 |
| YPL149W | ATG5 | -0.14 | 9.55 |
| YMR158W-A | - | 0.34 | 9.48 |
| YML001W | YPT7 | -0.07 | 9.07 |
| YNR007C | ATG3 | 0.54 | 9.06 |
| YJL036W | SNX4 | 0.27 | 8.97 |
| YPL120W | VPS30 | 0.52 | 8.91 |
| YNL242W | ATG2 | 0.50 | 8.86 |
| YBR131W | CCZ1 | 0.60 | 8.77 |
| YBL078C | ATG8 | -0.35 | 8.29 |
| YGL212W | VAM7 | 1.01 | 7.84 |
| YDL113C | ATG20 | 0.84 | 7.82 |
| YOL004W | SIN3 | 1.80 | 7.61 |
| YBL089W | AVT5 | -0.16 | 7.48 |
| YOR106W | VAM3 | 0.79 | 7.30 |
| YDR080W | VPS41 | 1.01 | 7.30 |
| YCR032W | BPH1 | -0.05 | 7.19 |
| YNL223W | ATG4 | n.a. | 7.16 |
| YFR021W | ATG18 | -0.04 | 7.13 |
| YEL013W | VAC8 | -0.70 | 7.12 |
| YGL180W | ATG1 | 0.85 | 7.03 |
| YGR159C | NSR1 | 0.03 | 6.86 |
| YHR171W | ATG7 | 0.37 | 6.46 |
| YDR108W | TRS85 | 0.25 | 5.98 |
| YGR064W | - | -0.53 | 5.31 |
| YBL072C | RPS8A | n.a. | 5.30 |
| YKR001C | VPS1 | 1.98 | 5.27 |
| YNR006W | VPS27 | 0.50 | 5.21 |
| YDR079C-A | TFB5 | -0.30 | 5.18 |
| YNL097C | PHO23 | 1.12 | 5.14 |
| YPR185W | ATG13 | 0.16 | 5.04 |
| YML010W-A | - | 0.28 | 4.82 |
| YMR263W | SAP30 | 1.50 | 4.73 |
| YPL002C | SNF8 | 0.09 | 4.65 |
| YJR073C | OPI3 | 0.77 | 4.56 |
| YPR173C | VPS4 | 0.70 | 4.28 |
| YLR423C | ATG17 | -0.06 | 4.21 |
| YJL124C | LSM1 | -0.73 | 4.03 |
| YLR148W | PEP3 | 0.99 | 3.97 |
| YCR068W | ATG15 | 0.10 | 3.94 |
| YEL062W | NPR2 | 0.12 | 3.83 |
| YPL139C | UME1 | 0.66 | 3.83 |
| YER014C-A | BUD25 | -0.15 | 3.52 |
| YHR025W | THR1 | 0.51 | 3.44 |
| YNL296W | - | 0.42 | 3.36 |
| YAL013W | DEP1 | 0.77 | 3.24 |
| YHR004C | NEM1 | 0.33 | 3.20 |
| YLR431C | ATG23 | 0.52 | 3.11 |
| YDL047W | SIT4 | -0.21 | 3.08 |
| YGL035C | MIG1 | -0.15 | 2.99 |
| YOL081W | IRA2 | -0.63 | 2.95 |
| YER161C | SPT2 | 0.86 | 2.95 |
| YLR119W | SRN2 | 0.27 | 2.80 |
| YBR251W | MRPS5 | n.a. | 2.78 |
| YNL136W | EAF7 | -0.18 | 2.62 |
| YDR225W | HTA1 | -0.23 | 2.60 |
| YHL023C | NPR3 | n.a. | 2.58 |
| YNL213C | RRG9 | -2.11 | 2.58 |
| YCL008C | STP22 | 1.09 | 2.53 |
| YIL125W | KGD1 | 0.48 | 2.53 |
| YKL155C | RSM22 | 0.25 | 2.46 |
| YGL148W | ARO2 | 0.90 | 2.43 |
| YKL002W | DID4 | 0.39 | 2.43 |
| YGL244W | RTF1 | -0.21 | 2.36 |
| YGR240C | PFK1 | 0.41 | 2.35 |
| YJL006C | CTK2 | 0.56 | 2.32 |
| YDR017C | KCS1 | 0.45 | 2.32 |
| YJL169W | - | 0.40 | 2.32 |
| YHR082C | KSP1 | -0.60 | 2.21 |
| YPL230W | USV1 | 1.50 | 2.19 |
| YKL053C-A | MDM35 | 0.04 | 2.17 |
| YBL042C | FUI1 | -0.16 | 2.10 |
| YDR074W | TPS2 | 0.75 | 2.07 |
| YPL180W | TCO89 | -0.07 | 1.99 |
| YPL097W | MSY1 | 0.46 | 1.94 |
| YGR223C | HSV2 | 0.68 | 1.94 |
| YIR009W | MSL1 | 0.23 | 1.92 |
| YML014W | TRM9 | -0.66 | 1.90 |
| YGR270W | YTA7 | -0.26 | 1.86 |
| YML073C | RPL6A | 1.29 | 1.84 |
| YDR207C | UME6 | -0.87 | 1.82 |
| YNR010W | CSE2 | 0.78 | 1.81 |
| YPL100W | ATG21 | 0.03 | 1.79 |
| YNL215W | IES2 | n.a. | 1.75 |
| YBR126C | TPS1 | 0.73 | 1.72 |
| YMR179W | SPT21 | 0.56 | 1.69 |
| YDR149C | - | -0.15 | 1.68 |
| YGR184C | UBR1 | 0.03 | 1.68 |
| YKL146W | AVT3 | -0.93 | 1.66 |
| YHR192W | LNP1 | -0.47 | 1.63 |
| YOR141C | ARP8 | -0.43 | 1.61 |
| YPR051W | MAK3 | 0.21 | 1.60 |
| YNL224C | SQS1 | -1.08 | 1.55 |
| YPR070W | MED1 | 0.27 | 1.55 |
| YPR133W-A | TOM5 | 1.05 | 1.54 |

**Supplementary Table 3.**
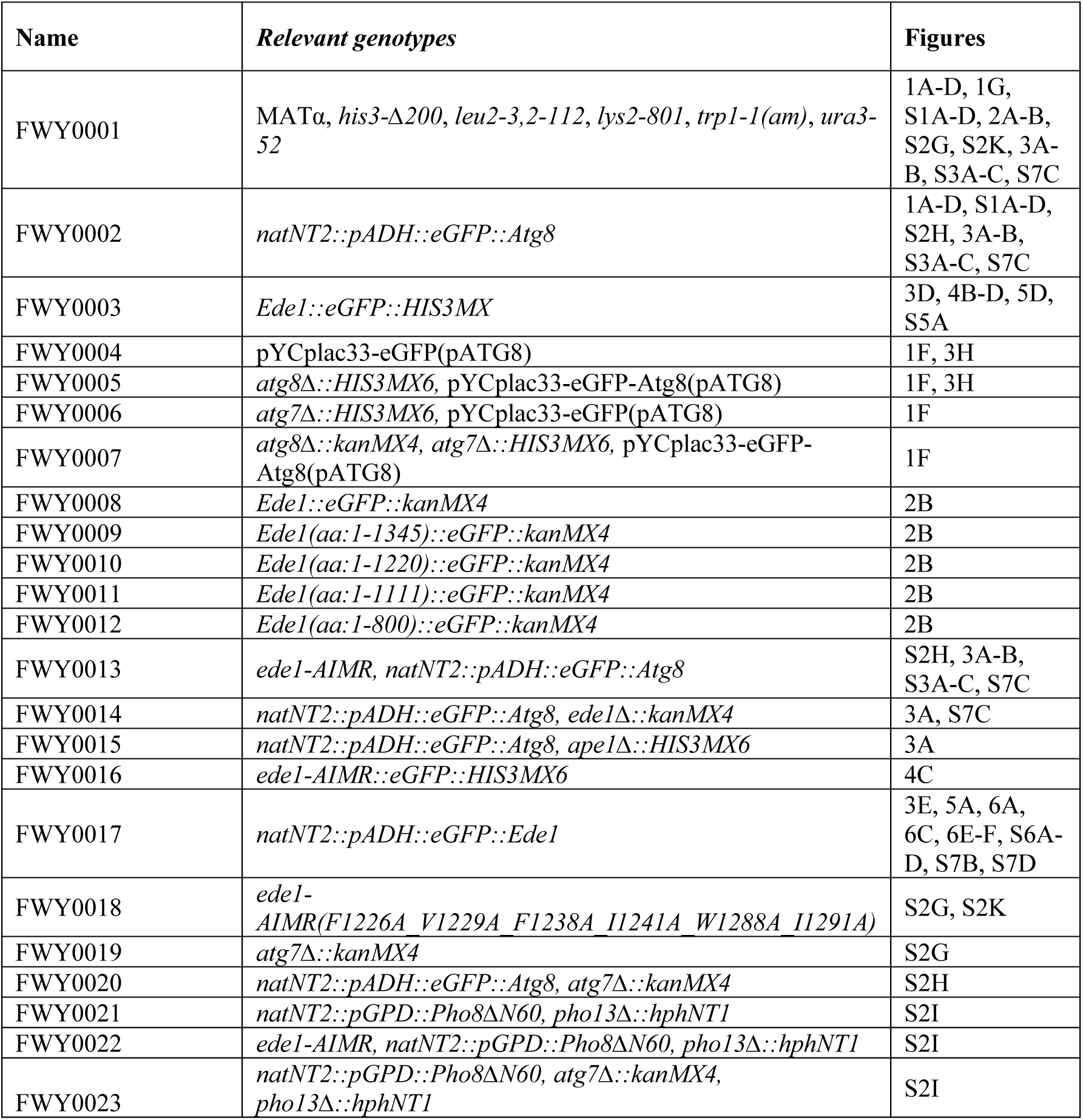
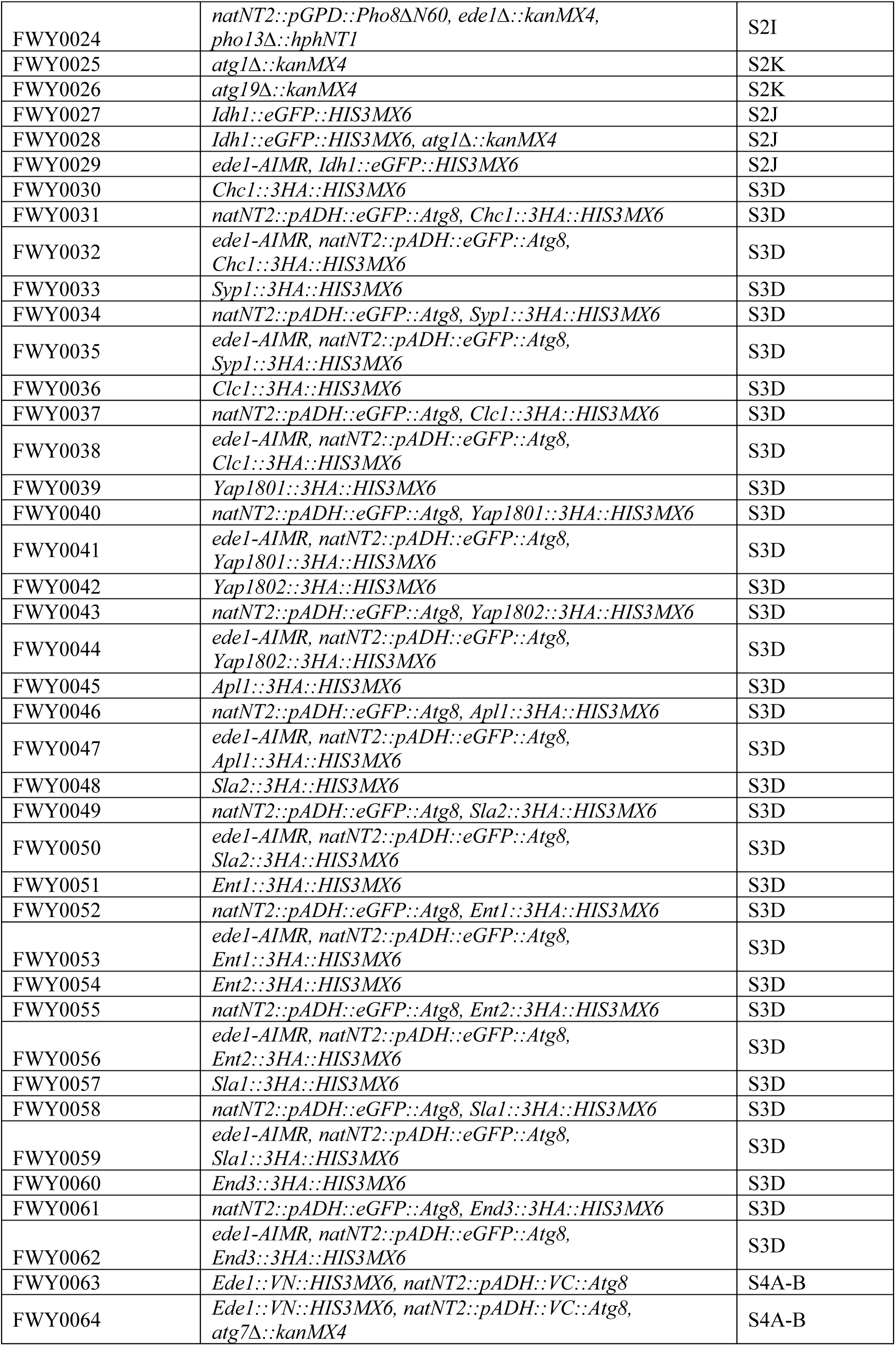
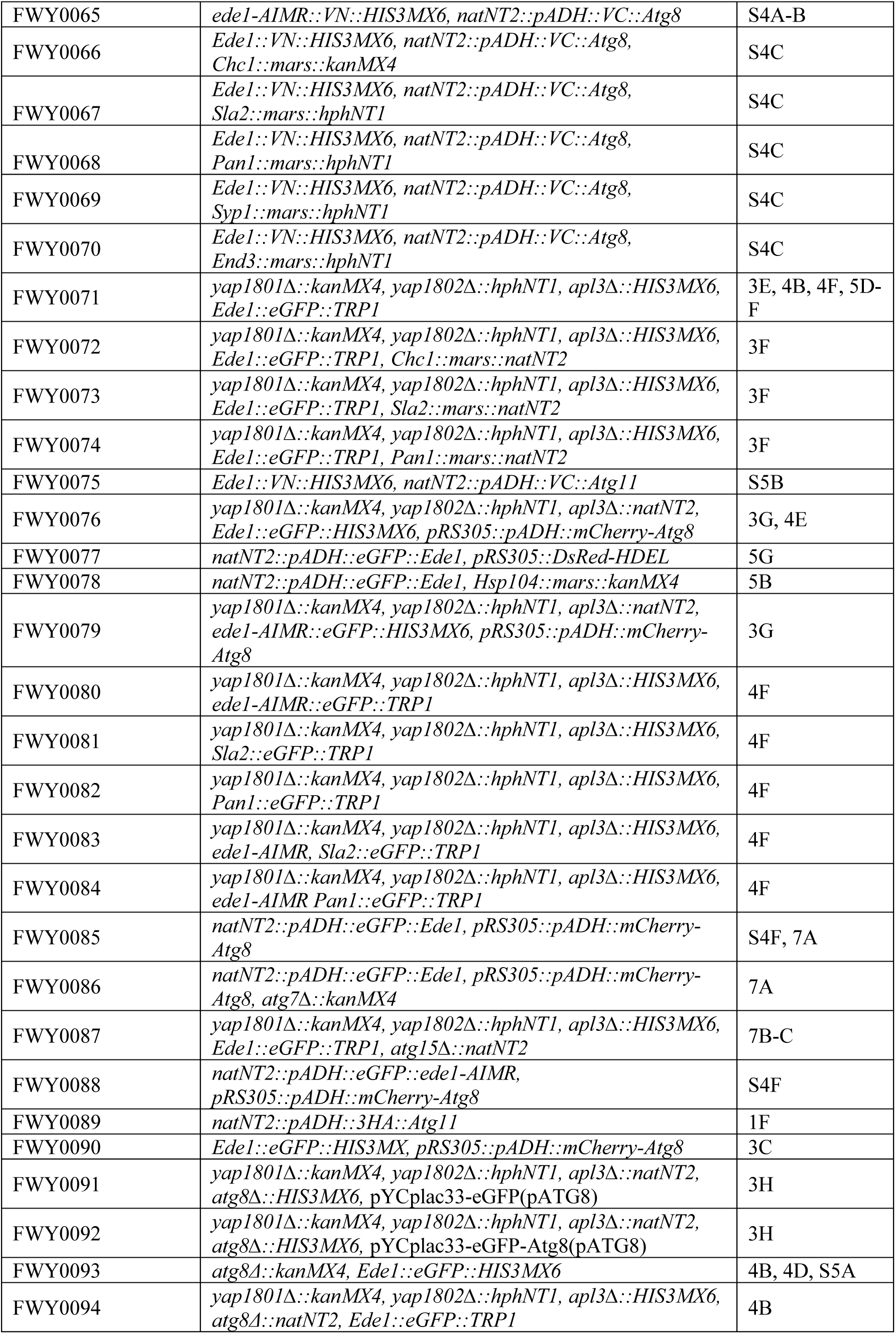
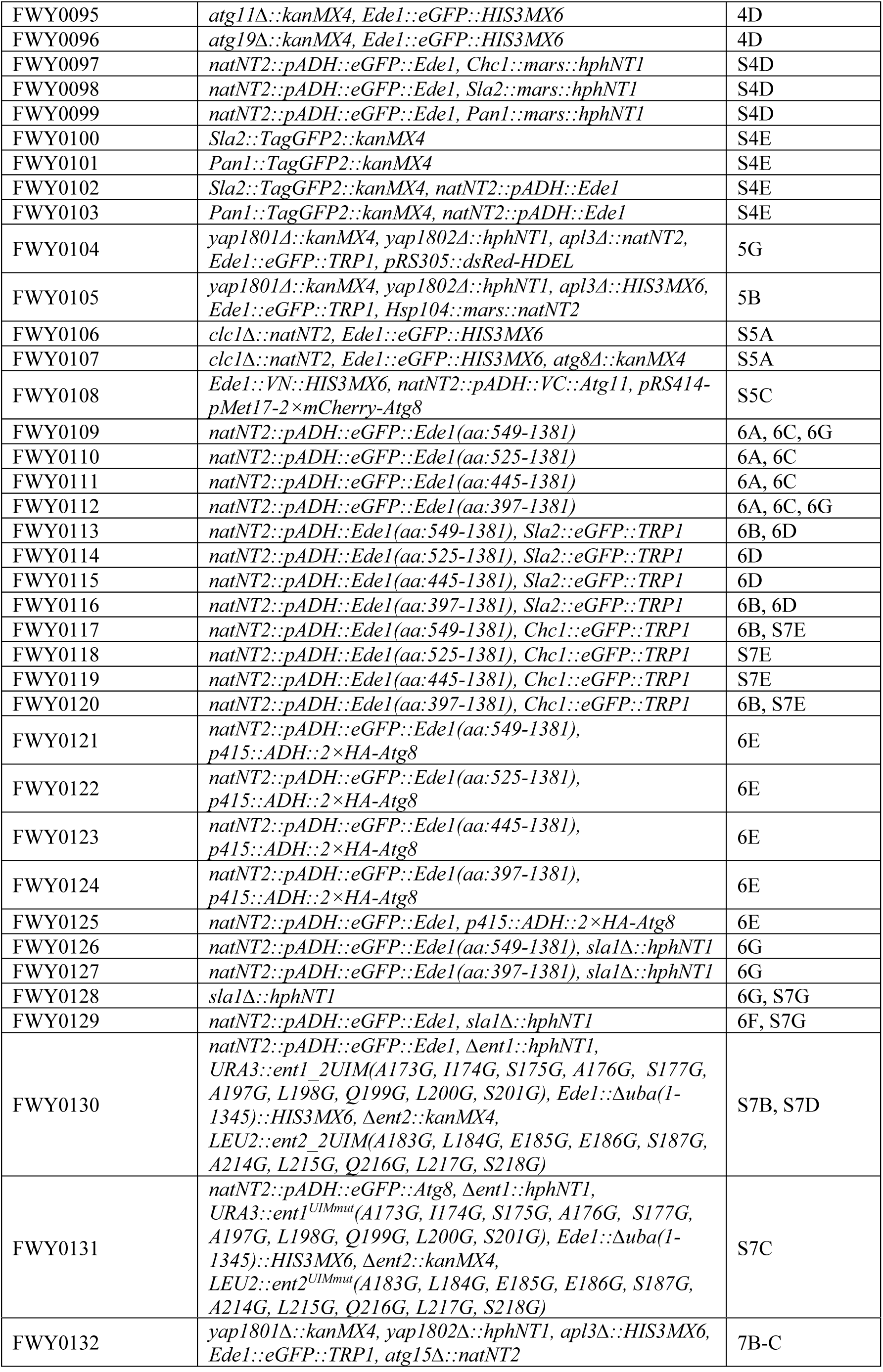
Yeast strains used in this study.

**Supplementary Table 4.** Plasmids used in this study.

| Name | Expression construct | Figures |
| --- | --- | --- |
| GST | *pGEX-4T3* | 1E, 2A-C, S2C |
| GST-Ede1 | *pGEX-4T3-Ede1* | 1E, 2C, S2C |
| GST-Syp1 | *pGEX-4T3-Syp1* | 1E |
| GST-Clc1 | *pGEX-4T3-Clc1* | 1E |
| GST-Ent2 | *pGEX-4T3-Ent2* | 1E |
| GST-End3 | *pGEX-4T3-End3* | 1E |
| GST-Pan1 | *pGEX-4T3-Pan1* | 1E |
| His-Atg8 | *pET28a-Atg8* | 1E, 2C, S2C |
| GST-Atg8 | *pGEX-4T3-Atg8* | 2A-B, S2D |
| GST-Atg8 Y49A, L50A | *pGEX-4T3-Atg8 (Y49A, L50A)* | 2A |
| GST-Ede1 AIM1 | *pGEX-4T3-Ede1 (F1226A, V1229A)* | 2C |
| GST-Ede1 AIM2 | *pGEX-4T3-Ede1 (F1238A, I1241A)* | 2C |
| GST-Ede1 AIM3 | *pGEX-4T3-Ede1 (W1288A, I1291A)* | 2C |
| GST-Ede1 AIM1+2 | *pGEX-4T3-Ede1 (F1226A, V1229A, F1238A, I1241A)* | S2C |
| GST-Ede1 AIM1+3 | *pGEX-4T3-Ede1 (F1226A, V1229A, I1241A, W1288A, I1291A)* | S2C |
| GST-Ede1 AIM2+3 | *pGEX-4T3-Ede1 (F1238A, I1241A, W1288A, I1291A)* | S2C |
| GST-Ede1 AIMR | *pGEX-4T3-Ede1 (F1226A, V1229A, F1238A, I1241A, W1288A, I1291A)* | 2C |
| GST-Ede1 EH 3W-A | *pGEX-4T3-Ede1 (W56A, W176A, W319A)* | 2C |
| RFP-Atg8 | *pRS414-pMet17-2xmCherry-Atg8* | S5C |
| GFP-Atg8 | *yCplac33-yeGFP-Atg8* | 1F, 3H |
| GFP | *yCplac33-yeGFP* | 1F, 3H |
| 2HA-Atg8 | *p415 ADH-2×HA-Atg8* | 6E |
| GST-Atg8 | *pGEX-2TK-GST-Atg8* | 2D, S2E-F |

## References

Alberti, S., Gladfelter, A., and Mittag, T. (2019). Considerations and Challenges in Studying Liquid-Liquid Phase Separation and Biomolecular Condensates. Cell 176, 419–434.

Banani, S.F., Lee, H.O., Hyman, A.A., and Rosen, M.K. (2017). Biomolecular condensates: organizers of cellular biochemistry. Nat Rev Mol Cell Biol 18, 285–298.

Behrends, C., Sowa, M.E., Gygi, S.P., and Harper, J.W. (2010). Network organization of the human autophagy system. Nature 466, 68–76.

Bejarano, E., Girao, H., Yuste, A., Patel, B., Marques, C., Spray, D.C., Pereira, P., and Cuervo, A.M. (2012). Autophagy modulates dynamics of connexins at the plasma membrane in a ubiquitin-dependent manner. Mol Biol Cell 23, 2156–2169.

Bergeron-Sandoval, L.-P., Heris, H.K., Chang, C., Cornell, C.E., Keller, S.L., François, P., Hendricks, A.G., Ehrlicher, A.J., Pappu, R.V., and Michnick, S.W. (2018). Endocytosis caused by liquid-liquid phase separation of proteins. bioRxiv, 145664.

Boeke, D., Trautmann, S., Meurer, M., Wachsmuth, M., Godlee, C., Knop, M., and Kaksonen, M. (2014). Quantification of cytosolic interactions identifies Ede1 oligomers as key organizers of endocytosis. Mol Syst Biol 10, 756.

Boettner, D.R., Chi, R.J., and Lemmon, S.K. (2011). Lessons from yeast for clathrin-mediated endocytosis. Nat Cell Biol 14, 2–10.

Carlsson, S.R., and Simonsen, A. (2015). Membrane dynamics in autophagosome biogenesis. J Cell Sci 128, 193–205.

Carroll, S.Y., Stimpson, H.E., Weinberg, J., Toret, C.P., Sun, Y., and Drubin, D.G. (2012). Analysis of yeast endocytic site formation and maturation through a regulatory transition point. Mol Biol Cell 23, 657–668.

Cheong, H., and Klionsky, D.J. (2008). Biochemical methods to monitor autophagy-related processes in yeast. Methods Enzymol 451, 1–26.

Day, K.J., Kago, G., Wang, L., Richter, J.B., Hayden, C.C., Lafer, E.M., and Stachowiak, J.C. (2019). Liquid-like protein interactions catalyze assembly of endocytic vesicles. bioRxiv, 860684.

De Tito, S., Hervas, J.H., van Vliet, A.R., and Tooze, S.A. (2020). The Golgi as an Assembly Line to the Autophagosome. Trends Biochem Sci.

Dikic, I. (2017). Proteasomal and Autophagic Degradation Systems. Annu Rev Biochem 86, 193–224.

Dikic, I., and Elazar, Z. (2018). Mechanism and medical implications of mammalian autophagy. Nat Rev Mol Cell Biol 19, 349–364.

Dores, M.R., Schnell, J.D., Maldonado-Baez, L., Wendland, B., and Hicke, L. (2010). The function of yeast epsin and Ede1 ubiquitin-binding domains during receptor internalization. Traffic 11, 151–160.

Epple, U.D., Suriapranata, I., Eskelinen, E.L., and Thumm, M. (2001). Aut5/Cvt17p, a putative lipase essential for disintegration of autophagic bodies inside the vacuole. J Bacteriol 183, 5942–5955.

Farre, J.C., and Subramani, S. (2016). Mechanistic insights into selective autophagy pathways: lessons from yeast. Nat Rev Mol Cell Biol 17, 537–552.

Fujioka, Y., Alam, J.M., Noshiro, D., Mouri, K., Ando, T., Okada, Y., May, A.I., Knorr, R.L., Suzuki, K., Ohsumi, Y., et al. (2020). Phase separation organizes the site of autophagosome formation. Nature 578, 301–305.

Fumagalli, F., Noack, J., Bergmann, T.J., Cebollero, E., Pisoni, G.B., Fasana, E., Fregno, I., Galli, C., Loi, M., Solda, T., et al. (2016). Translocon component Sec62 acts in endoplasmic reticulum turnover during stress recovery. Nat Cell Biol 18, 1173–1184.

Gatica, D., Lahiri, V., and Klionsky, D.J. (2018). Cargo recognition and degradation by selective autophagy. Nat Cell Biol 20, 233–242.

Goode, B.L., Eskin, J.A., and Wendland, B. (2015). Actin and endocytosis in budding yeast. Genetics 199, 315–358.

Harmon, T.S., Holehouse, A.S., Rosen, M.K., and Pappu, R.V. (2017). Intrinsically disordered linkers determine the interplay between phase separation and gelation in multivalent proteins. Elife 6.

Ho, K.H., Chang, H.E., and Huang, W.P. (2009). Mutation at the cargo-receptor binding site of Atg8 also affects its general autophagy regulation function. Autophagy 5, 461–471.

Hurley, J.H., and Young, L.N. (2017). Mechanisms of Autophagy Initiation. Annu Rev Biochem 86, 225–244.

Idrissi, F.Z., Grotsch, H., Fernandez-Golbano, I.M., Presciatto-Baschong, C., Riezman, H., and Geli, M.I. (2008). Distinct acto/myosin-I structures associate with endocytic profiles at the plasma membrane. J Cell Biol 180, 1219–1232.

Inoue, Y., Taguchi, H., Kishimoto, A., and Yoshida, M. (2004). Hsp104 binds to yeast Sup35 prion fiber but needs other factor(s) to sever it. J Biol Chem 279, 52319–52323.

Johansen, T., and Lamark, T. (2020). Selective Autophagy: ATG8 Family Proteins, LIR Motifs and Cargo Receptors. J Mol Biol 432, 80–103.

Kalvari, I., Tsompanis, S., Mulakkal, N.C., Osgood, R., Johansen, T., Nezis, I.P., and Promponas, V.J. (2014). iLIR: A web resource for prediction of Atg8-family interacting proteins. Autophagy 10, 913–925.

Kerppola, T.K. (2008). Bimolecular fluorescence complementation (BiFC) analysis as a probe of protein interactions in living cells. Annu Rev Biophys 37, 465–487.

Kirchhausen, T., Owen, D., and Harrison, S.C. (2014). Molecular structure, function, and dynamics of clathrin-mediated membrane traffic. Cold Spring Harb Perspect Biol 6, a016725.

Kirkin, V. (2020). History of the Selective Autophagy Research: How Did It Begin and Where Does It Stand Today? J Mol Biol 432, 3–27.

Kirkin, V., and Rogov, V.V. (2019). A Diversity of Selective Autophagy Receptors Determines the Specificity of the Autophagy Pathway. Mol Cell 76, 268–285.

Kozak, M., and Kaksonen, M. (2019). Phase separation of Ede1 promotes the initiation of endocytic events. bioRxiv, 861203.

Kukulski, W., Schorb, M., Kaksonen, M., and Briggs, J.A. (2012). Plasma membrane reshaping during endocytosis is revealed by time-resolved electron tomography. Cell 150, 508–520.

Lee, C.W., Wilfling, F., Ronchi, P., Allegretti, M., Mosalaganti, S., Jentsch, S., Beck, M., and Pfander, B. (2020a). Selective autophagy degrades nuclear pore complexes. Nat Cell Biol 22, 159–166.

Lee, J.E., Cathey, P.I., Wu, H., Parker, R., and Voeltz, G.K. (2020b). Endoplasmic reticulum contact sites regulate the dynamics of membraneless organelles. Science 367.

Li, J., Zhu, R., Chen, K., Zheng, H., Zhao, H., Yuan, C., Zhang, H., Wang, C., and Zhang, M. (2018). Potent and specific Atg8-targeting autophagy inhibitory peptides from giant ankyrins. Nat Chem Biol 14, 778–787.

Lu, R., and Drubin, D.G. (2017). Selection and stabilization of endocytic sites by Ede1, a yeast functional homologue of human Eps15. Mol Biol Cell 28, 567–575.

Lu, R., Drubin, D.G., and Sun, Y. (2016). Clathrin-mediated endocytosis in budding yeast at a glance. J Cell Sci 129, 1531–1536.

Lynch-Day, M.A., and Klionsky, D.J. (2010). The Cvt pathway as a model for selective autophagy. FEBS Lett 584, 1359–1366.

Maeda, S., Otomo, C., and Otomo, T. (2019). The autophagic membrane tether ATG2A transfers lipids between membranes. Elife 8.

Martin, E.W., and Mittag, T. (2018). Relationship of Sequence and Phase Separation in Protein Low-Complexity Regions. Biochemistry 57, 2478–2487.

Matscheko, N., Mayrhofer, P., Rao, Y., Beier, V., and Wollert, T. (2019). Atg11 tethers Atg9 vesicles to initiate selective autophagy. PLoS Biol 17, e3000377.

McMahon, H.T., and Boucrot, E. (2011). Molecular mechanism and physiological functions of clathrin-mediated endocytosis. Nat Rev Mol Cell Biol 12, 517–533.

Merrifield, C.J., and Kaksonen, M. (2014). Endocytic accessory factors and regulation of clathrin-mediated endocytosis. Cold Spring Harb Perspect Biol 6, a016733.

Mettlen, M., Chen, P.H., Srinivasan, S., Danuser, G., and Schmid, S.L. (2018). Regulation of Clathrin-Mediated Endocytosis. Annu Rev Biochem 87, 871–896.

Meyerholz, A., Hinrichsen, L., Groos, S., Esk, P.C., Brandes, G., and Ungewickell, E.J. (2005). Effect of clathrin assembly lymphoid myeloid leukemia protein depletion on clathrin coat formation. Traffic 6, 1225–1234.

Mizushima, N., Yoshimori, T., and Ohsumi, Y. (2011). The role of Atg proteins in autophagosome formation. Annu Rev Cell Dev Biol 27, 107–132.

Molliex, A., Temirov, J., Lee, J., Coughlin, M., Kanagaraj, A.P., Kim, H.J., Mittag, T., and Taylor, J.P. (2015). Phase separation by low complexity domains promotes stress granule assembly and drives pathological fibrillization. Cell 163, 123–133.

Monahan, Z., Ryan, V.H., Janke, A.M., Burke, K.A., Rhoads, S.N., Zerze, G.H., O’Meally, R., Dignon, G.L., Conicella, A.E., Zheng, W., et al. (2017). Phosphorylation of the FUS low-complexity domain disrupts phase separation, aggregation, and toxicity. EMBO J 36, 2951–2967.

Mulholland, J., Preuss, D., Moon, A., Wong, A., Drubin, D., and Botstein, D. (1994). Ultrastructure of the yeast actin cytoskeleton and its association with the plasma membrane. J Cell Biol 125, 381–391.

Nakatogawa, H., Suzuki, K., Kamada, Y., and Ohsumi, Y. (2009). Dynamics and diversity in autophagy mechanisms: lessons from yeast. Nat Rev Mol Cell Biol 10, 458–467.

Noda, N.N., and Inagaki, F. (2015). Mechanisms of Autophagy. Annu Rev Biophys 44, 101–122.

Noda, N.N., Ohsumi, Y., and Inagaki, F. (2010). Atg8-family interacting motif crucial for selective autophagy. FEBS Lett 584, 1379–1385.

Noda, T., and Ohsumi, Y. (1998). Tor, a phosphatidylinositol kinase homologue, controls autophagy in yeast. J Biol Chem 273, 3963–3966.

Ohsumi, Y. (2001). Molecular dissection of autophagy: two ubiquitin-like systems. Nat Rev Mol Cell Biol 2, 211–216.

Osawa, T., Kotani, T., Kawaoka, T., Hirata, E., Suzuki, K., Nakatogawa, H., Ohsumi, Y., and Noda, N.N. (2019). Atg2 mediates direct lipid transfer between membranes for autophagosome formation. Nat Struct Mol Biol 26, 281–288.

Patel, A., Lee, H.O., Jawerth, L., Maharana, S., Jahnel, M., Hein, M.Y., Stoynov, S., Mahamid, J., Saha, S., Franzmann, T.M., et al. (2015). A Liquid-to-Solid Phase Transition of the ALS Protein FUS Accelerated by Disease Mutation. Cell 162, 1066–1077.

Popelka, H., and Klionsky, D.J. (2015). Analysis of the native conformation of the LIR/AIM motif in the Atg8/LC3/GABARAP-binding proteins. Autophagy 11, 2153–2159.

Reggiori, F., Komatsu, M., Finley, K., and Simonsen, A. (2012). Autophagy: more than a nonselective pathway. Int J Cell Biol 2012, 219625.

Schutter, M., Giavalisco, P., Brodesser, S., and Graef, M. (2020). Local Fatty Acid Channeling into Phospholipid Synthesis Drives Phagophore Expansion during Autophagy. Cell 180, 135–149 e114.

Shibutani, S.T., and Yoshimori, T. (2014). A current perspective of autophagosome biogenesis. Cell Res 24, 58–68.

Shintani, T., and Klionsky, D.J. (2004). Cargo proteins facilitate the formation of transport vesicles in the cytoplasm to vacuole targeting pathway. J Biol Chem 279, 29889–29894.

Shorter, J., and Lindquist, S. (2004). Hsp104 catalyzes formation and elimination of self-replicating Sup35 prion conformers. Science 304, 1793–1797.

Smith, S.M., Baker, M., Halebian, M., and Smith, C.J. (2017). Weak Molecular Interactions in Clathrin-Mediated Endocytosis. Front Mol Biosci 4, 72.

Sun, D., Wu, R., Zheng, J., Li, P., and Yu, L. (2018). Polyubiquitin chain-induced p62 phase separation drives autophagic cargo segregation. Cell Res 28, 405–415.

Sun, Y., Leong, N.T., Jiang, T., Tangara, A., Darzacq, X., and Drubin, D.G. (2017). Switch-like Arp2/3 activation upon WASP and WIP recruitment to an apparent threshold level by multivalent linker proteins in vivo. Elife 6.

Sun, Y., Leong, N.T., Wong, T., and Drubin, D.G. (2015). A Pan1/End3/Sla1 complex links Arp2/3-mediated actin assembly to sites of clathrin-mediated endocytosis. Mol Biol Cell 26, 3841–3856.

Takagi, T., Ishijima, S.A., Ochi, H., and Osumi, M. (2003). Ultrastructure and behavior of actin cytoskeleton during cell wall formation in the fission yeast Schizosaccharomyces pombe. J Electron Microsc (Tokyo) 52, 161–174.

Torggler, R., Papinski, D., Brach, T., Bas, L., Schuschnig, M., Pfaffenwimmer, T., Rohringer, S., Matzhold, T., Schweida, D., Brezovich, A., et al. (2016). Two Independent Pathways within Selective Autophagy Converge to Activate Atg1 Kinase at the Vacuole. Mol Cell 64, 221–235.

Valverde, D.P., Yu, S., Boggavarapu, V., Kumar, N., Lees, J.A., Walz, T., Reinisch, K.M., and Melia, T.J. (2019). ATG2 transports lipids to promote autophagosome biogenesis. J Cell Biol 218, 1787–1798.

Wang, Z., and Zhang, H. (2019). Phase Separation, Transition, and Autophagic Degradation of Proteins in Development and Pathogenesis. Trends Cell Biol 29, 417–427.

Weinberg, J., and Drubin, D.G. (2012). Clathrin-mediated endocytosis in budding yeast. Trends Cell Biol 22, 1–13.

Wen, X., and Klionsky, D.J. (2016). An overview of macroautophagy in yeast. J Mol Biol 428, 1681–1699.

Wirth, M., Zhang, W., Razi, M., Nyoni, L., Joshi, D., O’Reilly, N., Johansen, T., Tooze, S.A., and Mouilleron, S. (2019). Molecular determinants regulating selective binding of autophagy adapters and receptors to ATG8 proteins. Nat Commun 10, 2055.

Yamasaki, A., Alam, J.M., Noshiro, D., Hirata, E., Fujioka, Y., Suzuki, K., Ohsumi, Y., and Noda, N.N. (2020). Liquidity Is a Critical Determinant for Selective Autophagy of Protein Condensates. Mol Cell 77, 1163–1175 e1169.

Zaffagnini, G., and Martens, S. (2016). Mechanisms of Selective Autophagy. J Mol Biol 428, 1714–1724.

Zambrano, R., Conchillo-Sole, O., Iglesias, V., Illa, R., Rousseau, F., Schymkowitz, J., Sabate, R., Daura, X., and Ventura, S. (2015). PrionW: a server to identify proteins containing glutamine/asparagine rich prion-like domains and their amyloid cores. Nucleic Acids Res 43, W331–337.

## Supplement References

Arnold, J., Mahamid, J., Lucic, V., de Marco, A., Fernandez, J.J., Laugks, T., Mayer, T., Hyman, A.A., Baumeister, W., and Plitzko, J.M. (2016). Site-Specific Cryo-focused Ion Beam Sample Preparation Guided by 3D Correlative Microscopy. Biophys J 110, 860–869.

Collins, S.R., Roguev, A., and Krogan, N.J. (2010). Quantitative genetic interaction mapping using the E-MAP approach. Methods Enzymol 470, 205–231.

Cox, J., Hein, M.Y., Luber, C.A., Paron, I., Nagaraj, N., and Mann, M. (2014). Accurate proteome-wide label-free quantification by delayed normalization and maximal peptide ratio extraction, termed MaxLFQ. Mol Cell Proteomics 13, 2513–2526.

Cox, J., and Mann, M. (2008). MaxQuant enables high peptide identification rates, individualized p.p.b.-range mass accuracies and proteome-wide protein quantification. Nat Biotechnol 26, 1367–1372.

Dunham, M.J., Dunham, M.J., Gartenberg, M.R., and Brown, G.W. (2015). Methods in yeast genetics and genomics : a Cold Spring Harbor Laboratory course manual / Maitreya J. Dunham, University of Washington, Marc R. Gartenberg, Robert Wood Johnson Medical School, Rutgers, The State University of New Jersey, Grant W. Brown, University of Toronto, 2015 edition / edn (Cold Spring Harbor, New York: Cold Spring Harbor Laboratory Press).

Hagen, W.J.H., Wan, W., and Briggs, J.A.G. (2017). Implementation of a cryo-electron tomography tilt-scheme optimized for high resolution subtomogram averaging. J Struct Biol 197, 191–198.

Hrabe, T., Chen, Y., Pfeffer, S., Cuellar, L.K., Mangold, A.V., and Forster, F. (2012). PyTom: a python-based toolbox for localization of macromolecules in cryo-electron tomograms and subtomogram analysis. J Struct Biol 178, 177–188.

Hubner, N.C., Bird, A.W., Cox, J., Splettstoesser, B., Bandilla, P., Poser, I., Hyman, A., and Mann, M. (2010). Quantitative proteomics combined with BAC TransgeneOmics reveals in vivo protein interactions. J Cell Biol 189, 739–754.

Knop, M., Siegers, K., Pereira, G., Zachariae, W., Winsor, B., Nasmyth, K., and Schiebel, E. (1999). Epitope tagging of yeast genes using a PCR-based strategy: more tags and improved practical routines. Yeast 15, 963–972.

Kremer, J.R., Mastronarde, D.N., and McIntosh, J.R. (1996). Computer visualization of three-dimensional image data using IMOD. J Struct Biol 116, 71–76.

Lu, A.X., Zarin, T., Hsu, I.S., and Moses, A.M. (2019). YeastSpotter: accurate and parameter-free web segmentation for microscopy images of yeast cells. Bioinformatics 35, 4525–4527.

Mastronarde, D.N. (2005). Automated electron microscope tomography using robust prediction of specimen movements. J Struct Biol 152, 36–51.

Noda, T., and Klionsky, D.J. (2008). The quantitative Pho8Delta60 assay of nonspecific autophagy. Methods Enzymol 451, 33–42.

Perez-Riverol, Y., Csordas, A., Bai, J., Bernal-Llinares, M., Hewapathirana, S., Kundu, D.J., Inuganti, A., Griss, J., Mayer, G., Eisenacher, M., et al. (2019). The PRIDE database and related tools and resources in 2019: improving support for quantification data. Nucleic Acids Res 47, D442–D450.

Pettersen, E.F., Goddard, T.D., Huang, C.C., Couch, G.S., Greenblatt, D.M., Meng, E.C., and Ferrin, T.E. (2004). UCSF Chimera--a visualization system for exploratory research and analysis. J Comput Chem 25, 1605–1612.

Phair, R.D., Gorski, S.A., and Misteli, T. (2004). Measurement of dynamic protein binding to chromatin in vivo, using photobleaching microscopy. Methods Enzymol 375, 393–414.

Rappsilber, J., Ishihama, Y., and Mann, M. (2003). Stop and go extraction tips for matrix-assisted laser desorption/ionization, nanoelectrospray, and LC/MS sample pretreatment in proteomics. Anal Chem 75, 663–670.

Rigort, A., Bauerlein, F.J., Leis, A., Gruska, M., Hoffmann, C., Laugks, T., Bohm, U., Eibauer, M., Gnaegi, H., Baumeister, W., et al. (2010). Micromachining tools and correlative approaches for cellular cryo-electron tomography. J Struct Biol 172, 169–179.

Rohou, A., and Grigorieff, N. (2015). CTFFIND4: Fast and accurate defocus estimation from electron micrographs. J Struct Biol 192, 216–221.

Schaffer, M., Mahamid, J., Engel, B.D., Laugks, T., Baumeister, W., and Plitzko, J.M. (2017). Optimized cryo-focused ion beam sample preparation aimed at in situ structural studies of membrane proteins. J Struct Biol 197, 73–82.

Scheres, S.H. (2012). RELION: implementation of a Bayesian approach to cryo-EM structure determination. J Struct Biol 180, 519–530.

Schindelin, J., Arganda-Carreras, I., Frise, E., Kaynig, V., Longair, M., Pietzsch, T., Preibisch, S., Rueden, C., Saalfeld, S., Schmid, B., et al. (2012). Fiji: an open-source platform for biological-image analysis. Nat Methods 9, 676–682.

Schneider, C.A., Rasband, W.S., and Eliceiri, K.W. (2012). NIH Image to ImageJ: 25 years of image analysis. Nat Methods 9, 671–675.

Sherman, F. (2002). Getting started with yeast. Methods Enzymol 350, 3–41.

Snapp, E.L. (2013). Photobleaching methods to study Golgi complex dynamics in living cells. Methods Cell Biol 118, 195–216.

Tal, R., Winter, G., Ecker, N., Klionsky, D.J., and Abeliovich, H. (2007). Aup1p, a yeast mitochondrial protein phosphatase homolog, is required for efficient stationary phase mitophagy and cell survival. J Biol Chem 282, 5617–5624.

Tyanova, S., Temu, T., Sinitcyn, P., Carlson, A., Hein, M.Y., Geiger, T., Mann, M., and Cox, J. (2016). The Perseus computational platform for comprehensive analysis of (prote)omics data. Nat Methods 13, 731–740.

Zheng, S.Q., Palovcak, E., Armache, J.P., Verba, K.A., Cheng, Y., and Agard, D.A. (2017a). MotionCor2: anisotropic correction of beam-induced motion for improved cryo-electron microscopy. Nat Methods 14, 331–332.

Zheng, Y., Qiu, Y., Gunderson, J.E., and Schulman, B.A. (2017b). Production of Human ATG Proteins for Lipidation Assays. Methods Enzymol 587, 97–113.

